# EpiATLAS – a reference for human epigenomic research

**DOI:** 10.64898/2026.06.22.729579

**Authors:** International Human Epigenome Consortium, Quirin Manz, Misha Bilenky, Dennis Hecker, Nihit Aggarwal, Juliana E Arcila-Galvis, Shamim Ashrafiyan, Nina Baumgarten, Fatemeh Behjati Ardakani, Paulo R Branco Lins, Charles E Breeze, David Brownlee, David Bujold, Alec R Chapman, Savio Ho-Chit Chow, Tevfik Umut Dincer, Charles Dupras, Gabriella Frosi, Jingyuan Fu, Deborah Gérard, Axel Hauduc, Jeffrey Hyacinthe, Artur Jaroszewicz, Runjia Li, Riley J Mangan, Aneta Mikulasova, Ismail Moghul, Maria Needhamsen, Nicole Palmour, Maria Pires Pacheco, Jacob Quon, Joanny Raby, Alex Reynolds, Laura Rumpf, Abdulrahman Salhab, Christina Huan Shi, Lasse Sinkkonen, Yosuke Tanigawa, R Matthew Tanner, Ha Vu, Frédérique White, Jacqueline T M Aw, Shirin Badii, Reanne Bowlby, Merrill Boyle, Carl Brown, Damini Chand, Marc Calingo, Qi Cao, Annaick Carles, Maxime Caron, Marcus Carreira, Alfred Sze Lok Cheng, Clooney Cheuk Yin Cheng, Dean Cheng, Young Cheng, Ming Fung Cheung, Gina Choe, Eric Chuah, Rola Dali, Athena Deng, Sitanshu Gakkhar, Marta Gut, An He, Simon Charles Heath, Vincy Ho, Angelica Grace Intan, Caryn Y Ito, Qinghong Jiang, Joost L Kluiver, Joon Lee, Seohyun Lee, Ziuwin Leung, Irene Li, Alireza Lorzadeh, Dan McKerricher, Neha Mishra, Karen Mungall, Luis Augusto Eijy Nagai, Mzwanele Ngubo, Diana Palmquist, Johnson Pang, Victoria Park, Patrick Plettner, Akshita Puri, Adriana Redensek, Zohreh Sharafian, Marie-Michelle Simon, Jonathan Steif, Edmund Su, Sabrina Ka Man Tam, Sindy Sing Ting Tam, Anke Van den Berg, Rachel Wong, Tina Wong, Jenny Ka Hei Wu, Alice Zhu, Tony Kwan, Michelle Moksa, Sinead T Aherne, Sam Aparicio, Takahiro Arima, Ho Ryun Chung, Joseph F Costello, Connie J Eaves, Susan J Fisher, Ivo Glynne Gut, Peter W Harrison, Bernhard Horsthemke, Garth R Ilsley, Maja Jagodic, Aly Karsan, Pascal M Lavoie, Danny Leung, Maxwell W Libbrecht, Thomas Manke, Marco A Marra, Wouter Meuleman, Fabian Müller, Ryuichiro Nakato, Hiroaki Okae, Daniel Rico, Philip Rosenstiel, Hiroyuki Sasaki, Thomas Sauter, Stefan Schreiber, David W Scott, William L Stanford, Christian Steidl, Hendrik G Stunnenberg, Mikita Suyama, Florian Tran, Toshikazu Ushijima, Andrew P Weng, Sam M Wiseman, Angela Ruohao Wu, Kevin Y Yip, Stephen Yip, Stephan Beck, Guillaume Bourque, Jason Ernst, Pierre-Étienne Jacques, Yann Joly, Manolis Kellis, Markus List, Theodore J Perkins, Marcel H Schulz, Jörn Walter, Martin Hirst

## Abstract

The sequence of the human genome provides a foundation for understanding cellular processes in health and disease^1^. The organisation of this primary genetic information into cell-specific structure and function is critical to understanding the cell type-specific interpretation and execution of the genome. Epigenetic processes are essential for packaging and higher-level functional organisation of the genome, and changes therein are increasingly recognised as contributors to human disease. Building on primary data generated by multinational consortia, the International Human Epigenome Consortium^2^ (IHEC) has uniformly processed a collection of more than 2000 comprehensive human reference epigenomes, collectively referred to as EpiATLAS. This effort involved the development of standardised molecular and bioinformatics protocols, metadata models, and analytical tools to manage, integrate, display, and share vast amounts of epigenomic data. This includes the creation of a publicly available Epigenome Reference Registry, which provides a system for accessing protected human subject datasets and facilitates open searching of de-identified samples and experimental data. The integrated EpiATLAS ecosystem and its comprehensive human reference epigenome maps provide an unprecedented resource for the biosciences, expanding the annotated epigenomic landscape while uncovering previously unappreciated relationships among regulatory layers and revealing how epigenetic inputs underpin fundamental cellular functions and disease associations.

## Main

Mammalian genomes carry non-uniform chemical marks covalently attached to DNA bases or to the chromatin proteins that regulate local activity states, supporting gene activation or silencing. Normal cell development is accompanied by marked changes in the epigenome, and specific epigenome signatures distinguish pluripotent, developing and terminally differentiated cells. Epidemiological and model organism studies have demonstrated that epigenetic modifications can be induced by a variety of modulators, including diverse environmental stimuli, such as stress, nutrient levels, microbiota, and toxin exposure. As epigenetic modifications can be transiently, mitotically and in some cases transgenerationally heritable, they provide a framework in which to investigate how environment and lifestyle can impact disease susceptibility and progression. Epigenetic modifications are also central to chromatin dynamics and play key roles in many biological processes, including DNA replication and repair, transcription and development through molecular interactions with regulatory machinery such as transcription factors. Anomalies in epigenetic programming characterise many human diseases, including cancer, cardiovascular disease, neuropsychiatric disorders, imprinting disorders, inflammation, and autoimmune disease^3^. Epigenome changes can be reversible, and several inhibitors of chromatin-modifying enzymes, including histone deacetylase and DNA methyltransferase inhibitors, have demonstrated clinical activity^3^.

Technological advances now enable reproducible, whole-genome profiling of human epigenomic marks, but the sheer diversity of epigenomes across hundreds of cell types has long limited the ability to quantify and interpret normal and disease-associated variation. To provide a unified framework for epigenomic exploration, the International Human Epigenome Consortium^2^ (IHEC; https://ihec-epigenomes.org/) was established in 2010 with a goal of producing 1000 reference epigenome maps for human cell types and states relevant to health and disease. IHEC integrates major international initiatives, including AMED CREST/IHEC Team Japan, the German Epigenome Programme (DEEP), the Canadian Epigenetics, Environment and Health Research Consortium (CEEHRC), the EU FP7 BLUEPRINT Project, the Hong Kong Epigenomics Project (EpiHK), the Korea Epigenome Project (KNIH), NHGRI ENCODE, the NIH Roadmap Epigenomics Program, and the Singapore Epigenome Project (GIS). Together, these efforts have generated more than 2000 high-quality reference epigenomes spanning DNA methylation, histone modifications, and coding and non-coding RNA expression. These datasets were systematically reprocessed using harmonised reference resources and standardised analytical pipelines to generate the EpiATLAS, a uniformly processed, interoperable epigenomic resource that enables robust comparative analyses across cell types, tissues, and studies. Together, the IHEC reference epigenomes, analytical standards, and ethical framework provide the foundation needed to chart the spectrum of human epigenomic variation and to accelerate the discovery of epigenetic mechanisms, biomarkers, and therapeutic opportunities. Highlights of our contributions and findings are given below.

- Guided by a consortium-developed ethical framework, the epigenomic atlas (EpiATLAS) provides the largest harmonised collection of multimodal human epigenomic data to date, integrating histone modifications, DNA methylation and RNA expression. The resource is enriched with standardised metadata and multiple complementary data representations generated using established and novel computational methods. This resource expands epigenomic annotation into previously uncharacterized genomic regions and strengthens the interpretability of public databases.
- Comparative analyses across thousands of biospecimens reveal distinct, cell- and tissue-specific epigenomic topographies, characteristic histone modification and DNA methylation profiles, and inter-individual variation.
- Whole-genome epigenomic measurements converge on a constrained set of constitutive and cell and tissue type–specific regulatory states. However, substantial dispersion in the genomic occupancy and activity of these states is observed within and across cell types, revealing extensive variability in regulatory marking.
- Integration of chromatin states, enhancers, DNA methylation, and transcription uncovers organizing principles of gene regulation: genes with stable regulatory complexity are enriched for core cellular functions and disease association.

### Reference epigenome mapping

The generation of a comprehensive, international reference of human epigenomes required the development of a robust ethical and governance framework to enable responsible data sharing. To address this challenge, IHEC established the first multi-disciplinary expert working group dedicated to Ethical, Legal and Social Issues (ELSI) in epigenetic research. A central focus of the ELSI group was the development of policies and infrastructure (**Fig. 1a**) that would allow controlled-access human epigenomic data to be discoverable, interoperable, and reusable while respecting participant consent and local regulatory constraints ^4–11^. This effort resulted in a standardised data-sharing framework and the creation of the Epigenomic Reference Registry (EpiRR; www.ebi.ac.uk/epirr), a metadata registry designed to expose rich, harmonised metadata for protected human datasets. EpiRR enables searching and discovery of epigenomic data housed across major international repositories, including the European Genome–phenome Archive (EGA), the DNA Data Bank of Japan (DDBJ) and the database Genotypes and Phenotypes (dbGAP), without requiring direct access to individual-level data. Together, these components provide the ethical and technical foundation upon which EpiATLAS was constructed.

**Figure 1.**
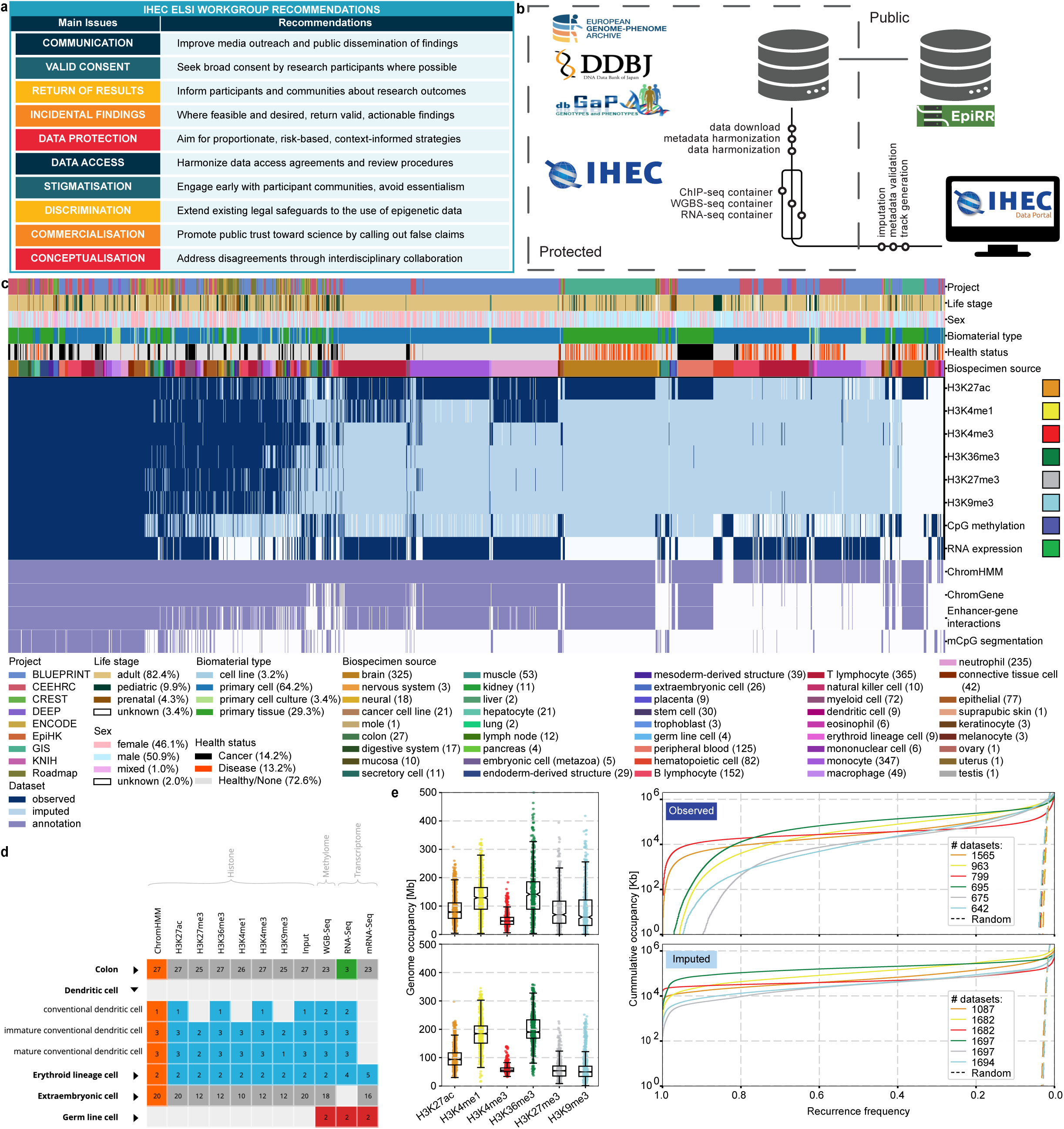
EpiATLAS Overview. **a,** Overview of the ethical considerations of epigenomic research considered by IHEC. **b,** EpiATLAS dataflow and sharing structure enabling public access. **c,** EpiATLAS data matrix showing experimentally observed primary datasets (dark blue), derived imputed datasets (light blue) and integrated state annotations (purple). Selected dataset annotations are shown in the top rows and described in the accompanying legends, with numbers in parentheses referring to observed data only. **d,** The IHEC Data Portal (https://epigenomesportal.ca) provides user-friendly tools to navigate the IHEC data compendium. **e,** Overview of observed and imputed histone modification occupancy. Left: Distribution of genomic occupancy per epigenome of enriched regions of observed (upper) and imputed (lower) datasets by histone modification; Right: Cumulative autosome occupancy distribution as a function of recurrence frequency by histone modification for observed and imputed data. Box plots show centre line, median; box limits, upper and lower quartiles; whiskers, 1.5× interquartile range; points, individual datasets.

To construct EpiATLAS, 7539 genome-wide datasets were generated by 10 IHEC production groups, comprising 5339 histone ChIP–seq datasets (H3K27ac, H3K4me1, H3K4me3, H3K36me3, H3K27me3 and H3K9me3), 1555 RNA-seq datasets, and 645 whole-genome bisulfite sequencing (WGBS) datasets (**Supplementary Table 1**). Datasets were downloaded from protected repositories, including the EGA, DDJB, and dbGAP, and uniformly processed from raw sequence data (.fastq files) using standardised analytical pipelines, yielding a total of 1.4 trillion mapped reads (**Fig. 1b, Supplementary Fig. 1a-c**). To enrich the IHEC EpiATLAS resource, we applied ChromImpute^12^ to impute missing epigenomic datasets. For the 1698 reference epigenomes containing at least one ChIP–seq experiment, we imputed 9539 histone modification and 1698 DNA methylation maps (**Fig. 1c**). In addition, for 449 epigenomes with gene expression data but lacking ChIP–seq measurements, we used the recently developed Gene Expression-based Chromatin State Imputation^13^ (GECSI) method to impute an additional set of chromatin state annotations. To facilitate community access, processed datasets are provided along with standardised quality-control metrics and genome browser tracks, available for both individual and bulk download (**Fig. 1d** and https://ihec-epigenomes.org/).

To assess the extent to which the EpiATLAS observed and imputed ChIP–seq compendium approaches asymptotic genomic occupancy beyond which additional datasets do not substantially increase the genomic coverage of histone modifications, we examined the cross-epigenome recurrence of peaks for each histone modification. Recurrence patterns differ markedly by histone modification (**Fig. 1e**). H3K4me3 displays the highest recurrence across epigenomes, whereas the repressive modifications H3K27me3 and H3K9me3 exhibit substantially lower recurrence across the compendium.

Notably, the recurrence derived from the observed datasets shows substantial dispersion, both within and across purified cell types (**Extended Data Fig. 1a,b, Supplementary Fig. 1d–h**). Rather than reflecting only technical variability, this heterogeneity is also consistent with an emerging view of epigenomic regulation in which cells adopt persistent, context-dependent chromatin states shaped by environmental exposure, tissue microenvironment, and physiological history^14^. Accordingly, epigenomic profiles derived from canonically defined primary cell and tissue types across individuals are expected to capture this accumulated history, manifesting as structured variation that reflects both conserved cell identity and individual-specific environmental and physiological influences. In contrast, recurrence curves from the imputed datasets are smoother and more consistent, providing a standardised, model-smoothed complement to the empirical measurements.

For both observed and imputed data, RNA-seq similarly reveals a core set of recurrently expressed genes (**Extended Data Fig. 1c-e)**. DNA methylation recurrence is highly asymmetric, with hypermethylated CpGs broadly shared across epigenomes and hypomethylated CpGs remaining largely epigenome-specific, further reflecting the interplay between stable regulatory architecture and context-dependent variation (**Extended Data Fig. 1f and g**).

### Integrative epigenome annotation

To complement the IHEC EpiATLAS resource, we generated a series of integrative epigenomic annotations that complement the individual tracks (**Fig. 2, Extended Data Fig. 2**). For each of the 1698 reference epigenomes for which we had at least one ChIP-seq experiment, we generated ChromHMM chromatin state annotations on a per epigenome basis^15^. Specifically, we used a ChromHMM model with 18 states based on six histone modifications, the same model first used in the context of the Roadmap Epigenomics project^16^, to annotate chromatin states at a 200 bp resolution in these IHEC epigenomes. To generate the chromatin state annotations for a reference epigenome, we used observed data for a histone modification where available, and otherwise we used imputed data. We visualised these chromatin state annotations with Epilogos^17^ (**Fig. 2**), which we have made available as an interactive resource (https://ihec-epigenomes.org/epilogos). To complement the individual reference chromatin state annotations, we generated summary ChromHMM annotations for 56 epigenome groups defined using tissue/cell ontology and disease state using CSREP^18^. We also generated universal chromatin state annotations using a stacked ChromHMM model^19^, integrating 5339 IHEC ChIP–seq experiments across 1698 reference epigenomes into a single, genome-wide annotation.

**Figure 2.**
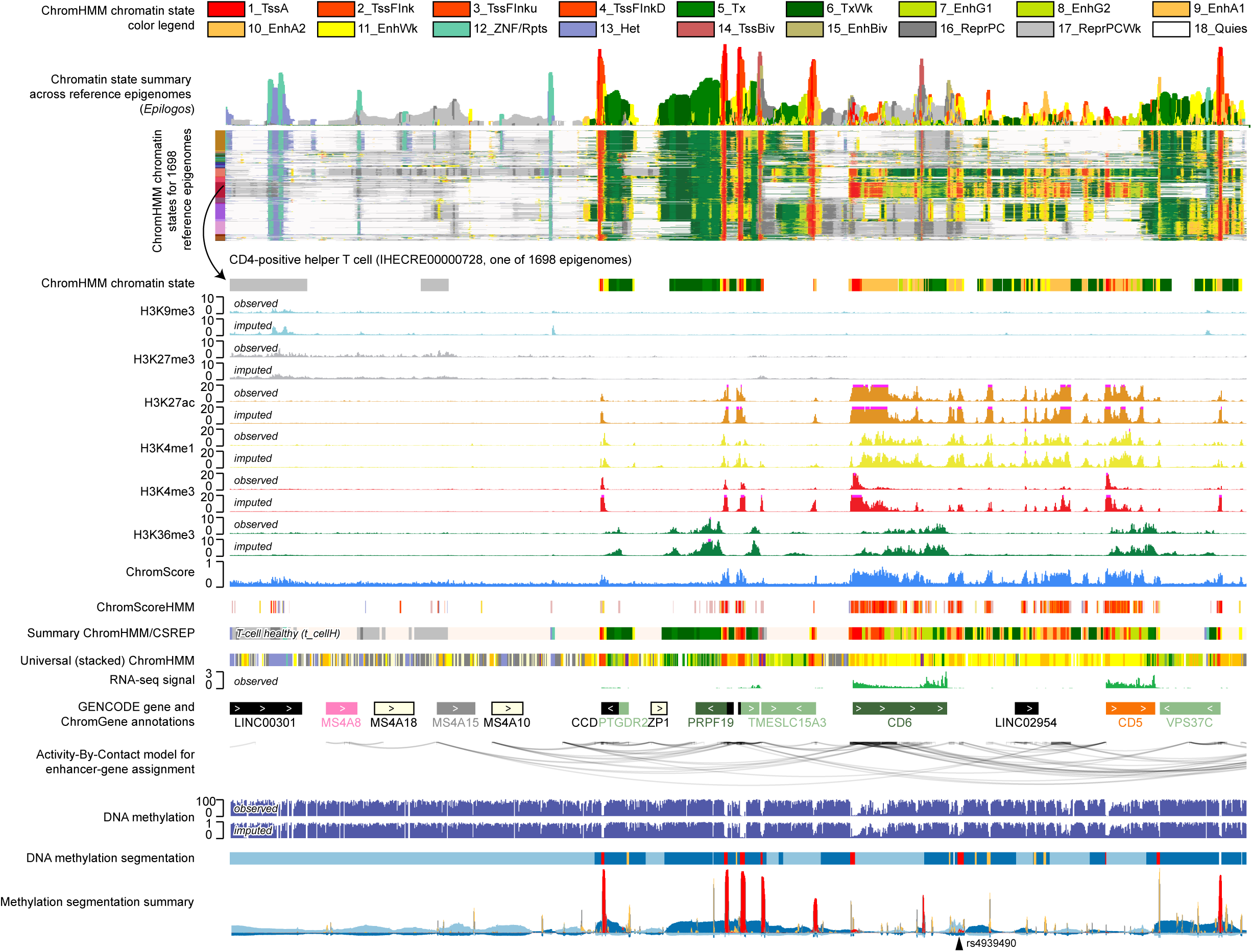
Overview of epigenomic data and integrative annotations. The coordinates of the genomic region in hg38 are chr11:60,650,000-61,175,000. From top to bottom: (1) Colour legend for ChromHMM^15^ chromatin state model for the 18-state model that was previously learned^16^ and applied here. (2) Epilogos visualization^17^ of the 18-state ChromHMM annotations for 1698 reference epigenomes. The upper facet is the Epilogos information-theoretic visualisation summary, followed by the individual epigenome chromatin state annotations. On the left is a colour bar corresponding to the biospecimen groups as in Fig 1c. (3) ChromHMM chromatin state for a CD4-positive helper T cell reference epigenome (IHECRE00000728). (4) The observed followed by the ChromImpute^22^ imputed signal tracks from this reference epigenome for H3K9me3, H3K27me3, H3K27ac, H3K4me1, H3K4me3, H3K36me3. (5) The ChromScore and ChromScoreHMM tracks for epigenome IHECRE00000728. Colour legend for ChromScoreHMM is available in **Extended Data Fig. 2**. (6) A summary ChromHMM track of state assignments based on the 18-state model for epigenomes of the t-cell healthy group, which is one of 56 summary tracks corresponding to combinations of tissue/cell ontology and disease state. (7) A universal chromatin state annotation based on the stacked modelling approach^19^ applied jointly to all EpiATLAS ChIP-seq data. The emission parameters and colour legend are shown in Fig. 4a. (8) Observed RNA-seq signal track for epigenome IHECRE00000728. (9) GENCODE gene annotations with protein coding genes colored by their ChromGene annotation^21^ in epigenome IHECRE00000728 (other genes colored black). ChromGene emission parameters and colour legend can be found in **Supplementary Fig. 2a**. (10) An activity-by-contact model enhancer gene assignments for epigenome IHECRE00000728 from the gABC framework^23^described in Fig. 6. (11) Observed and ChromImpute imputed DNA methylation signal tracks. (12) Four-state methylation segmentation track from MethylSeekR^24^ for epigenome IHECRE00000728. Corresponding colour legend can be found in Fig. 3b. (13) A summary track of methylation segmentations for 505 epigenomes, similar to the Epilogos summary track for chromatin states. (14) Genome-wide association study variant associated with multiple sclerosis (rs4939490) related to analyses in Fig 8.

To annotate genomic positions with predicted regulatory activity, we applied the ChromActivity framework^20^ to generate both ChromScoreHMM and ChromScore annotations. ChromActivity trains multiple supervised predictors based on high-throughput functional assays with various epigenomic features to infer regulatory activity at 25-bp resolution (Methods). ChromActivity then subsequently summarises the outputs from the individual predictors with state-based ChromScoreHMM annotations (**Extended Data Fig. 2**), as well as a single continuous summary metric, ChromScore.

To extend these position-based annotations, we generated gene-level epigenomic annotations using ChromGene^21^. ChromGene models the combinatorial and spatial organisation of epigenomic marks across gene bodies and flanking regions using a mixture of hidden Markov models, and the specific model we used here was composed of 12 mixture components, each with two states (**Supplementary Fig. 2a-c, Supplementary Table 2**). For each reference epigenome, ChromGene assigned each protein-coding gene to the mixture component with the highest posterior probability, providing a gene-centric resource for downstream analyses.

### DNA methylation variance within and across cell types

CpG methylation forms cell-specific patterns linked to gene regulation, chromatin organisation, and genome stability. The high-quality WGBS datasets collected in EpiATLAS (>10x coverage) provide the largest genome-wide methylome datasets across primary human cells and tissues to date. While the majority of cell type-specific methylomes exhibit an overall high methylation, the average CpG methylation across all (non-cancerous) EpiATLAS biosamples varies substantially, ranging from 46% in an extraembryonic tissue biospecimen to 82% in monocytes and brain tissue biospecimens, marking the extremes (**Fig. 3a**).

**Figure 3.**
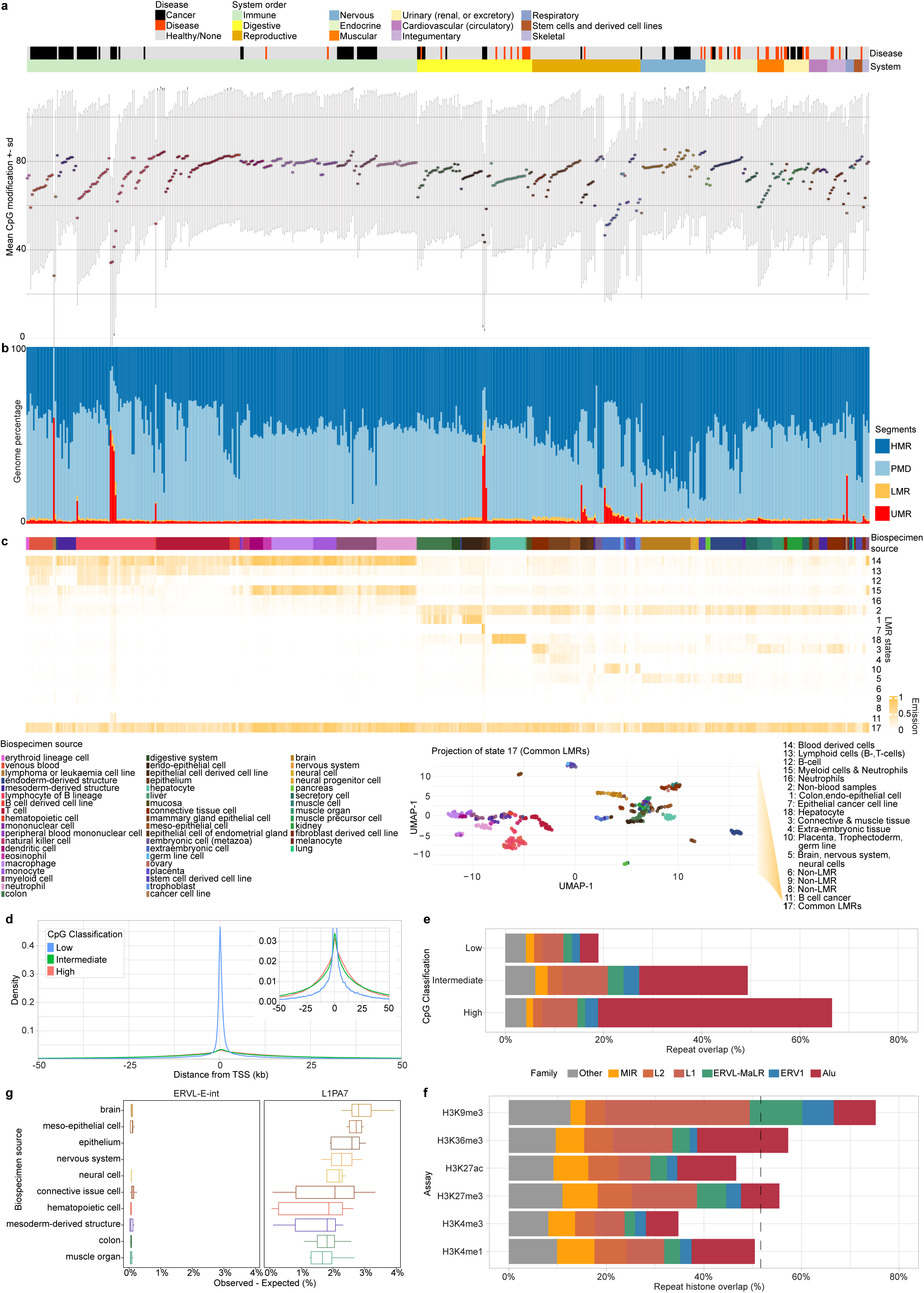
Characteristics of the lowly methylated and repressed regions across cell types: **a,** Average CpG modification with standard deviation across all measured CpGs per sample ordered by system, biospecimen source, and increasing mean methylation. **b,** Genome coverage percentage of methylation states defined by MethylSeekR^28^. We note that, with standard parameters provided by MethylSeekR segmentation, the model generates a high proportional calling of UMRs/LMRs in samples with a low average genome-wide methylation (< 60%), as seen for many cancer and some extra-embryonic samples. This effect has been discussed elsewhere^29^. (HMR: Highly methylated regions; dark blue, PMD: Partially methylated domains; light blue, LMR: Low methylated regions; orange, UMR: Unmethylated region; red). **c,** Cell type-specific LMRs. Top panel: Representation of the emission probabilities calculated meta-segmentation of LMRs using an 18-state model. Bottom panel: UMAP projection of LMRs average methylation based on state-17 (= shared LMRs). **d,** Distance of highly (>95%), intermediate (in 25-95%) and lowly (<25%) methylated CpGs sites across biospecimens to gene transcription start sites (TSS) (all genes, signed distance with respect to TSS direction). **e,** Classification of CpG sites based on methylation prevalence across samples and their overlap with repeats. Repeat families adding up to less than 2% were grouped as Other. **f,** Summary of the repeat overlap of samples across histone modifications. Percentage of peaks of each histone modification that overlap repeats, averaged across samples. The dotted line is the global mean. **g,** Example of a repeat subfamily significantly enriched in cell and tissue types. Enrichment (observed overlap - expected overlap in percent) of two repeat subfamilies (ERVL-E-int, L1PA7) across cell types within H3K9me3 peaks. Top 10 most enriched (median) cell types for L1PA7 shown. Box plots show centre line, median; box limits, upper and lower quartiles; whiskers, 1.5× interquartile range.

To investigate how these genome-wide differences relate to the distribution of CpG methylation across biospecimens, we applied a standard four-state HMM segmentation model (MethylSeekR^16^) for all 505 high-quality samples, partitioning individual methylomes into highly methylated regions (HMRs), partially methylated domains (PMDs), low-methylated regions (LMRs), and unmethylated regions (UMRs). We observe that while HMRs and PMDs together account for a large proportion (up to 96%) of the genome, their relative proportions vary substantially across the 505 individual samples, distinguishing cell and tissue types^25,26^. For a small subset of 24 strongly hypomethylated biospecimens (with <60% average genome-wide methylation), including selected primary cancer specimens, immortalised cancer cell lines and extraembryonic tissues, the four-state MethylSeekR segmentation model strongly increases the calling of UMRs and LMRs (**Fig. 3b**; highlighted by red (UMRs) and yellow (LMRs) spikes), reflecting a general genome-wide hypomethylated state. Collectively, these trends are further supported via an interactive browser (https://epi.genetik.uni-saarland.de/epigenome-segmentation/), showing methylation segmentation summaries for tracks across all 505 samples (**Fig. 2**, bottom two tracks).

To gain deeper insight into the relationship between DNA methylation variance and cell type specificity, we analysed LMRs known to overlap distal regulatory elements^24^. A meta-segmentation approach (as previously described^26^) generates clear LMR patterns which seem to be divided into cell type–specific and shared/common LMRs (**Fig. 3c**). We further investigated the common state 17, showing emissions across all 505 biospecimens. A UMAP projection of the 505 individual state 17 LMR emissions reveals a clear separation into cell and tissue types (**Fig. 3c**, bottom). We conclude that LMRs can be subdivided into distinct subtypes that form quantitative and qualitative variation patterns across biosamples, are linked to cell type–specific epigenomic regulation, and are therefore likely to prove informative for cell type classification.

### Repeat elements underlie distinct genomic distributions of CpG methylation and histone modification

To delineate the genomic context of CpG methylation within EpiATLAS, we analysed individual CpG sites across biospecimens and stratified them according to methylation prevalence. This analysis identified 4.1 M highly methylated CpGs (methylated in >95% of biospecimens), 17.1 M intermediately methylated CpGs (25–95%), and 3.7 M lowly methylated CpGs (<25%) (Methods). As expected, lowly methylated CpGs were markedly enriched within 2 kb of transcription start sites (TSS) (**Fig. 3d**), consistent with regulatory-associated hypomethylation. By contrast, highly and intermediately methylated CpGs displayed broadly similar genomic distributions, with only modest enrichment near TSS.

Highly methylated CpGs overlapped repetitive elements more frequently (67%) than intermediately (50%) or lowly methylated CpGs (21%) (**Fig. 3e**). Nearly half (48%) of highly methylated CpGs overlapped Alu elements. Both highly and intermediately methylated CpGs within repeats were depleted near TSSs compared with non-repetitive CpGs (**Supplementary Fig. 3a**), indicating distinct regulatory contexts.

Across the EpiATLAS, more than half of histone modification peaks overlapped annotated repeats (**Fig. 3f**), with the strongest enrichment observed for the canonical heterochromatic mark H3K9me3. Additional repressive and regulatory marks, including H3K27me3, as well as enhancer- and promoter-associated marks, also showed substantial overlap with repetitive elements. Repeat subfamily analysis further revealed pronounced overlap of L1 elements within H3K9me3-marked regions, consistent with their silencing through heterochromatin-mediated repression^27^, whereas Alu elements were increased in H3K36me3. ERVs were found across multiple marks, while MIR and L2 elements were preferentially associated with activating marks, including H3K27ac and H3K4me1, as well as H3K27me3. An L1 subfamily was enriched in H3K9me3, illustrating substantial cell and tissue type–specific variation within repeat families, while an ERVL-E subfamily lacked such enrichment (**Fig. 3g**; **Supplementary Fig. 3b**).

### Constitutive and cell or tissue type–specific chromatin states

The universal chromatin state annotations, which were defined jointly across all observed ChIP-seq datasets, captured both constitutive and cell or tissue type–specific chromatin activity (**Fig. 4a**). Consistent with prior frameworks, we adopted a 100-state model and assigned functional labels based on overlap with established universal chromatin states derived primarily from Roadmap Epigenomics data^16^ (**Extended Data Fig. 3a**).

**Figure 4.**
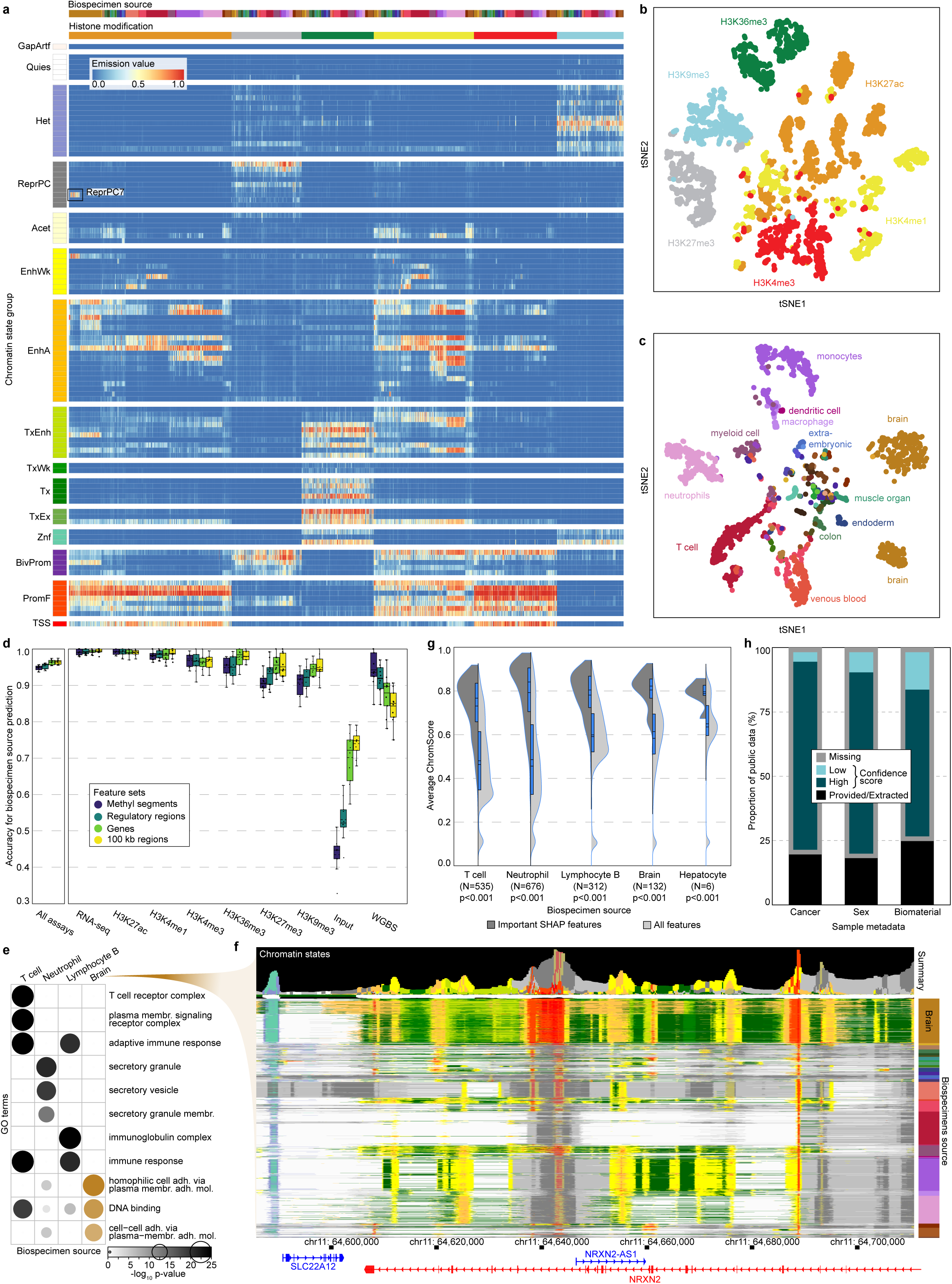
Epigenomic chromatin signatures distinguish cell identity. **a,** Heatmap representation of emission matrix from the EpiATLAS ChromHMM model based on the stacked modeling approach. Rows correspond to states organized into groups. Columns correspond to 5339 datasets, which are organized first by histone modification and within them by biospecimen source (colored as in Fig. 1c). Emission values are indicated by the color scale. **b,** tSNE embedding of histone modification data represented as binary peak presence/absence at promoters and enhancers, colored by histone modification. **c,** tSNE embedding of H3K27ac data, colored by biospecimen source. **d,** Distribution of the EpiClass accuracy per assay over the 10-fold cross-validation training for the classifiers trained on different types of genomic regions (∼300k most variable WGBS 200 bp bins, ∼300k top-correlated regulatory regions (average size ∼2.3 kb), ∼19k genes (average size ∼68 kb) and 30k non-overlapping 100 kb bins covering the whole genome). Box plots show solid centre line, median; dashed centre line, mean; box limits, upper and lower quartiles; whiskers, 1.5× interquartile range; points, individual folds. **e,** Gene Ontology term enrichment using gene names located within the most important regions of the genome according to SHAP analysis from the EpiClass Biospecimen classifier using 100 kb bins (535 important regions for T cell, 676 for Neutrophil, 312 for Lymphocyte B and 132 for Brain). **f,** Epilogos visualization of one of the important Biospecimen classifier regions encoding the Neurexin-2 (NRXN2) gene, enriched in the Brain GO term from panel e. **g,** Distribution of the average ChromScore over the most important SHAP regions used by EpiClass to predict the biospecimen (dark grey) and over the whole genome (light grey) per biospecimen. The number of important regions are shown in parenthesis, as well as the p-value calculated using a two-sided Welch’s t-test. **h,** Proportion of >350,000 public datasets for which the selected metadata was provided/extracted (black), or predicted using EpiClass with high (blue) or low (cyan) prediction score for those where the metadata was missing (grey).

Several states from this new model corresponded to a single enhancer-associated state from the Roadmap model. This was the case, for example, for both blood- and brain-associated enhancer states, revealing more refined annotations of tissue-specific regulatory regions. Notably, one state overlapping a previously repressive polycomb state showed a strong H3K27ac signal in cerebellar tissues, suggesting region-specific enhancers not well captured in the Roadmap reference set. Systematic enrichment analyses further distinguished states by enrichment for genomic features (**Extended Data Fig. 3b**) and for histone modification composition and metadata categories (**Supplementary Table 3**).

To compare chromatin profiles across the 5339 histone modification datasets, we represented each dataset by the presence or absence of peaks at gene promoters and putative regulatory elements defined by the universal chromatin states (**Supplementary Fig. 4a**). Dimensionality reduction revealed that overall similarity between datasets is driven primarily by histone modification type and secondarily by cell or tissue type (**Fig. 4b**), a pattern supported by clustering and classification analyses (**Extended Data Fig. 3c–d**). This separation was particularly pronounced for H3K27me3, H3K9me3, H3K36me3 and H3K4me3, indicating distinct distributions for these marks, whereas H3K27ac and H3K4me1 showed greater overlap, consistent with their shared roles at active regulatory regions^30^. The dominance of histone modification identity over cell type was robust across analytical choices and was also observed in genome-wide analyses based on signal intensities, as well as in joint analyses incorporating DNA methylation and gene expression data (**Extended Data Fig. 3f–j**).

When analysed separately by histone modification, datasets clustered primarily by cell and tissue type (Fig. 4c, Supplementary Fig. 4b), although substantial variability persisted within these groupings. To specifically assess epigenomic variation within a defined cellular context while minimizing confounding effects from cellular heterogeneity, we focused on breast cell subtypes within EpiATLAS, where primary populations from eight individuals had been purified to greater than 90% purity using cell surface markers. Despite this high degree of enrichment and the distinct functional identities of these cell types, per-epigenome ChromHMM state annotations exhibited extensive variability across most chromatin states both within and between cell types. For example, mean enhancer Jaccard values were 0.15 for comparisons within cell types versus 0.06 between cell types, highlighting the unexpectedly high degree of epigenomic divergence retained even among highly purified primary cell types^31^ (**Extended Data Fig. 3e**). These findings reinforce the concept that primary human reference epigenomes may capture substantial inter-individual and environmental influences superimposed upon core cell type–specific regulatory programs.

To systematically distinguish biospecimen categories and other metadata attributes, we developed EpiClass, a machine-learning framework trained to predict eight key metadata characteristics directly from whole-genome epigenomic signal files^32^. EpiClass was trained using epigenomic signal features for multiple representations of the genome, including highly variable DNA methylation segments, regulatory and genic regions, as well as non-overlapping 100 kb bins spanning the genome^33^. Across the 16 most common biospecimen categories, representing approximately two-thirds of the reference epigenomes, EpiClass achieved a mean prediction accuracy of ∼95%, largely independent of region type (**Fig. 4d**, **left**). Performance was consistent for RNA-seq and active histone modification densities regardless of the genomic regions used (**Fig. 4d**, **right**), with similar prediction score distributions across region types (**Supplementary Fig. 4c**). As expected, prediction accuracy was lower for ChIP–seq input controls (∼75% for 100 kb bins), yet remained sufficient to distinguish biospecimen sources, consistent with known biospecimen-specific chromatin accessibility patterns^34^. For WGBS data, variable DNA methylation segments yielded the highest classification performance (∼95%).

Using SHAP-based feature attribution analyses, we investigated the biological relevance of the large (100 kb) genomic regions driving biospecimen classification. Genes overlapping highly informative regions were enriched for biologically relevant Gene Ontology terms and exhibited strong biospecimen-specific epigenomic signatures, as illustrated by Epilogos visualization of a representative locus (**Fig. 4e-f**). In addition, informative regions were significantly enriched for regulatory activity as defined by the ChromActivity framework (**Fig. 4g**), further supporting their functional relevance.

Given the strong performance of the EpiClass classifiers in predicting the assay, cancer status, sex and biomaterial type within EpiATLAS (**Supplementary Fig. 4d**), we applied these models to annotate more than 350,000 uniformly reprocessed datasets from the ChIP–Atlas^35^ and Recount3^36^ resources, for which key metadata attributes were frequently incomplete or absent at submission (**Fig. 4h**). This large-scale metadata inference substantially enhances the usability, consistency and biological interpretability of these widely used public epigenomic collections^32^.

### EpiATLAS enables cell type mapping across bulk and single-cell modalities

To explore the utility of EpiATLAS as an informative reference for single-cell studies, we next assessed its capacity to support the annotation of single-cell datasets and to infer cell type contributions within bulk epigenomic profiles. Single-cell chromatin accessibility and transcriptome data from human bone marrow and peripheral blood mononuclear cells (PBMCs)^37^ were integrated with blood-derived EpiATLAS samples (**Supplementary Table 4, 5**).

Joint embedding of deep-coverage bulk RNA-seq datasets with single-cell RNA-seq profiles enabled projection of single-cell transcriptomes into the EpiATLAS reference space, facilitating transfer of curated cell type annotations and providing a unified framework for cross-analysis between bulk and single-cell data (**Extended Data Fig. 4**). This analysis revealed strong concordance between cell type assignments derived from single-cell annotations and those inferred using the EpiATLAS reference. Cells annotated as progenitors through unsupervised single-cell clustering were consistently assigned progenitor identities by EpiATLAS-based prediction. Similarly, myeloid populations, including CD14⁺ and CD16⁺ monocytes and dendritic cells, showed close correspondence with EpiATLAS samples annotated as monocytes or neutrophils, whereas B-cell and T-cell clusters mapped to their respective lymphoid counterparts while preserving distinctions related to lymphocytic memory states.

Reciprocally, projection of EpiATLAS transcriptomes into single-cell RNA-seq–derived embeddings enabled transfer of single-cell annotations to bulk samples, facilitating potential downstream applications like cell type deconvolution of EpiATLAS datasets using single-cell–resolved labels. This bidirectional mapping demonstrated overall strong agreement between inferred and reference cell type identities, highlighting the utility of EpiATLAS as a coherent framework for integrating bulk and single-cell transcriptomic landscapes (**Supplementary Fig. 5a,b**).

To enable joint analyses across bulk epigenomes and single-cell data modalities, we used the epigenetic signal in putative cis-regulatory elements (uCREs) defined based on the universal ChromHMM annotations, resulting in a shared feature space for analysing bulk epigenome and single-cell chromatin accessibility data. Within uCREs, H3K27ac data for different EpiATLAS samples resulted in a clear separation that corresponded to annotated cell types (**Fig. 5a**). Projection of single-cell chromatin accessibility profiles into the low-dimensional space defined by H3K27ac in putatively regulatory regions, followed by the prediction of EpiATLAS annotated labels, showed strong agreement between bulk epigenomes and single-cell ATAC–seq cell types defined by unsupervised analysis (**Fig. 5a,b**). Conversely, the projection of EpiATLAS samples into scATAC-seq-derived embeddings and subsequent prediction of single-cell labels facilitates refinement of EpiATLAS epigenomes using single-cell accessibility (**Supplementary Fig. 5c,d**).

**Figure 5.**
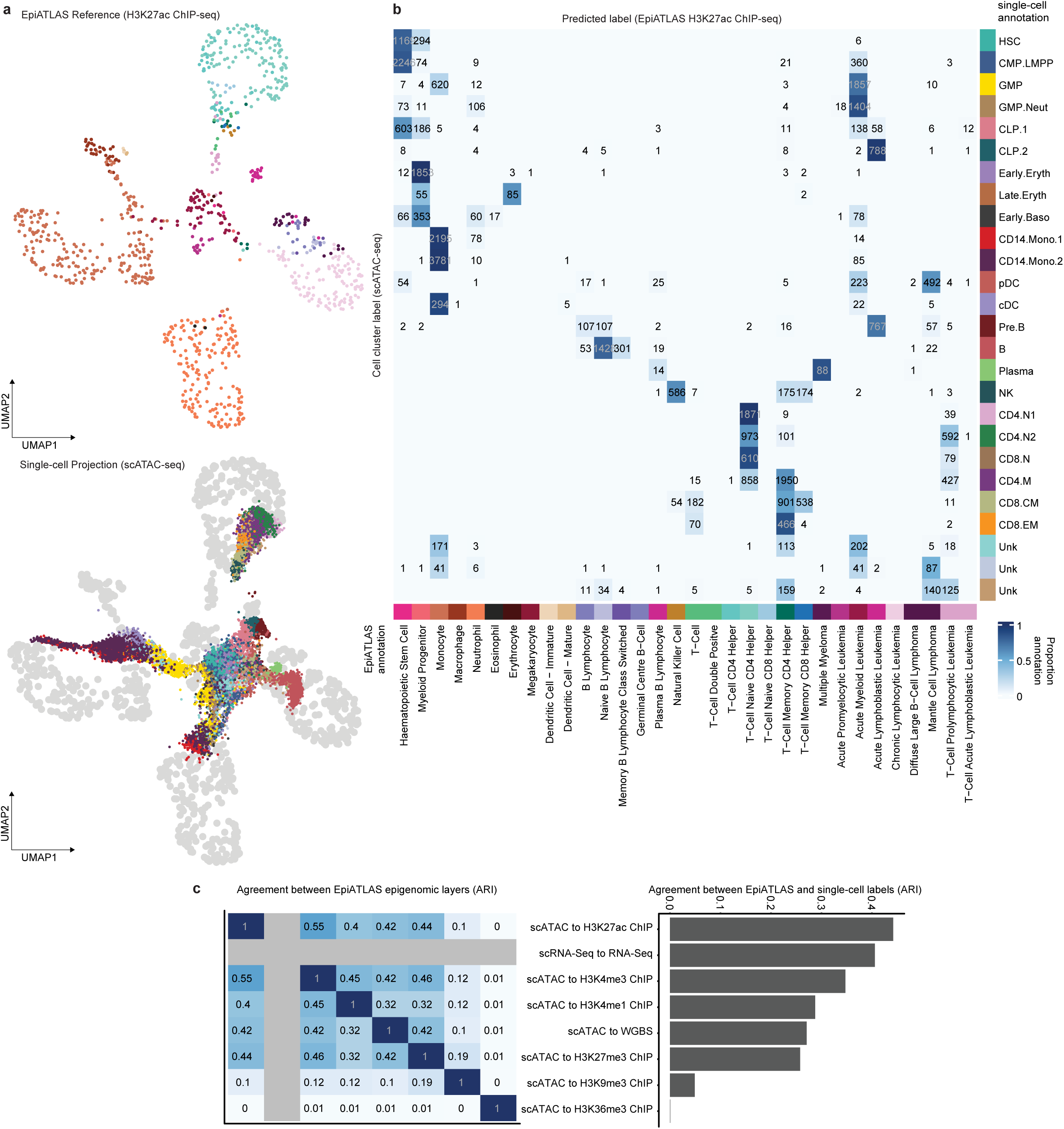
Mapping of single-cell epigenomes to the EpiATLAS reference. **a,** UMAP embedding of blood-related EpiATLAS H3K27ac epigenomes (top) and projection of single-cell accessibility data into the embedding (bottom). EpiATLAS bulk cell types are annotated from the ‘harmonized_sample_label’ metadata. Single-cell ATAC-seq data and cell type annotation were taken from Granja et al 2019^37^. In the projection UMAP, the EpiATLAS reference embedding is depicted as grey colored points. **b,** Heatmap denoting the number of predictions of single-cell ATAC-seq as EpiATLAS annotated cell types (‘harmonized_sample_label’). Values are absolute and colors are scaled by row. Predictions were made using classification with linear discriminant analysis (LDA) based on projected principal component embeddings. **c,** Comparison between different single-cell projection layers in their agreement of predicted EpiATLAS bulk labels with each other (left) and the single-cell cell type annotation (right). EpiATLAS cell type classification was performed using LDA based on projected principal component embeddings. Agreement was quantified by the Adjusted Rand Index (ARI).

Systematic comparison of cross-modality projections demonstrated high concordance between single-cell cluster identities and EpiATLAS-annotated reference cell types across epigenetic modalities including H3K27ac, gene expression, H3K4me3 and H3K4me1, as quantified by the adjusted Rand index (ARI) (**Fig. 5c, right**). These modalities also showed strong mutual agreement in their predicted EpiATLAS cell type assignments (**Fig. 5c, left**), indicating consistent capture of cell identity across active regulatory layers. In contrast, H3K36me3 and H3K9me3 exhibited reduced cell type separation and lower concordance with other modalities, consistent with their broader genomic distributions and association with domain-level chromatin organization rather than lineage-specific regulation.

Reciprocal prediction of single-cell labels from embedded EpiATLAS samples yielded similarly strong concordance for transcriptional profiles and enhancer-associated marks, whereas broader or repressive features—including H3K27me3, H3K9me3 and DNA methylation—showed comparatively lower agreement (**Supplementary Fig. 5e**). Together, these findings demonstrate that EpiATLAS provides a robust reference framework for integrating bulk and single-cell epigenomic datasets, enabling cell type-resolved interpretation across diverse regulatory modalities.

### Genome-wide maps of enhancer–gene regulation and transcript-level control

The EpiATLAS provides a unique resource for systematic analysis of enhancer-gene interactions and transcriptional regulation. Using the uCREs, we linked enhancers to target genes in biospecimens with H3K27ac ChIP–seq data (n = 1565) using the gABC framework^23^. Aggregating predictions across biospecimens yielded 21,106,658 enhancer–gene interactions (mean per biospecimen: 390,810) (**Fig. 6a)**. The predicted interactions varied across biospecimens, enabling separation of cell and tissue types, while biological replicate samples showed high concordance (**Extended Data Fig. 5a–c**).

**Figure 6.**
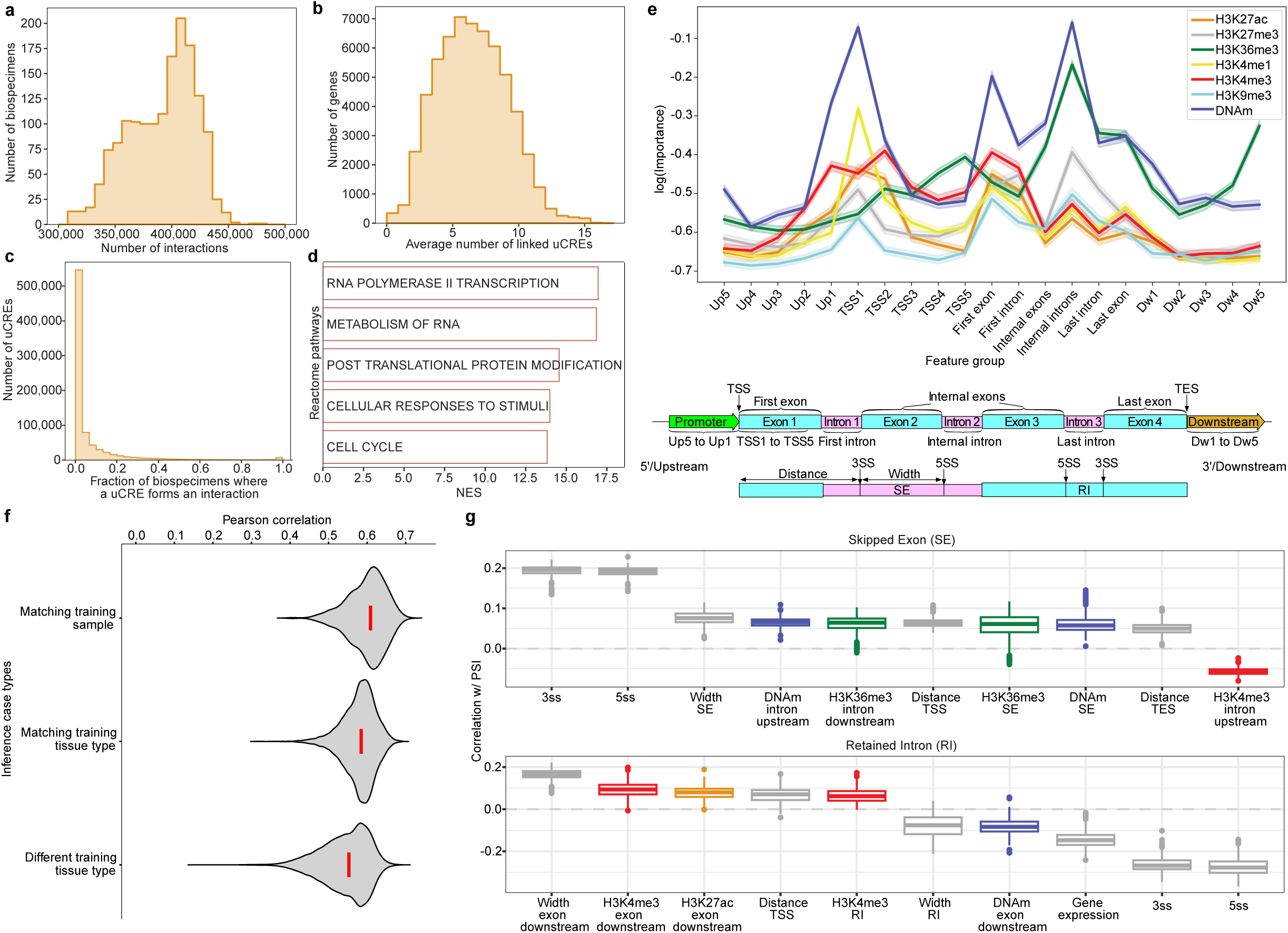
Epigenetic control of gene expression and splicing. **a,** Number of predicted interactions (x-axis) between cis-regulatory elements based on the universal ChromHMM annotation (uCREs) and genes per biospecimen. **b,** Average number of uCREs per gene across biospecimens, only considering genes with at least one uCRE. **c,** For each uCRE, the percentage of biospecimens in which it is targeting at least one gene (x-axis) is quantified for all 808,470 uCREs with at least one target gene (y-axis), i.e., minimum 1/number of biospecimens. **d,** Gene set enrichment of genes sorted in ascending order by their standard deviation of the number of interactions across biospecimens. Shown are the five terms with the highest normalised enrichment score (NES) (Reactome v2024.1^45^), calculated on the sorted genes with GSEApy^39,46,47^. A positive NES means that the annotated genes are enriched at the upper end of the list and have a low standard deviation. **e,** Normalised feature importance of graph convolutional networks (y-axis, log scale) predicting transcript expression based on the signal from histone modifications and DNA methylation. We differentiate between the gene body (first, internal, and last intron/exon), upstream (Up), and downstream (Dw) sub-regions (x-axis) across all samples (400 bp each). Line shading denotes the 95% confidence interval. **f,** Modeling performance quantified by Pearson correlation between computationally inferred transcript expression levels and their experimental measurements. All the training-testing pairs are divided into three categories: the same sample (top), different samples with the same tissue annotation (middle), or different tissues (bottom) (metadata column ‘tissue_type’). **g,** Partial Pearson correlation between (epi-)genomic features and Proportion Spliced-In (PSI) value controlling for gene expression for the ten highest absolute median correlations per event type. The dashed line depicts zero correlation. Box plots show centre line, median; box limits, upper and lower quartiles; whiskers, 1.5× interquartile range; points, outliers. SE: skipped exon; RI: retained intron; SS: splice site.

On average, each gene was associated with 6.5 enhancers (**Fig. 6b**), and most enhancer–gene interactions were highly context specific: 82% of interacting enhancers were linked to target genes in ≤10% of biospecimens (**Fig. 6c**). Genes exhibiting low variability in regulatory input across biospecimens were enriched for constitutive biological processes, including RNA metabolism and cell-cycle regulation (**Fig. 6d**). These genes were also significantly enriched for disease-associated genes from DisGeNET^38^ and for ubiquitously expressed genes (normalized enrichment scores^39^ 52.2 and 25.5, respectively; both P ≤ 0.001).

As a complementary approach, we correlated H3K27ac signal at enhancers with RNA-seq gene expression for all enhancer–gene pairs within 250 kb distance (n = 7,865,786). Although partial correlation analysis controlling for co-expression improved agreement with CRISPRi- and eQTL-supported interactions, performance remained substantially lower than that of the gABC score (**Supplementary Fig. 6**). Consistent with this, very few correlations remained significant relative to a shuffled background^40^ (14 interactions at FDR ≤ 5%; **Supplementary Fig. 6c**).

We next examined how epigenomic features relate to transcript-level expression output. We trained graph convolutional network (GCN) models integrating EpiATLAS data, averaged chromatin contact maps (Hi-C), and protein–protein interaction networks^41^ to predict expression of individual transcript isoforms. Unlike previous approaches, these models operate at transcript resolution and are trained predominantly on tissue-derived samples. Feature importance analysis revealed that DNA methylation and H3K36me3 signals within gene bodies, particularly at internal introns, were the strongest predictors of transcript expression, exceeding the contribution of promoter-associated marks (**Fig. 6e**; **Extended Data Fig. 5d**).

The GCN models generalized well to unseen data, achieving a median Pearson correlation of 0.60 when predicting expression on held-out chromosomes from the same sample, 0.55 when transferring across individuals within the same tissue, and 0.52 when transferring across tissue types (**Fig. 6f**). These results indicate robust modelling of transcript expression across diverse biological contexts, while retaining sensitivity to sample- and tissue-specific effects.

Finally, we investigated the relationship between epigenomic features and alternative splicing, which has been implicated in cell type–specific regulation and disease^42,43^. Genome-wide partial correlation analyses revealed that, alongside intrinsic sequence features such as splice-site strength and exon length, epigenetic signals in and around alternatively spliced regions are associated with exon and intron inclusion levels, independent of gene expression, (**Fig. 6g**)^44^. DNA methylation and H3K36me3 were positively associated with exon inclusion, whereas upstream H3K4me3 showed a negative association. In contrast, DNA methylation and H3K4me3 associations are inverted for intron inclusion, while H3K27ac in downstream exons showed a positive correlation.

### Epigenetic regulation of metabolic networks across cell types

EpiATLAS provides a unique opportunity to investigate epigenetic regulation at the level of biological networks, including cellular metabolism. We integrated EpiATLAS RNA-seq profiles with Recon3D, a curated and comprehensive generic human metabolic reconstruction, and contextualised the models using the rFASTCORMICS framework to generate metabolic models for 1555 samples spanning 58 cell and tissue types^48,49^. These models were then combined with EpiATLAS ChromGene annotations and cell and tissue type-specific enhancer–gene interactions derived using gABC framework to examine epigenetic control across metabolic pathways. We focused on the expression of 1844 metabolic genes across 33 cell and tissue types with at least three samples each (**Fig. 7a**). Metabolic genes spanned a continuum from ubiquitously and highly expressed loci encoding core metabolic enzymes and transporters to genes exhibiting pronounced cell type-restricted expression, particularly in hepatocytes. Consistent with these expression patterns, broadly expressed genes were associated with active ChromGene annotations and extensive networks of highly active enhancers, whereas selectively expressed genes displayed engagement of cell type-specific enhancer repertoires (**Fig. 7a**). Notably, genes expressed specifically in hepatocytes displayed particularly strong enhancer activity, consistent with the high metabolic demands of this cell type^50,51^ (**Fig. 7a**, right columns).

**Figure 7.**
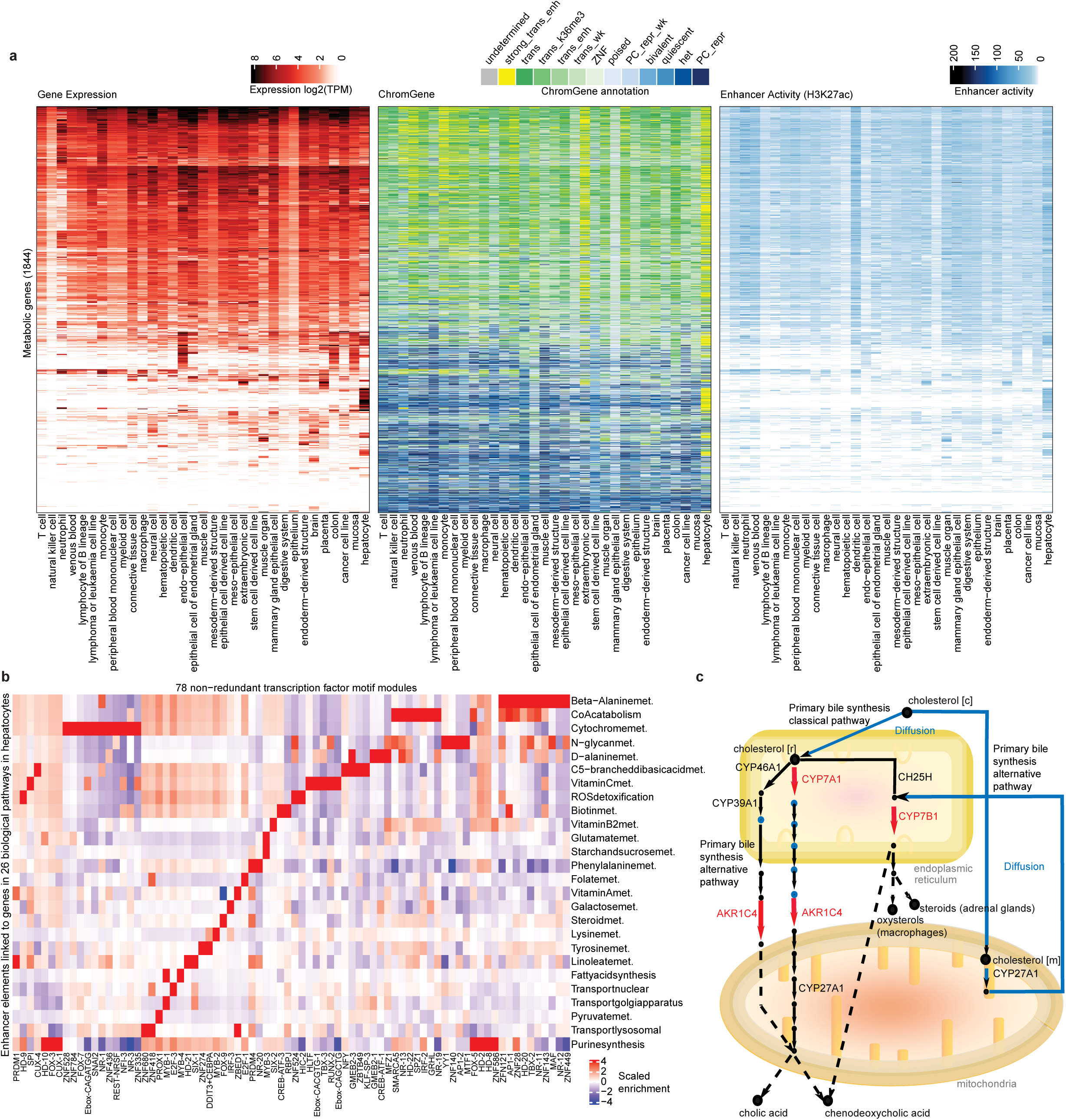
Epigenetic control of metabolic pathway activity. **a,** Heatmap depicting the mean gene expression (in log2 TPM), the ChromGene annotation for the majority of the samples or undetermined if there was no such annotation and the mean of the sum of the enhancer activities for 1844 metabolic genes across 33 cell and tissue types (metadata column ‘harmonized_sample_ontology_intermediate’). The heatmap is clustered by rows. **b,** Transcription factor module enrichment map showing association between 78 transcription factor modules and enhancers linked to genes in 26 biological pathways active in hepatocytes (met.: metabolism). The heatmap displays scaled enrichment values capped at a saturation threshold of 5 (equivalent to 32-fold enrichment) for ease of visualization. **c,** Reduced schematic of the primary bile synthesis pathway in hepatocytes and positioning of reactions controlled by liver-specific genes (red). These reactions are located at key positions in the network such as initial transporter or first reactions after diffusion steps (blue).

Using transcription factor (TF) binding site annotations and enhancer–gene interactions, we identified 78 of 280 predefined TF modules enriched in pathway-associated enhancers across 26 metabolic pathways in hepatocytes^52^ (**Fig. 7b**). Several of these TFs have established roles in metabolic regulation, including E2F family members in folate metabolism^53^ and PROX1 in pyruvate metabolism^54^, supporting the biological relevance of the inferred regulatory programs.

We next examined cell type-specific metabolic regulation at the network level. For each pathway, we visualized epigenomic and transcriptomic activity and defined cell type-specific epigenetic control points as genes showing predominantly active ChromGene annotations in a single cell type and predominantly repressive states in all others (**Fig. 7c**). This analysis identified 121 metabolic genes with cell type-specific active chromatin states. Integration of epigenomic and transcriptional data highlighted, for example, hepatocyte-specific regulation of primary bile acid synthesis, driven by selective expression of *CYP7A1*, *CYP7B1* and *AKR1C4*. These genes catalyse early steps following compartmental transitions in bile acid metabolism and are uniquely targeted by highly active enhancers in hepatocytes in our analysis (**Supplementary Table 6**). Across all pathways, metabolic genes with cell type-specific active chromatin states were significantly enriched for transporters (hypergeometric P = 0.031) and cytochrome P450 enzymes (hypergeometric P = 3.4 × 10⁻⁵) relative to all metabolic genes, consistent with specialised metabolic functions across tissues^49^. Together, these results demonstrate how EpiATLAS enables dissection of epigenetic control mechanisms operating at the level of metabolic networks.

### Epigenomic annotation of disease risk across traits, tissues and regulatory mechanisms

EpiATLAS provides a comprehensive framework for investigating the epigenomic basis of human disease. Among the 2279 references, 625 were associated with at least one disease annotation, spanning cancer, developmental, autoimmune, metabolic, inflammatory, vascular and infectious disorders (**Fig. 8a**). Blood- and immune-related diseases were particularly well represented, including cancers and autoimmune conditions.

**Figure 8.**
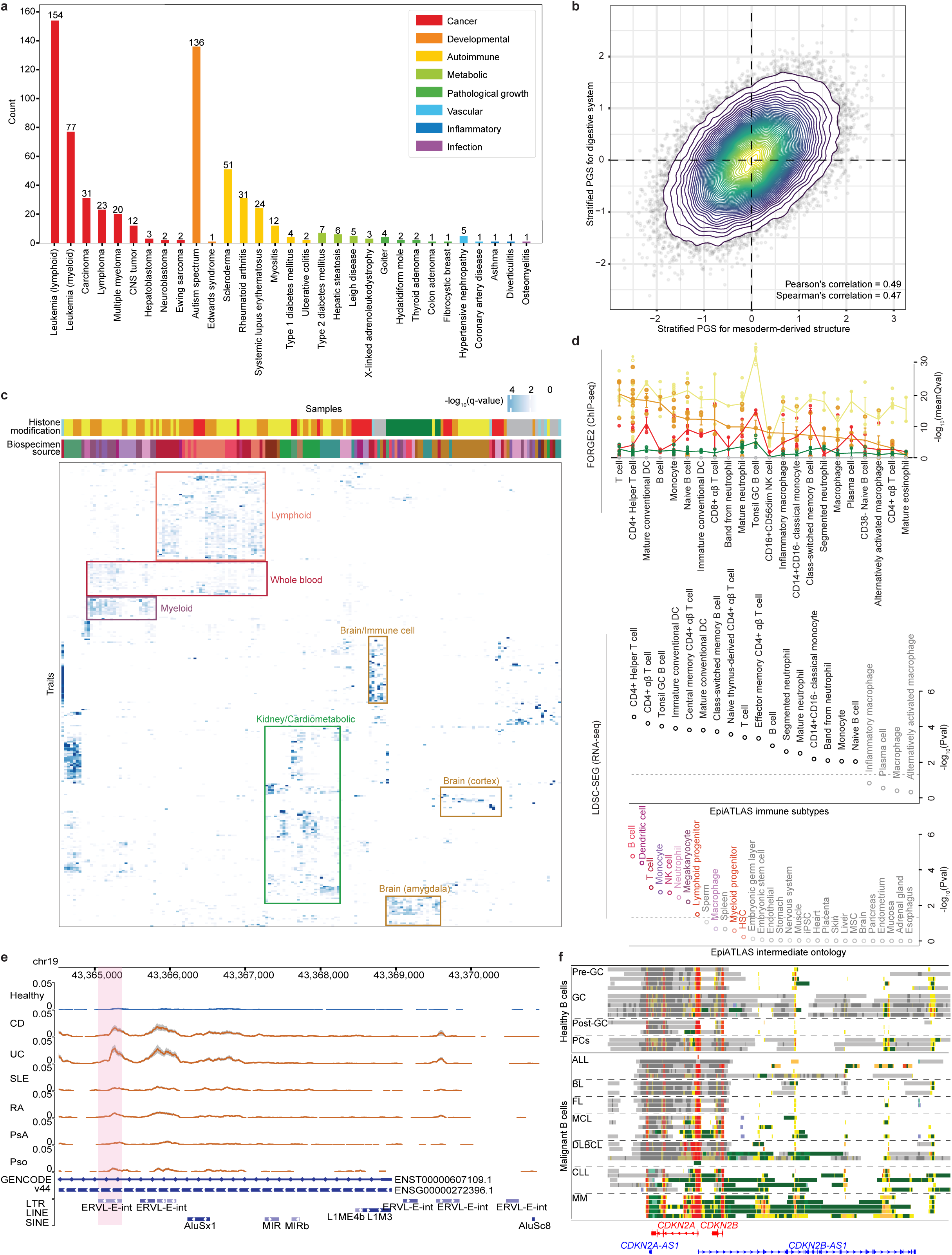
Use of EpiATLAS in disease studies. **a,** Overview of the disease categories in the EpiATLAS dataset (n=625). **b,** Tissue-stratified polygenic score (PGS) partitioning for standing height using EpiATLAS annotations and individual-level genetic data from the n_test_ = 67,730 white British individuals in the UK Biobank resource. The contour plot shows the distribution of individuals based on their stratified PGS contributions (Z-scores) from the digestive system and mesoderm-derived tissue enhancer regions, highlighting inter-individual variability in the dominant tissue-specific components of genetic liability. We show the Pearson and Spearman correlation coefficients between the two stratified scores. **c,** Global visualisation of FORGE2 functional overlap analysis (FOA) of all GWAS variants (version Dec-19-2021) with the six types of EpiATLAS histone modification data, clustered by traits and samples. Here, FOA of coding and non-coding variants integrates data from multiple regulatory and annotation levels (including histone mark broad peaks across thousands of datasets) to establish the biospecimen tissue- and cell type-specific architecture of disease-associated regulation. Highlighted are different tissue- and cell type-specific clusters, such as a brain/immune cell cluster for age associated/neurodegenerative conditions, such as “Alzheimer’s disease age at onset”, “cognitive decline measurement”, “ischemic stroke”, “late onset Alzheimer’s disease”, as well as GWAS for inflammatory markers, e.g. “interleukin 9 measurement”. **d,** Linkage disequilibrium score regression of specifically expressed genes (LDSC-SEG) analysis based on EpiATLAS RNA-seq data reveals that MS susceptibility is largely associated with blood/immune cells. Further subgrouping shows that CD4+ Helper T cell is the most significant cell type both in LDSC-SEG (RNA-seq) and FORGE2 H3K27ac (ChIP-seq) analyses (error bars showing standard error of the mean). Colour key as in Fig 1. **e,** Dysregulation of repetitive elements in Chronic Inflammatory Diseases (CID) patients. A genome browser screenshot showing an example of RNA-seq of a differentially expressed individual repetitive element. The pink shading highlights a disease-specific individual ERVL-E-int element that is significantly upregulated (Fold change > 4 and two-tailed Wald test p.value < 1e-4) only in Inflammatory Bowel Diseases patients (CD and UC) but not in other disease patients (SLE, RA, Pso, PsA). The blue and orange lines represent the mean expression value across all healthy controls and patients with the disease, respectively. The grey shades denote the standard errors. All datasets are displayed as Reads Per Million mapped reads (RPM) with y-axis ranging from 0 to 0.05. Healthy (N = 197); CD: Crohn’s Disease (N = 155); UC: Ulcerative Colitis (N = 107); SLE: Systemic Lupus Erythematosus (N = 66); RA: Rheumatoid Arthritis (N = 128); Pso: Psoriasis (N = 38); PsA: Psoriatic Arthritis (N = 93). **f**, Chromatin state segmentation of the *CDKN2A/B* locus, which encodes key tumor suppressors and includes the antisense transcripts *CDKN2A-AS1* and *CDKN2B-AS1*, reveals a largely conserved regulatory architecture across normal B-cell differentiation. In contrast, malignant samples show increased heterogeneity and redistribution of chromatin states. Active promoter-associated states around transcription start sites remain broadly conserved, whereas elongation-associated states are expanded. Together, these patterns are consistent with substantial cancer-specific remodelling of chromatin organisation at the locus and may also be influenced by underlying genetic alterations, such as copy-number loss. GC: germinal center; PCs: plasma cells; ALL: acute lymphoblastic leukemia; BL: Burkitt lymphoma; FL: follicular lymphoma; MCL: mantle cell lymphoma; DLBCL: diffuse large B-cell lymphoma; CLL: chronic lymphocytic leukemia; MM: multiple myeloma.

To assess how EpiATLAS annotations inform tissue-specific genetic contributions to complex traits at the individual level, we applied biosample-stratified polygenic score partitioning using inclusive polygenic scores^55^ (iPGS) and EpiATLAS-derived ChromHMM enhancer annotations. By decomposing individual polygenic scores into biosample-specific components, we inferred the relative contribution of different tissues to genetic liability. Using standing height as an example trait, we evaluated stratified scores in a held-out test set of 67,730 individuals from the UK Biobank. Genetic signal was most strongly enriched in enhancer regions from digestive system and mesoderm-derived tissues, but the relative contribution of these tissues varied substantially across individuals (**Fig. 8b**). Although stratified scores were moderately correlated (Spearman’s ρ = 0.47), the observed inter-individual variability highlights the potential of EpiATLAS-guided polygenic profiles to provide interpretable, tissue-resolved views of genetic risk.

To systematically characterise disease-associated regulatory architectures, we performed a global phenotype–tissue association analysis by intersecting genome-wide association study (GWAS) loci with EpiATLAS histone modification data. Using FORGE2^56^, we tested enrichment of GWAS variants from the GWAS Catalog (December 19, 2021) in six histone modification peak sets across tissues and cell types. We identified 317 significant phenotype–tissue associations (q < 0.01), representing a 1.84-fold increase compared with analogous analyses using consolidated Roadmap Epigenomics data^16^ (**Fig. 8c, Supplementary Fig. 7**). These associations formed coherent tissue-specific clusters, including immune and brain-related samples enriched for age-associated and neurodegenerative traits, myeloid clusters enriched for haematological traits, and lymphoid clusters enriched for autoimmune diseases. Additional clusters corresponding to kidney and brain subregions showed enrichments consistent with known disease aetiologies (**Supplementary Table 7**). This analysis highlights the systems-level relevance of the updated larger IHEC dataset, which can be used to improve our understanding of disease-associated regulation.

We next examined a specific disease context in greater detail by analysing multiple sclerosis (MS) susceptibility. Stratified linkage disequilibrium score regression of specifically expressed genes (LDSC-SEG) using EpiATLAS RNA-seq data and MS GWAS summary statistics^57^ revealed strong enrichment in immune cell types (**Fig. 8d**). T and B lymphocytes, as well as dendritic cells, ranked highest, with further stratification implicating CD4⁺ helper T cells and tonsillar germinal centre B cells. Consistent results were obtained using FORGE2 analysis of H3K27ac and H3K4me1 ChIP–seq data, with previously reported MS susceptibility polygenic risk score (PRS) SNPs as input^58^, where CD4⁺ helper T cells were again the top-enriched immune subtype (see highlighted SNP in **Fig. 2**). In contrast, macrophages and plasma cells showed little enrichment, and repressive marks such as H3K9me3 and H3K27me3 were not associated with MS susceptibility, supporting a central role for active regulatory programs in antigen-specific immune responses. A recent MS GWAS, focusing on disease severity, was enriched in the central nervous system^59^ (see highlighted SNP in **Extended Data Fig. 2**).

Given the extensive epigenomic reprogramming observed in chronic inflammatory diseases (CIDs), we next investigated whether dysregulation of repetitive elements accompanies disease-associated regulatory changes. Analysis of mRNA-seq data from CID patients and controls^60^ (n = 784) revealed aberrant expression of multiple repetitive elements (**Fig. 8e, Extended Data Fig. 6a**). Notably, a specific ERVL-E-int element showed strong and selective upregulation in inflammatory bowel disease (Crohn’s disease and ulcerative colitis), but not in other CID cohorts, including systemic lupus erythematosus, rheumatoid arthritis and psoriatic arthritis (**Extended Data Fig. 6b**). This element showed minimal enrichment for active histone modifications in healthy EpiATLAS samples and no detectable activity at the ERVL-E subfamily level (**Fig. 3g**), highlighting the importance of analysing transposable elements at individual-element resolution.

Finally, we leveraged the large collection of harmonised cancer epigenomes in EpiATLAS to identify cancer-specific regulatory alterations. Comparing healthy B cells with malignant counterparts, we analysed chromatin state patterns at the *CDKN2A/CDKN2B* tumour suppressor locus, which is frequently silenced or deleted in cancer^61,62^. Promoter-associated activity marks (co-occurrence of H3K27ac and H3K4me3) were largely preserved between healthy and cancer samples. In contrast, cancer-specific transcriptional elongation states marked by H3K36me3 were observed in subsets of mantle cell lymphoma, diffuse large B-cell lymphoma, chronic lymphocytic leukaemia and multiple myeloma (**Fig. 8f**). These elongation states predominantly overlapped the antisense transcript CDKN2B-AS1, which was overexpressed in cancers showing the strongest *CDKN2A* signal (**Extended Data Fig. 6c,d**). Together, these results illustrate how EpiATLAS enables discovery of disease- and cancer-specific regulatory mechanisms across multiple layers of epigenomic control.

## Discussion

EpiATLAS provides the largest harmonised collection of human epigenomes to date, accompanied by standardised metadata models, containerised and reproducible workflows, and improved mechanisms for consent-aware data sharing within the international epigenomics community. By unifying thousands of epigenomic profiles across diverse tissues, cell types, and individuals, EpiATLAS establishes a foundational reference resource designed not only for immediate reuse but also to support the next generation of large-scale epigenomic studies. Importantly, these efforts were developed within the broader framework of IHEC, whose ELSI working group helped shape international discussions surrounding responsible epigenomic data generation, governance, sharing, and consent. This multidisciplinary collaboration between biological and social scientists provided an important model for integrating ethical and societal considerations directly into large-scale reference epigenome initiatives.

A central advance of the EpiATLAS is the development and adoption of standardised metadata models for epigenomic datasets. These models provide a common framework for describing biological context, experimental design, and technical provenance, addressing a longstanding barrier to interoperability across consortia. Their uptake by national and international initiatives and by the Global Alliance for Genomics and Health (www.ga4gh.org) underscores the need for, and feasibility of, community-driven standards to enable scalable data sharing while respecting ethical and consent constraints. Together with containerised workflows, these metadata standards ensure that analyses are reproducible, portable, and extensible, facilitating both reanalysis of existing data and seamless integration of future datasets.

Analyses enabled by this resource provide new insights into the structure and variability of the human epigenome. Within biospecimen types, we observe substantial dispersion in epigenomic profiles between individuals, consistent with prior large-scale studies suggesting that inter-individual epigenomic variation is a pervasive feature of human biology. Although harmonised processing and standardised workflows reduce known sources of technical heterogeneity, fully disentangling technical contributions from biological signal remains challenging. Nevertheless, our analyses are concordant with regulatory mapping efforts^63^ indicating that a large fraction of the human genome can be marked by regulatory elements in a cell type-specific manner. These findings argue strongly for the next phase of reference epigenome profiling to move beyond single exemplars of canonical cell types toward designs that explicitly capture inter-individual variation within diverse populations in relation to phenotypic variables such as age, biological sex and body mass index.

EpiATLAS further provides a reference coordinate framework for interpreting both bulk and single-cell epigenomic data. Integration of bulk and single-cell atlases yields complementary advantages: bulk data provide deep, whole-genome coverage, while single-cell profiles offer high-resolution views of cellular heterogeneity. Joint analyses mitigate sparsity inherent to single-cell data while enriching bulk analyses with refined cell type annotations. Notably, putative regulatory regions defined by EpiATLAS serve as informative features for single-cell analyses, improving interpretability and enabling cross-modal comparisons.

Beyond descriptive annotation, EpiATLAS supports predictive and mechanistic analyses. Sample-specific predictions of uCRE–gene interactions constitute a rich resource for downstream investigations, as illustrated by the identification of regulatory regions linked to specific metabolic pathways and the transcription factors enriched within them. These analyses highlight the capacity of integrated epigenomic data to illuminate regulatory programs underlying complex cellular phenotypes. In addition, EpiATLAS corroborates accumulating evidence that DNA methylation and histone modifications are associated with alternative exon inclusion and intron retention, reinforcing the intimate coupling between chromatin state and RNA processing. However, despite these associations, disentangling causality remains challenging, emphasising the need for perturbation-based and temporal studies to resolve directionality.

Importantly, EpiATLAS enables the discovery of epigenome-mediated disease associations that are not readily apparent from genetic data alone. By integrating disease-relevant epigenomic states with phenotype information, we identify novel phenotype–tissue and risk associations. Epigenomic state analysis further reveals that individual repeat elements can be dysregulated in a disease-specific manner, exemplified by their selective involvement in inflammatory bowel disease. Moreover, integrative analyses combining GWAS with EpiATLAS shed light on disease-associated variants that exert their effects in specific cellular contexts, as illustrated by multiple sclerosis risk variants acting in distinct immune cell populations. These findings highlight the value of epigenomic context for interpreting non-coding genetic variation and for refining disease mechanisms.

Despite its breadth, EpiATLAS has limitations. While harmonisation reduces technical variability, residual differences in experimental design and sample composition persist. Moreover, most datasets represent static snapshots, limiting inference about temporal dynamics and causal relationships. Addressing these challenges will require systematic inclusion of longitudinal data, perturbation experiments, and deeper integration with multi-omic modalities, including transcriptomic, proteomic, and spatial data.

In summary, EpiATLAS provides a comprehensive, interoperable reference for human epigenomics, coupling large-scale data harmonisation with analytical frameworks. By capturing both shared regulatory architectures and meaningful biological variation, this resource lays the groundwork for future studies of human development, cellular identity, and disease, and offers a scalable model for community-driven data sharing in functional genomics.

## Methods

For information on the methods, refer to the accompanying document.

## Supporting information

Supplementary Material Details

## Extended and supplementary data

Extended and supplementary figures can be found in the accompanying document. Supplementary tables and data can be found in the accompanying archive.

## Data Availability

Processed EpiATLAS data are available on the IHEC data portal (https://epigenomesportal.ca/ihec/) and the IHEC website (https://ihec-epigenomes.org/epiatlas/data/). Source data are available through accessions found in the epigenome reference registry (https://www.ebi.ac.uk/epirr) and access requests to respective public or protected repositories. MS cohort data are stored and managed by the PopGen biobank at the Institute of Epidemiology at Kiel University, Germany. Researchers can apply for data access to the Kiel cohort data by submitting a research proposal, including the scientific background, research question, success prospects, study design, potential conclusions, and scientific collaborators of their study, at the local biobank P2N via the following form: http://www.uksh.de/p2n/Information+for+Researchers.html. Due to the informed consent obtained from the participants, phenotypes, as well as RNA-sequencing data, cannot be accessed without restriction.

## Acknowledgements

The authors would like to express their sincere gratitude to the many individuals, institutions, and funding agencies whose contributions, leadership, and support have shaped the International Human Epigenome Consortium (IHEC) over the past decade. The success of IHEC and the realisation of this work are the result of a truly global collaborative effort involving hundreds of investigators, trainees, technical staff, program managers, and administrators whose collective dedication has advanced the field of human epigenomics and enabled the creation of an unprecedented reference resource for the scientific community.

While it is impossible to recognise every individual contribution, we wish to extend our deepest appreciation to all those who have supported the consortium throughout its evolution. Their vision, commitment, and collaborative spirit have been essential to achieving the scientific goals of IHEC.

We offer special thanks to **Dena C. Procaccini**, **Eric R. Marcotte**, **John S. Satterlee**, and **Tomasz Dylag**, whose tireless efforts provided critical leadership, coordination, and organisational cohesion across the consortium. Their dedication to fostering collaboration, maintaining momentum, and navigating the complexities of a large international scientific enterprise created the essential framework that enabled this work to be completed. Their contributions have been instrumental not only to the success of this project but also to the enduring impact and legacy of IHEC.

We would like to acknowledge Genome British Columbia (C32EMT, C41EMT) and the Canadian Institutes of Health Research (EP1-120589, CEE-151619, PJT-175218) as part of the Canadian Epigenetics, Environment and Health Research Consortium Network (CIHR-262119, EPT-165702). Additional support was provided by FEDER/Spanish Ministry of Science and Innovation (PID2023-148272OB-I00). This work was also partially supported by awards to EMBL-EBI from the Wellcome Trust for the Ensembl Project [222155/Z/20/Z, 226083/Z/22/Z, 226458/Z/22/Z] and the European Commission Horizon 2020 project MultipleMS [Grant Agreement Number 733161]. The work was further supported by European Union’s Horizon 2020 research and innovation programme SYSCID under grant agreement No 733100 and the DFG Cluster of Excellence Precision Medicine in Chronic Inflammation, EXC 2167/2, 390884018. Q. Manz was supported by a fellowship within the IFI programme of the German Academic Exchange Service (DAAD). N. Baumgarten, F. Behjati Ardakani, L. Rumpf, S. Ashrafiyan, D. Hecker, and M. H. Schulz have been supported by the DZHK (German Centre for Cardiovascular Research, 81Z0200101 and 81Z0200113), the Cardio-Pulmonary Institute (CPI) [EXC 2026] ID: 390649896, the DFG SFB TRR267 (Z03, project ID 403584255), the DFG SFB1531 (S03, project number 456687919), and the Hessian.AI center. This work was partially supported by the Natural Sciences and Engineering Research Council (NSERC) of Canada to P.-É. Jacques [#435710]. J. Raby is the recipient of a Graduate Scholarship from NSERC and the Fonds de Recherche du Québec – Nature et Technologie (FRQNT). P.-É. Jacques holds a Fonds de Recherche du Québec – Santé (FRQS) Research Scholar Senior Career Award. This research was enabled in part by computing support provided by Calcul Québec and the Digital Research Alliance of Canada. The imputation used computational and storage services associated with the Hoffman2 Cluster which is operated by the UCLA Office of Advanced Research Computing’s Research Technology Group and the Mammouth computing infrastructure of the Université de Sherbrooke. This work was also partially supported by the Hong Kong Epigenome Project (Lo Ka Chung Charitable Foundation), the Hong Kong Research Grant Council (GRF16104122 and GRF16102124), and the Croucher Innovation Award. D. Gérard and L. Sinkkonen were supported by funding from Fondation du Pélican de Mie et Pierre Hippert-Faber. J. Ernst was supported by the National Institutes of Health (DP1DA044371, U01HG012079, R21AG092008, P30AG073104). This work was supported by ELIXIR-DE (de.NBI), the research infrastructure for life science data to N. Aggarwal (grant number W-de.NBI-021).

## Author Contributions

**Co-First** Q. Manz, M. Bilenky, D. Hecker; **Lead Analysts** N. Aggarwal, J. E. Arcila-Galvis, S. Ashrafiyan, N. Baumgarten, F. Behjati Ardakani, P. R. Branco Lins, C. E. Breeze, D. Brownlee, D. Bujold, A. R. Chapman, S. H. Chow, T. U. Dincer, C. Dupras, G. Frosi, J. Fu, D. Gérard, A. Hauduc, J. Hyacinthe, A. Jaroszewicz, R. Li, R. J. Mangan, A. Mikulasova, I. Moghul, M. Needhamsen, N. Palmour, M. Pires Pacheco, J. Quon, J. Raby, A. Reynolds, L. Rumpf, A. Salhab, C. H. Shi, L. Sinkkonen, Y. Tanigawa, R. Tanner, H. Vu, F. White; **Production Team Members** J. T. M. Aw, S. Badii, R. Bowlby, M. Boyle, C. Brown, D. Chand, M. Calingo, Q. Cao, A. Carles, M. Caron, M. Carreira, A. S. L. Cheng, C. C. Y. Cheng, D. Cheng, Y. Cheng, M. Cheung, G. Choe, E. Chuah, R. Dali, A. Deng, S. Gakkhar, M. Gut, A. He, S. C. Heath, V. Ho, A. G. Intan, C. Y. Ito, Q. Jiang, J. L. Kluiver, J. Lee, S. Lee, Z. Leung, I. Li, A. Lorzadeh, D. McKerricher, N. Mishra, K. Mungall, L. A. E. Nagai, M. Ngubo, D. Palmquist, J. Pang, V. Park, P. Plettner, A. Puri, A. Redensek, Z. Sharafian, M. Simon, J. Steif, E. Su, S. K. M. Tam, S. S. T. Tam, A. Van den Berg, R. Wong, T. Wong, J. K. H. Wu, A. Zhu; **Production Team Leads** T. Kwan, M. Moksa; **IHEC PIs and PMs** S. T. Aherne, S. Aparicio, T. Arima, H. Chung, J. F. Costello, C. J. Eaves, S. J. Fisher, I. G. Gut, P. W. Harrison, B. Horsthemke, G. R. Ilsley, M. Jagodic, A. Karsan, P. M. Lavoie, D. Leung, M. W. Libbrecht, T. Manke, M. A. Marra, W. Meuleman, F. Müller, R. Nakato, H. Okae, D. Rico, P. Rosenstiel, H. Sasaki, T. Sauter, S. Schreiber, D. W. Scott, W. L. Stanford, C. Steidl, H. G. Stunnenberg, M. Suyama, F. Tran, T. Ushijima, A. P. Weng, S. M. Wiseman, A. R. Wu, K. Y. Yip, S. Yip; **Corresponding Authors** S. Beck, G. Bourque, J. Ernst, P. Jacques, Y. Joly, M. Kellis, M. List, T. J. Perkins, M. H. Schulz, J. Walter, M. Hirst.

