## Supplementary material for "EpiATLAS – a reference for human epigenomic research": Methods

### EpiATLAS Methods

#### Epigenome Mapping

*Data download verification and staging*

Data access applications for the IHEC EpiATLAS datasets were submitted to the European Genome-phenome Archive (EGA) (Blueprint, DEEP, Korea Epigenome Project (KNIH), CEEHRC, GIS, EpiHK), the database of Genotypes and Phenotypes (dbGaP) (NIH Roadmap Epigenomics), the National Bioscience Database Center (NBDC) (AMED-CREST), and the ENCODE portal (ENCODE) (**Supplementary Table 1**). Upon download, data files and accompanying metadata were reviewed, curated, and organized into a local database at Canada's Michael Smith Genome Sciences Centre to support analysis and tracking. Uniform processing was completed on 7539 analysis objects spanning RNA-seq, whole genome bisulfite sequencing (WGBS), and ChIP-seq analysis objects, representing 337 complete (that include 6 histone modifications ChIP-seq data, RNA-Seq and DNA methylation) and 1978 partial epigenomes. The final base data footprint was 28.8 TB, with an additional 6TB of post-processed data. A total of 923 analysis objects were excluded from analysis due to issues including corrupted files, data of non-human origin, or absence of required ChIP input controls. Raw sequencing data files from the collections were re-analysed in fully containerised workflows to guarantee bit-wise reproducibility.

Across assays, FASTQs were extracted either (i) from downloaded BAM files using Picard (v.2.4.1) *SamToFastq* or (ii) directly from SRA binary files via *fastq-dump*, retaining only reads that passed the Illumina chastity filter, when chastity information was available.

A full description of the harmonized processing can be found here (**Supplementary Data 1**) summarized below.

*Sequencing length and depth normalization*

Sequence files from multiple runs for the same experiment were merged into a single ANALYSIS object. RNA-seq reads were trimmed to 75 nt. ChIP-seq and WGBS datasets were left untrimmed unless specified (see experimental metadata). The final selection of ChIP-seq datasets resulted in the median sequence depth of 38 M and 39 M fragments for narrow marks (H3K27ac, H3K4me3) and 51 M, 52 M, 52 M and 54 M fragments for broad marks (H3K4me1, H3K36me3, H3K9me3, H3K27me3, respectively) (see details in **Supplementary Fig1a**). For the majority of ChIP-seq datasets the corresponding DNA input controls were available (median sequencing depth 57 M fragments). When not available, DNA input control from the closest dataset was used (i.e. from the same biospecimen source, but different donor) (**Supplementary Table 8)**. WGBS aimed for ~30X genomic coverage (resulting datasets have median sequence depth ~610 M fragments; **Supplementary Fig. 1b**), while the RNA-seq datasets have ~90 M fragments median sequencing depth (**Supplementary Fig. 1c**). Following preprocessing the EpiATLAS dataset was composed of 1639 trillion sequenced reads.

*Mixed Data Strategies*

For mixed single-end (SET) and paired-end (PET) datasets, PET data were preferred, given that the sequencing depth thresholds were met, otherwise SET data with only first read from the PET data added were used. A number of epigenomes contained multiple different read length sequence files. The following selection was applied: first, files with the same sequence length that had highest sequence depth were selected; if sequence depth was not sufficient, files with different sequence length were added; data with 75bp read length was favored. To minimize input redundancy, one representative input control dataset was randomly selected per reference epigenome.

*Identifier Assignment*

For each analysis object the unique UUID was generated and it was used in the name prefix of every data file resulting from the analysis - see **Supplementary Data 1** for details.

*Data processing*

All experiment metadata was collated and is made available on GitHub (https://github.com/IHEC/EpiATLAS_experiment_metadata/).

**ChIP-seq:** Fastq file were processed using the IHEC Integrative Analysis ChIP Pipeline Singularity container (https://github.com/IHEC/integrative_analysis_chip/releases/tag/v1.0) using BWA-MEM2 alignment to the GRCh38 "no-alt-analysis-set" reference (NCBI build GCA_000001405.15; identical copies hosted at EBI and epigenomes.ca). Median mapping rate was >95% for all histone modifications and DNA input (see **Supplementary Fig. 1a**).

Samples with low quality control metrics were manually reviewed and 144 experiments were excluded from the final collection, 134 for low QC or visual inspection and 10 that were mislabelled (**Supplementary Table 9**). This resulted in six biospecimens having ChIP-seq data completely removed from the EpiATLAS: IHECRE00000833, IHECRE00003135, IHECRE00003202, IHECRE00003229, IHECRE00003326, IHECRE00003355. ChIP-seq output files include sequence stripped BAM files, raw fragment coverage file (bigWig format) and results of MACS2 enrichment analysis - narrowPeaks files (with and without removal of enriched regions falling into annotated alignment artifacts regions [ENCODE: https://www.encodeproject.org/files/ENCFF356LFX/] for p-value threshold 0.01), as well as fold-change and -log10(p-value) coverage tracks (bigWig). Pipeline QC results were summarized and available in two formats: .json and .html. One per reference epigenome DNA input alignment (modified) BAM file as well as raw coverage file was generated. In addition to narrowPeaks, standalone MACS2 was run in a broadpeaks mode (*--broad*).

**RNA-seq** libraries were processed with *grape-nf* (release 2024-02-01, Docker sha256:8f64…) on the same GRCh38 assembly used for ChIP-seq and WGBS alignments, using GENCODE v29. Median mapping rate was ~98% (**Supplementary Fig. 1b**). Gene and isoform RSEM[1] quantification files (containing read counts, RPKM, and TPM metrics) as well as read coverage files and QC reports were available for each epigenome.

**Whole-genome bisulfite sequencing (WGBS)** data were run through *gemBS* (v.3.5.0) (Docker sha256:c4e7…) in PBAL-stranded mode when single- and paired-end runs co-existed. The median mapping rate was ~77% (see **Supplementary Fig. 1c)**. The methylating calling results are summarised in the bed format for CpGs and cytosine in non-CpG context. Detailed gemBS QC results were summarized for each dataset.

#### Sample Metadata Harmonization

The metadata harmonization working group adopted a collaborative, reproducible approach using OpenRefine (v.3.4.1) and GitHub (https://github.com/IHEC/epiATLAS-metadata-harmonization) to harmonize sample metadata across the different consortia included in EpiATLAS. Firstly, all metadata attributes for all available entries were pulled via HTTP API from the Epigenome Reference Registry (EpiRR) and merged into a shared working table. The group decided on relevant attributes to keep for harmonization and which analogous attributes to merge. Next, the remaining ones were distributed among members in a shared effort to harmonize their entries with OpenRefine, which allows for (semi-)manual editing of tables to solve simple inconsistencies and subsequent export of the changes in JSON format, which were applied programmatically by using the openrefine-client (v.0.3.10) and uploading the changes to GitHub for transparent and accessible tracking of changes. Complex inconsistencies were documented in GitHub issues and resolved in dedicated meetings. New versions of the harmonized metadata table resulting from this iterative process were tagged and released on GitHub. During the effort, the previously established metadata standards (https://github.com/IHEC/ihec-ecosystems) were followed for selecting appropriate ontology terms to describe sample origin for cell lines (Experimental Factor Ontology), primary tissues (Uberon), primary cells and primary cell cultures (Cell Ontology), and disease and health status (NCI Metathesaurus). Biospecimens were annotated as specifically as possible in the respective ontology. Based on sample origin and biomaterial type, biospecimens were also grouped into intermediate (Intermediate Biospecimen Label) and broad ontology terms (Broad Biospecimen Label). Finally, a curated label was added that describes biospecimens across ontologies, e.g., by capturing similar cell and tissue terms or including disease information (Curated Biospecimen Label). In addition to a core table containing the most important biospecimen attributes as described above and the EpiRR identifier, an extended metadata table was compiled. It consists of more detailed descriptions of the donor, automatically generated columns that simplify programmatic access to the table, and further manual annotations by additional members that offer alternative groupings into cell types and tissues.

#### Recurrence Analysis

##### *ChIP-seq*

We studied the enrichment recurrence frequency genome-wide for all 6 histone modifications and compared distributions for observed and imputed data. The following analysis was performed for each histone modification. First, for each epigenome, each genomic 50 bp was marked as enriched if it overlapped MACS2-identified narrow peaks for at least 25 bp. Otherwise, the bin was marked as not enriched. Then, for each bin, the recurrence frequency within EpiATLAS was defined as the number of epigenomes in which the bin was enriched, divided by the total number of epigenomes.

*RNA-seq*

We calculated recurrence properties for RNA-seq considering protein-coding genes (Ensembl v94) and set three discovery thresholds on TPM (as reported by *grape_nf* pipeline): 0.1, 1, 10. For each dataset, genes with TPM above a threshold were considered discovered, and recurrence frequency was calculated per gene as the ratio of the number of datasets with the gene to the total number of datasets. Since TPM distributions differ between mRNA (polyA-selected) and total-RNA (ribodepleted) libraries, we performed calculations separately for the two RNA-seq experiment strategies. Ubiquitously expressed genes, using recurrence frequency ≥ 0.99 for TPM threshold=1, were tested for overlap (hypergeometric test) against published lists of housekeeping genes[2,3].

*WGBS*

For DNA methylation, for every EpiATLAS dataset, we identified CpGs with coverage of at least 5 and marked CpGs as hypomethylated or hypermethylated, if fractional methylation was ≤ 0.25, or > 0.75, respectively. Then, for every CpG, we defined recurrence for both hypo- and hypermethylated CpGs. This was done for both observed and imputed DNA methylation data.

*Saturation*

The saturation study for a histone modification is intended to answer a question: how many ChIP-seq experiments for this histone modification will discover the entire potentially enriched genomic estate, i.e., adding a new experiment would barely add any new enriched locations. We used only datasets for which both observed and imputed data were available, to compare the saturation dynamics of both types of data.

**ChromImpute imputation of epigenomic data**

*ChIP-seq imputation from other ChIP-seq data*

We performed imputation of six IHEC histone modifications: H3K27ac, H3K27me3, H3K36me3, H3K4me1, H3K4me3, H3K9me3 for 1704 reference epigenomes with ChIP-seq data for at least one histone modification using ChromImpute (v.1.0.5)[4]. We used the -log p-value signal tracks at a 25 bp resolution as computed by the Convert command of ChromImpute. The imputation was restricted to chr1-22 and chrX. We used the '-tieglobal' flag to break ties. The number of input datasets for each mark was as follows: H3K27ac - 1581, H3K27me3 - 703, H3K36me3 - 719, H3K4me1 - 985, H3K4me3 - 803, H3K9me3 - 682 and included the 134 ChIP-seq datasets that were later excluded from other analyses for QC reasons, which included lack of agreement between observed and imputed data (see below). In addition, for epigenomes with two or more ChIP-seq experiments, we imputed data for each mark and epigenome combination for which we had observed data. For these imputations, we did not use information from the observed data for the combination being imputed. In total, the number of imputed datasets for each histone modification that we generated was: H3K27ac - 1088; H3K27me3 - 1703; H3K36me3 - 1703; H3K4me1 - 1688; H3K4me3 - 1688; H3K9me3 - 1700.

For those reference epigenome-mark combinations for which we had both observed and imputed data, we computed the agreement between observed and imputed data as QC (**Supplementary Table 10**). For this, we focused on two metrics. The first was the overlap percentage of the top 1% 25-bp intervals with the highest signal between observed and imputed data. The second was the Pearson correlation between observed and imputed data, where we set values above 500 to 500 so the metric would be more robust to outliers. The imputed ChIP-seq data for the six reference epigenomes with no observed ChIP-seq data after excluding observed ChIP-seq data for QC reasons (see above) were excluded from the release and any further analyses.

*ChIP-seq and chromatin state imputation from RNA-seq data*

For those reference epigenomes that had RNA-seq data available but not ChIP-seq data, we generated an auxiliary set of imputed ChIP-seq data. This is described in Fu and Ernst[5]. In brief, signal tracks were generated from gene-level quantification based on assigning intergenic positions a value of 0 and intragenic positions the expression value of an overlapping gene. The gene-level quantification was log2(TPM+1), and the assignment was done at 200 bp resolution. ChromImpute was applied as described above but imputed data was only generated for the 449 epigenomes with RNA-seq and without observed ChIP-seq data that passed QC. The flag '-p LOG2RNA' was used, which restricted the model to only using gene expression for making predictions. As described in Fu and Ernst[5], we also used the Gene Expression-based Chromatin State Imputation (GECSI) method to generate ChromHMM chromatin state predictions for the 449 epigenomes that had RNA-seq but no observed ChIP-seq data that passed QC.

*Peak calls on imputed ChIP-seq data*

We called both narrow and broad peaks using MACS2 (v.2.2.7.1)[6] on each imputed ChIP-seq data. To call narrowPeaks, we used the command `macs2 bdgpeakcall -c 2 --no-trackline` and to call broadpeaks, we used the command `macs2 bdgbroadcall -c 2 -C 1`. Any peaks overlapping the ENCODE excluded regions (hg38, accession: `ENCFF356LFX`)[7,8] were filtered out using the bedtools command `bedtools subtract -A`[9].

*DNA Methylation imputation*

We imputed WGBS DNA methylation for the samples with available ChIP-seq data using ChromImpute (v.1.0.5)[4]. For training we used methylation grids that had coverage of at least three reads. This imputation was done based on the 1704 reference epigenomes with ChIP-seq data for at least one histone modification. We followed procedures as above for imputing ChIP-seq data from other ChIP-seq data but included the '-dnamethyl' flag. As above, when imputing ChIP-seq data from other ChIP-seq data, we excluded imputed DNA methylation from the six reference epigenomes with no observed ChIP-seq after excluding observed

ChIP-seq data for QC reasons from the release and further analyses.

#### Chromatin state annotations

*ChromHMM annotations per reference epigenome*

We generated per-reference epigenome chromatin state annotations using ChromHMM (v.1.24)[10] for all 1698 reference epigenomes with at least one ChIP-seq experiment included in the integrative analysis. We applied ChromHMM's MakeSegmentation command to binarised data with the 18-state model from the Roadmap Epigenomics Consortium[11]. We used observed data when available; otherwise, we used imputed data from ChromImpute, as described above. Only chr1-22 and chrX were annotated. To generate binarised data for observed data, we used the BinarizedBam command of ChromHMM using aligned reads for a mark and its control as the input. We included the '-paired' flag for those datasets annotated as being based on paired-end reads. We used the default resolution (200 bp) and binarisation threshold of BinarizeBam. For imputed data, the imputed signal data was first converted from 25 bp to 200 bp resolution by averaging the imputed values in each 200 bp bin. The binarisation threshold for a 200 bp bin was set in a mark-specific manner so that the average fraction of bins receiving a present call, over all datasets for imputed data for the mark, matched that average fraction for all observed data.

*Universal chromatin state annotation*

We generated a universal chromatin state annotation based on a 100-state full-stack model by applying ChromHMM (v.1.24)[10] to all 5339 observed IHEC ChIP-seq datasets in EpiATLAS. In total, 1698 reference epigenomes and six histone modifications (H3K4me1, H3K4me3, H3K9ac, H3K27ac, H3K27me3 and H3K36me3) were represented in the input dataset. Using these data, we generated a single genome annotation – shared across all biospecimens– assigning one chromatin state to each genomic position. As input we used the same binarised data as was used for the observed data for the per reference epigenome ChromHMM annotations described above. To train the 100-state stacked model, we followed the approach described previously for learning a stacked ChromHMM model primarily on Roadmap Epigenomics data[12]. Briefly, each chromosome was divided into 1-Mb segments (with the terminal segment ≤1 Mb). During each iteration of the Baum–Welch algorithm, 300 segments were randomly sampled and used for training rather than using the entire genome in every iteration. This subsampling strategy improves computational efficiency while maintaining robust parameter estimation.

*Summary chromatin state annotations*

We generated summary 18-state ChromHMM annotations for groups of reference epigenomes, which we refer to as combinations. Combinations were defined using the metadata fields harmonized_sample_ontology_intermediate and harmonized_sample_disease_high from the harmonised metadata version 1.2 (https://github.com/IHEC/epiATLAS-metadata-harmonization). Summary chromatin state annotations were computed for combinations containing at least two reference epigenomes, yielding a total of 56 combinations. Combination names were constructed by appending a suffix to the harmonized_sample_ontology_intermediate term indicating disease status: H, D, or C, corresponding to healthy, disease, or cancer categories in harmonized_sample_disease_high. For example, t-cellH denotes the group of healthy T-cell reference epigenomes. For the 50 combinations with fewer than 100 reference epigenomes, summary annotations were generated with the CSREP method[13]. For the six combinations with ≥ 100 reference epigenomes, summary annotations were generated by simply taking the fraction at each genomic position of total reference epigenomes that are annotated as each state (e.g. if 60 out of 100 reference epigenomes are annotated as state 1_TssA at position *i*, then this position *i* is summarised as state 1_TssA with probability 0.6). The six such combinations were Brain Disease (brainD), Brain Healthy (brainH), Monocyte healthy (monocyteH), Neutrophil healthy (neutrophilH), t cell healthy (tH), and venous blood cancer (venous_bloodC).

*ChromGene based annotations*

We generated ChromGene[14] annotations for the 1698 IHEC reference epigenomes with at least one ChIP-seq experiment based on observed data. Here, the ChromGene model was trained using binarised ChIP-seq data for the six IHEC histone modifications: H3K4me3, H3K4me1, H3K27ac, H3K27me3, H3K36me3, and H3K9me3 for protein coding genes. We used the same binarisation procedure as for the per-epigenome ChromHMM annotations described above, and also used binarised observed data where available and otherwise used binarized imputed data. We used GENCODE v29 for the gene annotations and included 2 kb flanking regions as previously[14]. We used 12-mixture components in the ChromGene model as previously[14] (**Supplementary Table 3**). Here, we used two states per mixture component instead of three as used previously because of the reduced diversity of input marks available to the model. We ordered mixture components based on decreasing order of median expression level, and states in mixture components by decreasing enrichment at the transcription start sites (TSS).

*ChromScore and ChromScoreHMM annotations from ChromActivity*

We applied the ChromActivity framework to generate ChromScore and ChromScoreHMM annotations[15] for the 1698 IHEC epigenomes with at least one ChIP-seq experiment included in the integrative analysis. As input to ChromActivity, we used features derived from -log10(p-value) signal tracks, narrow peak calls and chromatin states. Signal and peak features were generated for the six IHEC histone modifications H3K27ac, H3K27me3, H3K36me3, H3K4me1, H3K4me3 and H3K9me3, using observed data when available and imputed data otherwise. Chromatin state features were obtained from the IHEC per-epigenome 18-state annotations described above. Individual experiment predictors were trained using the same functional testing dataset used in a previous application of ChromActivity[15] to produce an overall ChromScore. For the ChromScoreHMM model, we applied the previously trained model to the binarized top 2% predictions from each expert predictor, as described previously[15].

#### Methylation segmentation based Epilogos

The genome was binned into 200-bp bins, and each bin was assigned a methylation state (HMR, PMD, LMR, or UMR) across samples based on the overlap between the bin and the corresponding methylation segment. Unlike ChromHMM-based Epilogos[16], in which chromatin states are defined over fixed 200-bp intervals, methylation segmentation is based on CpG windows, meaning that some 200-bp bins may span multiple states. Such bins were assigned a "mixed" state. Genomic gaps, including telomeric and centromeric regions obtained from the UCSC Genome Browser, were assigned a "gap" state. Using these six state annotations (HMR, PMD, LMR, UMR, mixed, and gap) across 505 samples, Epilogos was run in single-group mode (-m single) with saliency level S1 (-s S1; standard Kullback–Leibler relative entropy). To visualise regions of interest, the Epilogos output scores were converted to 'qcat' format and plotted using pyGenomeTracks.

#### Analysis of DNA methylation and repeats

*Methylation Segmentation*

Methylation segmentation was performed using MethylSeekR tool (v.1.38)[17] to classify the genome into four distinct states: Highly Methylated Regions (HMRs), Partially Methylated Domains (PMDs), Low Methylated Regions (LMRs), and Unmethylated Regions (UMRs). The following parameters were used: a methylation level threshold at 0.5, a coverage cutoff at five reads per CpG, and a maximum FDR of 0.05, resulting in a threshold of at least four CpGs per LMR. Only samples with at least 10X overall mean CpG coverage were used.

*Analysis of low methylated regions*

To jointly analyse LMRs across all samples and cell types, we employed the ChromH3M method[18]. In brief, the genome was divided into 200 bp bins, with each bin assigned a value of 1 if it overlapped with an LMR and 0 otherwise. The resulting binarised matrix (genomic bins × samples) was then processed using ChromHMM[10] to identify combinatorial LMR signatures across samples using an 18-state model. Further analysis of State-17 (common LMRs) was conducted using UMAP, based on the average methylation of CpGs. We prepared an input matrix for the UMAP analysis by calculating the mean methylation for all CpGs within each bin annotated as common LMRs (state 17). Additionally, we merged all overlapping and adjoining 200 bp bins, and calculated the mean methylation for 12,166 common LMR (state 17 in the 18-state model) bins across 505 samples. We then conducted the UMAP analysis with a setting of 10 nearest neighbours and a minimum distance of 0.4.

*Categorisation and characterisation of individual CpG sites*

CpG sites with fewer than 300 missing values (NA) across the 645 biospecimens were retained for analysis. Sites with methylation levels greater than 80% were classified as methylated, whereas those below this threshold were classified as unmethylated. Based on the prevalence of methylation across biospecimens, CpG sites were further grouped into three categories: High, for sites methylated in more than 95% of biospecimens; Intermediate, for sites methylated in 25–95% of biospecimens; and Low, for sites methylated in fewer than 25% of biospecimens. CpG coordinates were then resized to 10 bp intervals. For each CpG site, the distance to the nearest TSS was computed using bedtools[9] closest against GENCODE[19] v29 gene annotations (all genes), reporting the signed distance relative to the TSS using the -D option. For illustrations, we only used sites that were found within 50 kb of a TSS. For the overlap with repeats, we counted the number of CpG sites (using the 10 bp window) that overlapped repeats for each CpG classification using bedtools *intersect -wa –wb* on the UCSC hg38 RepeatMasker track[20].

*Overlap of repeats with histone marks*

We used the histone ChIP-seq narrow peaks and resized them to 200 bp around their centre. We counted the number of peaks that overlapped with UCSC hg38 RepeatMasker[20] track using bedtools[9] intersect. When more than one repeat overlapped a peak, the largest overlap was kept. We established a random baseline for comparison. For each sample, we generated a simulated sample of 200 bp random regions matching the sample's distribution of distances to nearest genes. For each sample, the simulation was repeated 1000 times and we counted the number of times our observed count was higher than the simulated one for each repeat family. A repeat subfamily was identified as significantly over-represented (enriched) when the over-represented incidence was greater than 995/1000 ($p < 0.005$). Coverage percentages were measured as *peaks overlapping a repeat subfamily/total sample peak count*. The (Observed – Expected) metric was calculated by subtracting the expected coverage (mean across the 1000 simulations) from the observed coverage in the sample.

#### Universal chromatin state enrichment analyses

*Overlap with prior universal chromatin state annotation and mnemonics*

The state labels, colors, and ordering were automatically generated such that each of the 100 states of the universal annotation is associated with the state from the Vu and Ernst[13] model for which it has maximum fold enrichment. If multiple states were associated with the same state, letter suffixes were appended to the states and ordered alphabetically based on decreasing enrichment values for the state (e.g. two states from the EpATLAS annotation were associated with the state znf1 and were respectively named znf1_a and znf1_b based on decreasing fold-enrichment values for the znf1 state). We used annotations from Vu and Ernst[13] that had previously been lifted over from hg19 to hg38.

*Metadata analysis for universal chromatin states*

We characterized the specificity of each chromatin state from the universal chromatin state annotation to the various categories of epigenomes defined in the metadata (v.1.3). To do so, for each mark in a given state, we first tested if the emissions for the mark were higher than those for the other marks using a one-sided t-test. Then, for each mark with a nominally significant p-value ($p<0.05$), we performed the procedure listed below:

- Let *L* be an empty list
- For each metadata column *M**:
    - For each unique category *C* in the metadata column *M* with at least 5 epigenomes represented in the chromatin state model for the mark:
        - Use logistic regression to test if the mark has different emissions in epigenomes in *C* (labelled as 1) compared to the epigenomes not in *C* (labelled as 0) while using one-hot encoded sequencing consortium information (the project column) as a covariate.
        - Let *p* and *coef* be the p-value and coefficient for the logistic regression
        - If *coef* > 0 (indicating the mark has higher emission in *C*)
            - Add the tuple (*M, C, p*) to *L*

**A list of all columns used can be found in* **Supplementary Table 3**

By doing so, for each state-mark combination, we obtained a list of metadata categories showing higher emissions for the mark. Significant categories were defined using a Bonferroni correction as those with p <

0.05/n where n=73,028 is the total number of logistic regressions performed. For each state-mark combination, we output up to the ten most significant categories in **Supplementary Table 3**.

*Overlap with external genomic annotations*

We computed the EpiATLAS chromatin state enrichments for six external genomic annotations using the "OverlapEnrichment" command of ChromHMM[10] software, including CpG islands, included in the ChromHMM software by default and originally obtained from the UCSC genome browser, as well as transcription end sites (TES), TSS, 2 kb regions surrounding TSS (TSS 2 kb), exons, and gene bodies obtained from the GENCODE v29.

**Similarity of ChIP-seq data across biospecimens**

*Embedding of biospecimens based on binarised ChIP-seq data*

We generated a binarised matrix of the ChIP-seq data based on the presence of peaks at promoters and at putative regulatory regions based on the universal chromatin state annotation (uCREs). To define uCREs, we took all positions assigned to any of the following types of states: 'EnhWk', 'EnhA', 'TxEnh', 'BivProm', 'PromF', 'TSS', or 'Acet' (**Supplementary Table 11**). Directly neighbouring regions were merged, resulting in 829,862 regions. Promoter regions were defined as 1 kb upstream of the TSS for each gene, as defined in GENCODE v29 human genes. Promoter and uCREs were concatenated into a single bed-file. Using bedtools, that bed-file was overlapped individually with the narrow peak files for each ChIP-seq sample. This resulted in a binary peak presence-absence matrix, with one dimension equal to the number of promoters and uCREs, and the other dimension equal to the number of ChIP-seq datasets. An entry is 1 if the promoter or uCRE overlaps at least 1 bp with a peak in the ChIP-seq dataset.

For clustering the matrix, the scipy.cluster.hierarchy.linkage function (v.1.3.1) was used with average linkage and optimal leaf ordering using Euclidean distance. To compute the pairwise correlation distances between rows of the binary peak presence-absence matrix, numpy.corrcoef (v.1.26.4) was used. The correlation matrix was clustered using scipy.cluster.hierarchy.linkage with default parameters.

To construct 2D embeddings of the ChIP-seq datasets, principal component analysis was applied to the binary matrix, retaining the first 50 principal components (chosen by the elbow in the scree plot) using the sklearn library (v.1.3.1). Then, the tSNE class of sklearn was used to reduce the 50-dimensional PCA representation to a 2-dimensional representation (using n_components=2, random_state=1, and all other parameters default).

*Similarity of binarised ChIP-seq data across metadata categories*

To quantify the degree to which a particular metadata category was related to similarity between datasets, the following approach was taken: First, for each dataset (i.e., ChIP-seq represented by binary peak presence-absence vector), the distance to all other datasets was computed and sorted in order of increasing distance. This was done using Euclidean distance both for the full-length binary vectors and for the distances in the tSNE representation of the data. For each dataset and for a given label, a receiver operator characteristic curve (ROC curve) was computed by proceeding from nearest to farthest datasets and testing whether they had the same label value (counting as a true positive) or a different label value (counting as a true negative). The area under the ROC curve, AUROC, up to the point of 0.1 false positive rate (FPR) was measured for: the histone mark (6 values, H3K27ac, H3K27me3, etc.); the biospecimen label (57 values, B cell derived cell line, KMS-11, T cell, etc.); the project or consortium that generated the data (8 values, AMED-CREST, BLUEPRINT, CEEHRC, etc.); and the harmonized sample disease high term (3 values, Cancer, Disease, and Healthy/None).

*Genotype-aware variability of histone marks in breast epigenomes*

A subset of EpiATLAS breast references for which WGS was performed were selected (**Supplementary Table 12**). As described[21], WASP[22] was used to filter out aligned reads whose position shifted when aligned on individual-specific references versus hg38. Peaks were then called with ChIP-seq read alignments using MACS2[6] described above. To generate individual-specific references, accompanying WGS data from the samples was variant called on GATK (v.4.1.4.1)[23]. Duplicate reads were marked using GATK MarkDuplicatesSpark, and base quality score recalibration was performed with GATK BaseRecalibrator and ApplyBQSR using dbSNP SNPs[24], 1000 Genomes Project Phase 3 indels[25], and Mills indels[26] as gold standards. GATK HaplotypeCaller was used to call variants, and SNVs were selected and filtered per GATK recommendations for hard-filtering germline short variants. Variants were then phased using WhatsHap[27]. Scripts used for variant calling and downstream steps can be found in the following repository: https://github.com/hauduc/cis-hbtl.

Peaks were called for all reference bias-robust ChIP-seq alignments for all six histone marks, and then binarized into 200 bp bins. ChromHMM (v.1.21) MakeSegmentation was then run on the binarized files using the EpiATLAS pre-trained six-mark model (model_18_core_K27ac). In the resulting segmentations, for each of the 18 state genomic ranges, Jaccard indices were computed for 1) pairs of samples of the same cell type and different individuals, and 2) pairs of samples of different cell types and different individuals.

**Metadata prediction with EpiClass**

The training and performance characterization of the EpiClass tool (v.1.0) are fully described in a companion article[28]. Briefly, a series of eight metadata classifiers based on a Multilayer Perceptron (MLP) model were trained on the EpiATLAS data and their classification performances evaluated through a stratified 10-fold cross-validation approach. The total training set corresponds to 20,922 BigWig signal files (comprising multiple normalisation outputs per ChIP-seq dataset (raw, fold change and p-value) and strand-specific files for RNA-seq and WGBS assays) from 7464 independent experiments (datasets) conducted on 2216 biological samples within the EpiATLAS. The biospecimen classifier was restricted to the 16 most common categories of metadata, representing ~2/3 of the epigenomes, corresponding to cell and tissue types as captured by the harmonized_sample_ontology_intermediate field. For each BigWig file, the signal was averaged in genomic bins to generate vector representations. Several different sets of genomic bins (features) were explored, including: 303,114 highly variable DNA methylation segments of 200 bp[29], 303,114 uCREs showing highest absolute spearman correlation coefficient between H3K27ac level and gene expression (average length ~2.3 kb, 19,864 protein coding genes (average length ~68 kb), and 30,321 non-overlapping 100 kb bins covering the whole genome. The most predictive 100 kb regions of the genome were identified through a SHapley Additive exPlanations (SHAP)[30] analysis for each biospecimen category, the selection process of regions being described in Raby et al.[28]. The protein-coding gene list they each contain was then submitted to g:Profiler with background adjusted to only protein-coding genes[31] and the top 3 enriched terms shown for specific biospecimens. The averages of the max ChromScore (from ChromActivity, see above) from the most predictive 100 kb regions over all files per biospecimen were also computed and compared to the equivalent values over the 30,321 regions of the genome for the same files. Statistical significance was assessed using a two-sided Welch's t-test.

For the application of EpiClass to public data (**Fig. 4h**), we retained only predictions from classifiers that achieved >80% label concordance on their known-label subset, with the prediction-score threshold for the "high-confidence" category chosen per metadata attribute and per data source to meet this concordance criterion. Applying these thresholds, we annotated 28,312 cancer status labels (ChIP-Atlas), 125,146 donor sex labels (ChIP-Atlas and Recount3), and 166,924 biomaterial type labels (ChIP-Atlas and Recount3), for a total of over 320,000 newly annotated metadata attributes. Averaged across the three attributes, 85% of the files missing the corresponding label in the included datasets were assigned a prediction.

**Integration of the EpiATLAS with single-cell data**

Single-cell RNA-seq and single-cell ATAC-seq data was obtained from Granja et al 2019[32]. Fragment files and gene expression matrices were downloaded from GEO (GSE139369). Single-cell ATAC-seq data was reprocessed on genome assembly hg38 using CellRangerATAC (v.1.1.0) and ArchR (v.1.0.2)[33]. Tn5 insertions were counted in all uCRE regions (**Supplementary Table 11**). ArchR's iterativeLSI method was run on this region set to obtain Principal Component and UMAP embeddings (binarising the signal, using 2 iterations, 25,000 variable features and 30 PCs). Single-cell RNA-seq data was reprocessed using Seurat (v.5.0.1)[34]. The scRNA PCA and UMAP embeddings were created using 50 Principal Components based on the 2000 most variable genes (selected by the VST method).

From IHEC EpiATLAS, samples annotated with hematopoietic cell type labels were combined into feature matrices (see **Supplementary Table 4, 5**). For RNA-seq data, corresponding FPKM values were aggregated across all samples in annotated genes (GENCODE v29). To obtain EpiATLAS reference embeddings for transcriptome data, we used Seurat's workflow for processing single-cell data, involving log-normalisation, identification of variable genes using VST transformation, and PCA with subsequent UMAP dimensionality reduction. To obtain reference embeddings for each histone modification, a binary region x sample matrix was created by overlapping called peaks with the uCREs. Methylation levels for uCRE regions were aggregated by computing aggregate methylation levels for each uCRE by dividing the number of unconverted cytosines at CpG sites in the region by the total coverage of these sites from the bulk WGBS methylation calling results (GemBS pipeline[35]). The resulting region x sample matrix was binarised using a threshold of 0.7 for creating a reference embedding. The binary matrices for putatively repressive marks (H3K27me3, H3K9me3 and DNA methylation) were inverted. Dimensionality reduction was applied to the binary matrices with the 50,000 most variable regions (based on non-binarized signal) selected as features. The matrix was transformed using the TF-IDF transformation and principal components/singular values were computed as reduced dimensions. Based on these reduced dimensions (top 50 PCs), UMAP embeddings were created and Lovain clustering was performed.

We used the aggregated feature matrices to project single-cell transcriptome profiles into EpiATLAS reference embeddings generated from the same feature set. For this task, we used Seurat's FindTransferAnchors and MapQuery functions using the aggregated EpiATLAS object as reference and the single-cell object as query, using 30 PCs. To project scATAC-seq profiles into the ChIP-seq reference embeddings, we extracted the submatrix corresponding to features selected for the EpiATLAS reference embedding method. To project scATAC-data into EpiATLAS reference epigenome embeddings, we applied the same transformations and feature selection to the single-cell data that was used to generate the reference embedding. Data was then projected onto the reference embedding using matrix multiplication on the scaled TF-IDF transformed, binarized single-cell counts to obtain PC embeddings and subsequently applying the reference UMAP model (using the umap_transform() function of the uwot package).

We then predicted the cell type annotated in EpiATLAS annotated cell type (`harmonized_sample_label` in the metadata) from the embedded scRNA-seq and scATAC-seq data[32] by training a linear discriminant analysis (LDA) model using the reference principal component embeddings as features. These models were then used to predict the EpiATLAS class labels for the embedded single cells. Agreement between the harmonized EpiATLAS cell types ('harmonized_sample_label' metadata) and the single-cell cluster annotation[32] was computed using the Adjusted Rand Index (ARI).

For the reverse projection of EpiATLAS samples into single-cell reference embeddings (**Supplementary Fig. 5**), the same binarisation procedure described above was used to generate uCRE region × sample matrices. The resulting binary matrices for all histone marks were then transformed using the TF–IDF normalization parameters derived from the single-cell reference dataset. The normalised matrices were projected into the reference principal component (PC) space by matrix multiplication with the PC loading vectors computed from the single-cell reference. UMAP coordinates were subsequently obtained from the projected PCA embeddings using the UMAP model trained on the single-cell data.

WGBS data was projected in a similar fashion: the WGBS methylation level matrix by sample and uCRE region was binarised into "unmethylated" (binary value 1) and "methylated" (binary value 0) using a threshold methylation level of 0.7, encoding the globally anticorrelation of methylation levels and chromatin accessibility. The remaining steps were the same as for the histone marks. We then predicted the cell type annotated in the scRNA-seq and scATAC-seq data for the projected EpiATLAS samples by training a Linear Discriminant Analysis (LDA) model for single-cell class labels using the single-cell principal component embeddings as features. These models were then used to predict the class labels for the embedded EpiATLAS samples. Agreement between the harmonised EpiATLAS cell types ('sample_harmonized_cell_type' metadata column) was computed using the Adjusted Rand Index (ARI).

**Interactions between potential regulatory elements and genes**

*Predicting interactions with the generalized ABC-score*

As a set of putative regulatory regions, we used the uCREs derived from the universal ChromHMM annotation (**Supplementary Table 11**). STARE (v.1.0.4)[36] was used to compute biospecimen-specific uCRE-gene interactions for all biospecimens where observed H3K27ac ChIP-seq data were available (n = 1565). The H3K27ac signal (fold-change over input) was used as uCRE activity. To reduce noise in the ChromHMM regions, which are large (average 1235 bp) in comparison to sample-specific H3K27ac ChIP-seq peaks (average 650 bp), we set their activity to zero if no biospecimen-specific narrow H3K27ac ChIP-seq peak overlapped the ChromHMM region. As contact data, an average Hi-C matrix derived from 35 biosamples was taken[37]. The window size was set to 5 Mb, the GENCODE v38 annotation was used[19], regions overlapping ENCODE excluded regions (v.2) were removed[7], and the gABC-cutoff was set to 0.02. Besides using the regulatory regions defined by the universal ChromHMM annotation as candidate CREs, we generated a separate set of predicted interactions, by scoring the CRE-gene pairs for each biospecimen individually based on the H3K27ac ChIP-seq peaks. MACS2's signalValue was used as CRE activity; all other parameters were kept the same. Processing of genomic coordinates was done with pybedtools (v.0.9.0)[38]. To calculate enrichment of disease genes among genes with a low variability of interactions, all genes annotated for any disease from DisGeNET were used (v.24.3, https://www.disgenet.com)[39]. Genes were then sorted ascendingly by their standard deviation of the number of interactions across biospecimens and the enrichment of disease genes within the ranked list was calculated with the prerank function of GSEApy (v.1.1.2)[40]. The number of permutations was set to 1000 and the weight of the sorting metric to zero. To get the enrichment of ubiquitously expressed genes, all genes with an expression of TPM ≥ 0.5 in at least 90% of all biospecimens were considered to be ubiquitous. Then, GSEApy was used in the same way as for the disease genes.

*Correlation of signal in uCRE-gene pairs*

We calculated the Spearman correlation between the uCRE's activity (mean H3K27ac fold-change over input) and gene expression (TPM) for all possible uCRE-gene pairs with ≤ 250 kb distance (n = 7,865,786) across all biospecimens where observed H3K27ac ChIP-seq and RNA-seq were available (n = 965). If multiple RNA-seq measurements were available for the same EpiRR ID, total RNA-seq was chosen. Interactions where the uCRE overlapped the promoter of the gene (± 200 bp around any annotated TSS) were not included. To reduce the influence of gene co-expression on the derived correlations, we calculated the partial correlation of the interactions by controlling for the expression of all other genes within 250 kb distance (G) (pingouin v.0.5.4):

$$partial\ \rho(H3K27ac_{CRE}, RNA_g | RNA_G) .$$

For a null model, we followed Xie et al.[41] and randomly rewired the uCRE-gene pairs while maintaining the total number of interactions and interactions per gene and uCRE. The rewiring was not restricted to intrachromosomal interactions. The p-value for the partial correlation of an interaction was then taken from the empirical cumulative distribution function of the null distribution (statsmodels v.0.14.0) and subsequently adjusted for multiple testing (Benjamini-Hochberg).

*Comparison with CRISPRi and eQTL data*

To compare different metrics for scoring uCRE-gene pairs, we used the CRISPRi-interactions from Gschwind et al.[37], more specifically, the file EPCrisprBenchmark_ensemble_data_GRCh38.tsv.gz from their GitHub (https://github.com/EngreitzLab/CRISPR_comparison/tree/main/resources/crispr_data) and the column "Significant" to identify true interactions. To include the gABC-score in the comparison, the interaction scores based on the uCREs for the K562 biospecimen (IHECRE00001887) were used. Here, no threshold was set on the gABC-score.

As eQTL-gene pairs, we took the data that was fine-mapped with DAP-G from GTEx[42] (dbGaP accession number phs000424.v8.p2). We matched GTEx tissues to IHEC biospecimens (**Supplementary Table 13**). To calculate the normalized enrichment score (NES), we used GSEApy (v.1.1.2)[40], while setting the number of permutations to 100 and the weight of the sorting metric to zero. To compare the interaction scoring metrics for a biospecimen, the same set of uCRE-gene pairs was ranked for each metric. For each IHEC biospecimen, only interactions where the uCRE overlapped a narrow H3K27ac ChIP-seq peak from that biospecimen were kept.

**Transcript expression prediction using networks and epigenetic data**

We used 6 histone modifications (H3K4me1, H3K4me3, H3K9me3, H3K27ac, H3K27me3 and H3K36me3) and DNA methylation to compute sample-specific feature values for each transcript by segmenting a transcript into 15 fixed-sized (400 bp) windows (5 features each upstream of TSS, downstream of TSS, and downstream of TES) and 6 variably-sized transcript body regions: first exon, first intron, internal exons, internal introns, last intron and last exon. For histone modifications, feature values for each region were then computed, defined as the log2 transformed average fold change over control across the region with a pseudo count of 1e-9 and shifted with a minimum value starting at 0. For DNA methylation, the average beta values were used. Missing values were substituted with -1, and in the case of DNA methylation, this specifically corresponds to a feature region having none of its bases with CpG coverage of at least five. TPM values of the mRNA-seq or total RNA-seq datasets were log2 transformed with a pseudocount of 1e-3, which served as the modelling target of the transcripts.

We used the epigenetic features defined above (21 features each for 6 histone marks and DNA methylation) from each sample to train their respective Graph Convolutional Neural Network (GCN) models to assess feature importance, as described in[43]. The GraphConv module of Deep Graph Library (DGL) (v.0.6.1) and PyTorch (v.1.8.1) was used. The graphs additionally included a physical protein-protein interaction (PPI) network from BioGRID (v3.4.162)[44] for gene-gene edges and 10 cell type-specific Hi-C chromatin interaction networks from GM12878, H1-hESC, HCT-116, HeLa, HepG2, HMEC, HUVEC, IMR-90, K562 and NHEK from 4DNucleome[45] and Rao and Huntley et al.[46] for gene-bin edges. We used 6 hidden graph convolutional layers with 128 neurons in each hidden layer, in addition to the input and output layers. Out of the overall 324 samples, 34 had both mRNA-seq and total RNA-seq data. For those, we built two separate models using transcript expression levels from the two individual datasets. This resulted in a total of 358 trained GCN models. The regression models were then applied to infer expression levels of the left-out transcripts in the testing set (transcripts in chromosomes 18 and 19), both in this sample and the remaining 357 samples, using their corresponding epigenomic features. To group samples by tissue to evaluate different inference cases, the column ‘tissue_type’ from metadata version 1.0 was used. We then computed the saliency as the feature importance score, defined as the gradient of the GCN output of a node with respect to a feature. We then normalised the feature importances across all features in each model and plotted the log10-transformed normalised importance across feature regions separately for each assay type. In addition, the overall importance of each type of epigenetic signal was aggregated by averaging feature importances across different regions.

**Analysis of epigenetic associations with alternative exon and intron inclusion**

Epigenomes where RNA-seq (mRNA-seq if available, otherwise total RNA-seq), observed DNAm, and all histone marks as either imputed or observed data were available were used (n=405). RNA-seq transcript expression files from RSEM served as input to SUPPA2 (v.2.3)[47] to identify splicing events on high-confidence transcripts, i.e, annotated transcript support levels one or two, and calculate their proportion spliced in (PSI) values. We filtered for skipped exons (SE) and retained introns (RI) on autosomes, which do not overlap with other events, to remove potential noise.

To assess epigenomic associations to exon/intron inclusion, we averaged the epigenetic signal in the SE/RI and its adjacent areas, i.e., the preceding and succeeding introns/exons. Additionally, for each event, we gathered the corresponding gene's expression value and intrinsic features, such as the regions' widths, their distance to the TSS and TES, and their splice site strength using MaxEntScan[48]. Since the overall gene expression is closely related to the PSI values as they are computed on the same data, we calculate the partial Pearson correlation coefficient between the PSI and the features mentioned above over all events for each biospecimen, controlling for gene expression measured by TPM.

**Estimation of metabolic networks from RNA expression data**

Mean log2 expression values (TPM) of Recon3D[49] metabolic genes were calculated for each of the 33 cell types (based on the metadata v1.2 category *harmonized_sample_ontology_intermediate*) with at least three samples with available gene expression (mRNA-seq or total RNA-seq), ChromGene annotations, and predicted enhancer activities (based on the gABC-interactions, averaged per cell type). A majority ChromGene annotation was assigned for each gene within each cell type if representing at least 50% of the samples, otherwise it was marked "undetermined". For each gene and sample, the activity (H3K27ac ChIP-seq fold-change over ChIP input) of all enhancers predicted with the previously described gABC model[36] to interact with the gene (based on uCREs, see above) was summed. The summed activity for genes without ≥ 2 H3K27ac signal at their promoter (±200 bp around any annotated TSS) was set to zero. The activity was subsequently averaged across samples of each cell type. Samples were ranked using the median number of reactions included in the metabolic reconstructions obtained with the RNA-seq-based rFASTCORMICS workflow[50] for each sample. Enrichment for transporters or cytochrome P450 enzymes of unique metabolic genes exhibiting cell type-specific active ChromGene annotations (annotations 1-6 in one cell type vs. 7-12 in all other cell types) against all metabolic genes of Recon3D was assessed with a hypergeometric test. For more details please refer to the respective companion paper[51].

*Transcription factor motif enrichment analysis in pathway-associated elements*

To identify upstream regulators of metabolic pathways in hepatocytes, we performed an overlap enrichment analysis between transcription factor motif occurrences and pathway-linked regulatory elements. We used the previously described set of sample-specific gABC interactions to identify sets of gene-associated regulatory elements. We paired these elements with gene metabolic pathway annotations to aggregate the set of pathway-associated regulatory elements for each pathway. Samples were then grouped by tissue type (e.g., hepatocyte), and pathway-associated elements were merged across all samples within each tissue category to create tissue-level summary sets of regulatory elements for each pathway. Next, we accessed genome-wide annotations of motif occurrences for 280 non-redundant archetypal transcription factor motif modules for the human reference genome[52]. These motif modules represent clusters of related TF binding patterns. We then calculated a tissue-specific binomial-based enrichment between pathway-associated elements and motif archetype occurrences using the *overlapEnrichments* tool from the Gonomics package (v.1.0.1)[53] using a merged superset of biospecimen-specific ChromHMM enhancer states (including EnhG1, EnhG2, EnhA1, EnhA2, and EnhWk) across all biosamples as background regions. TF-pathway associations with FDR-adjusted p-values below 0.01 were considered significant, using the more significant of enrichment or depletion p-values for FDR adjustment.

#### EpiATLAS as resource for disease studies

625 disease-related biospecimens were selected by filtering the *harmonized_sample_disease_high* column from metadata version 1.3 for entries labelled as *Disease* and *Cancer*.

*Tissue-specific genetic contributions to complex traits*

To investigate the tissue-stratified genetic contributions to complex traits across individuals, we applied polygenic score (PGS) partitioning using epigenomic annotations from EpiATLAS in combination with inclusive polygenic scores (iPGS) and a pathway-based PGS approach in PRSet[54](version 2021-09-20) using the UK Biobank resource[55,56]. We conducted the PGS analysis using the UK Biobank Resource under Application Number 21942, "Integrated models of complex traits in the UK Biobank" (https://www.ukbiobank.ac.uk/enable-your-research/approved-research/integrated-models-of-complex-traits-in-the-uk-biobank). All participants of the UK Biobank resource provided written informed consent. More information is available at https://www.ukbiobank.ac.uk/explore-your-participation/basis-of-your-participation/.

We utilized the iPGS models trained on $n_{train}$ = 284,661 individuals with hyperparameter tuning conducted on

$n_{validation}$ = 40,667 individuals, as previously described[55]. We focused on the iPGS model for standing height. We evaluated the biosample enrichment and individual-level partitioned PGS on a held-out test set of $n_{test}$ = 67,730 white British individuals from the UK Biobank resource.

For epigenomic annotations, we used enhancer regions defined by summary ChromHMM annotations from CSREP files derived from healthy biosamples in EpiATLAS. We used the following ChromHMM states to define enhancer regions: 7_EnhG1, 8_EnhG2, 9_EnhA1, 10_EnhA2, and 11_EnhWk.

Using the PRSet[54] framework, we computed the stratified PGS for each biosample group by summing the weighted genotype dosages for variants located within the annotated enhancer regions. We quantified the biosample-specific enrichment by calculating the ratio of the variance explained by the stratified PGS relative to the fraction of the genome covered by the biosample's enhancer regions, as in the enrichment calculation in PRSet.

The stratified PGS model provides individual-level polygenic scores partitioned by biosample group. It enables the decomposition of each individual's total polygenic score into biosample-annotated components, reflecting the tissues most likely to mediate genetic liability. We assessed interindividual variability in the dominant biosample contributions and visualized biosample-specific polygenic profiles across the white British individuals in the held-out test set ($n_{test}$ = 67,730).

To analyse the IHEC EpiATLAS histone mark dataset against the entire GWAS catalog[57] (version Dec-19-2021), we utilized FORGE2[58], command line version, default settings: 1000 background repetitions, LD filtering at r2=0.8, default adjustment for Minor Allele Frequency, TSS distance and GC content. FORGE2 analysis was applied with this expanded atlas of histone mark broad peaks, using the Benjamini-Hochberg procedure to control the false discovery rate when identifying tissue- and cell type-specific associations.

*Heritability enrichment of expressed genes for multiple sclerosis susceptibility*

To detect cell/tissue-specific expression, healthy samples were selected and grouped according to EpiATLAS ontology intermediate classifications (metadata version 1.3). Further sub-grouping applied to blood/immune samples was based on cell- and/or tissue type classifications. Harmonised RNA-seq data from all samples processed through the grape-nf pipeline were filtered using the filterByExpr() function from the edgeR package (v.3.32.1)[59] with default settings and subsequently normalised using the calcNormFactors() function. Differential expression analysis was conducted using Limma (v.3.46.0) and Voom[60] with a minimum of 3 samples per group and similar sample types excluded from the contrast. T-statistics from each contrast were ranked and the top 10% were considered cell/tissue-specific genes as previously described[61]. Gene coordinates were extracted using biomaRt (v.2.46.3)[62]. Annotation files were generated using the make_annot.py script from the LDSC tool (v.2.0.1)[63] with a window size of 1Mb and subsequently LD scores were computed using the ldsc.py script. The GWAS summary statistics for MS susceptibility[64] were filtered for HapMap3 SNPs using the munge_sumstats.py script[65]. Heritability enrichment of specifically expressed genes, a.k.a. LDSC-SEG, was estimated using the ldsc.py script with the --h2-cts flag and baselineLD (v.2.2). For visualization purposes, the -log10(Coefficient_P-value), derived from output cell_type_results.txt files, was plotted according to EpiATLAS ontology intermediate classifications and blood/immune cell/tissue subtypes. Polygenic risk scores (PRS) for MS susceptibility were previously reported[66].

*Repetitive elements in patients with Chronic Inflammatory Diseases (CID).*

For quantification and differential analysis of repetitive elements in CID, the Subfamily Assignment for Multiple Alignment (SAMA) pipeline[67] was used for the alignment of repetitive elements in the SYSCID dataset. Briefly, paired-end RNA-seq sequencing reads from each individual were mapped to the GRCh38/hg38 reference genome assembly using STAR (v.2.7.10b)[68] with the parameter --outFilterMultimapNmax 150. Raw read counts for the annotated genes and repetitive elements were generated using the R package

GenomicAlignments summarizeOverlaps function[69], referencing the GENCODE v43 transcriptome assembly and the UCSC RepeatMasker annotated repeats, respectively. The raw read count matrix of repetitive elements across samples was normalized by the size factor vector estimated from the corresponding gene counts and then used for differential analysis with DESeq2 (v.1.38.3)[70]. For track visualization, Reads Per Million (RPM) signals of uniquely aligned reads were utilized to calculate the mean expression value and standard errors across all healthy controls and patients in the disease groups.

*Chromatin states at the CDKN2A locus*

Chromatin states were taken from the ChromHMM 18-state model segmentation of the EpiATLAS collection of harmonised cancer reference epigenomes. We compared healthy B cells and their malignant counterparts, selecting five random samples per category (defined by B-cell differentiation stage for healthy cells and by disease subtype for malignant cells) when more than five were available. The chromatin landscape of the 9p21.3 locus (chr9:21,942,375-22,141,264, hg38) was visualised using the R package karyoploteR (v.1.32.0)[71].
