## Extended and Supplementary Figures for "EpiATLAS – a reference for human epigenomic research"

**EpiATLAS Extended and Supplementary Figures**

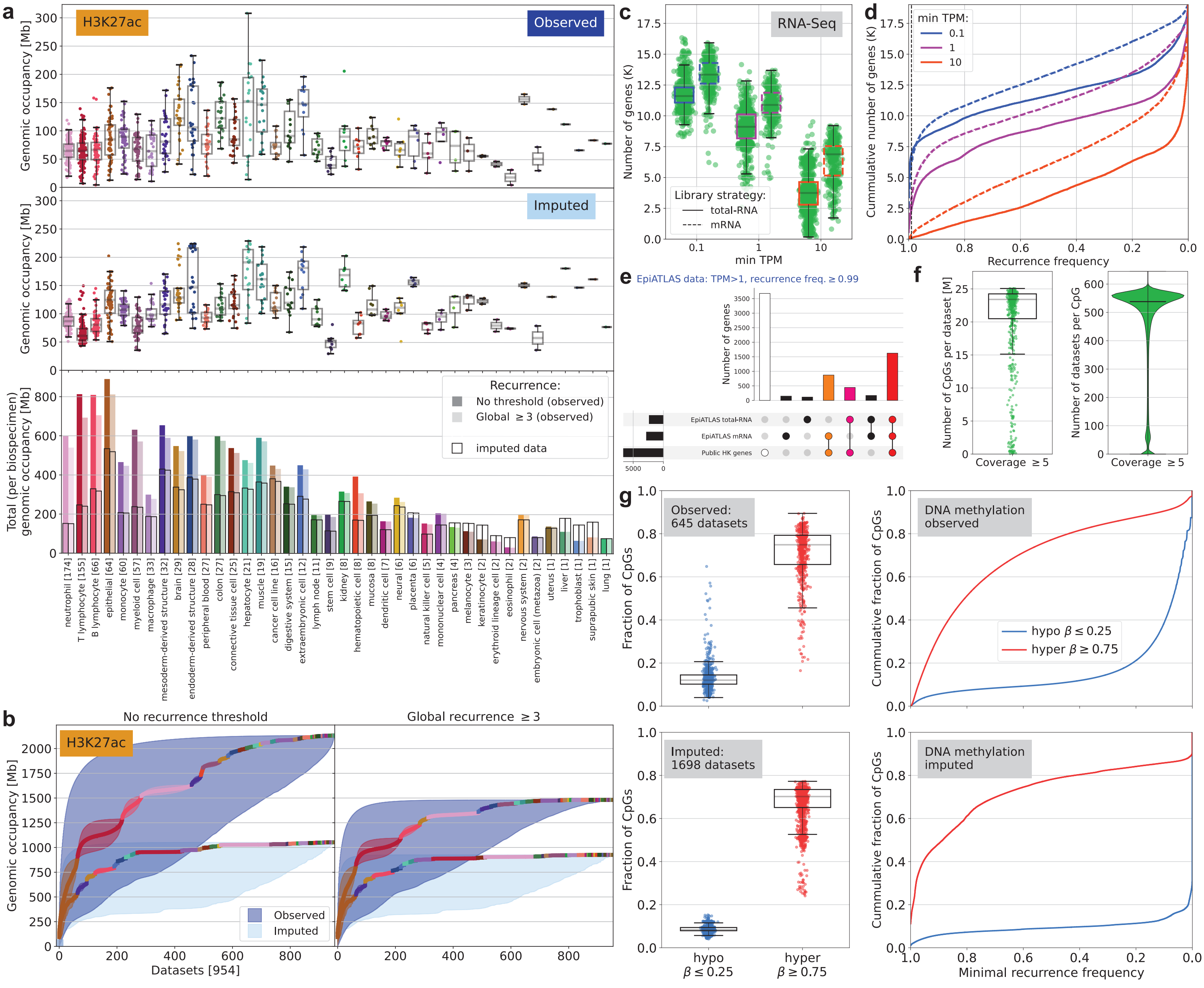

a
H3K27ac
Observed
Imputed
Genomic occupancy [Mb]
Total (per biospecimen) genomic occupancy [Mb]
Recurrence:
No threshold (observed)
Global ≥ 3 (observed)
imputed data
neutrophil [174]
T lymphocyte [155]
B lymphocyte [66]
epithelial [64]
monocyte [60]
myeloid cell [57]
macrophage [33]
mesoderm-derived structure [32]
brain [29]
endoderm-derived structure [28]
peripheral blood [27]
colon [27]
connective tissue cell [25]
hepatocyte [21]
muscle [19]
cancer cell line [16]
digestive system [15]
extraembryonic cell [12]
lymph node [11]
stem cell [9]
kidney [8]
hematopoietic cell [8]
mucosa [8]
dendritic cell [7]
neural [6]
placenta [6]
natural killer cell [5]
mononuclear cell [4]
pancreas [4]
melanocyte [3]
keratinocyte [2]
erythroid lineage cell [2]
eosinophil [2]
nervous system [2]
embryonic cell (metazoa) [2]
uterus [1]
liver [1]
trophoblast [1]
suprapubic skin [1]
lung [1]
b
No recurrence threshold
Global recurrence ≥ 3
H3K27ac
Genomic occupancy [Mb]
Datasets [954]
Observed
Imputed
c
RNA-Seq
Number of genes (K)
Library strategy:
total-RNA
mRNA
min TPM
d
min TPM:
0.1
1
10
Cummulative number of genes (K)
Recurrence frequency
e
EpiATLAS data: TPM>1, recurrence freq. ≥ 0.99
Number of genes
EpiATLAS total-RNA
EpiATLAS mRNA
Public HK genes
f
Number of CpGs per dataset [M]
Coverage ≥ 5
Number of datasets per CpG
Coverage ≥ 5
g
Observed:
645 datasets
Imputed:
1698 datasets
Fraction of CpGs
hypo
β ≤ 0.25
hyper
β ≥ 0.75
DNA methylation
observed
DNA methylation
imputed
Cummulative fraction of CpGs
hypo β ≤ 0.25
hyper β ≥ 0.75
Minimal recurrence frequency

**Extended Data Figure 1 | a,** Distributions of genomic enrichments per H3K27ac sample for observed (top) and imputed (middle) data. Bottom barplot shows genomic occupancy for each biospecimen source, given by the union of all samples with and without a recurrence threshold. The recurrence threshold is calculated globally using all datasets regardless of their biospecimen source. The black outline shows the same for imputed data. The enriched regions on autosomes only are summarised. **b,** Saturation dynamics without (left) and with (right) recurrence threshold of three**.** Middle curves are coloured and grouped by biospecimen source, and groups are sorted to reach the fastest saturation. Dark blue shows all possible saturation paths, independent of biospecimen source, for the observed data; light blue is for imputed data. **c,** Distributions of the number of discovered genes for three TPM thresholds for total-RNA and mRNA. **d,** Gene discovery recurrence across EpiATLAS for total-RNA (solid) and mRNA (dashed). It drops significantly with increasing TPM threshold, indicating that many highly expressed genes are tissue-specific. **e,** Highly recurrent genes, 2811 and 2352 genes for mRNA and total-RNA, respectively, using recurrence frequency ≥ 0.99 for TPM threshold=1, significantly overlap (hypergeometric p-value<10^-16) published lists of housekeeping (HK) genes [1,2]. **f,** Summary of CpG coverage for WGBS datasets. **g,** Left plots distributions of fractions hypo- and hypermethylated CpGs over EpiATLAS DNA methylation data. Imputed data (bottom plot) shows a much tighter distribution for hypomethylated CpGs. Two panels on the right are recurrence frequency plots showing similar patterns for both observed and imputed data. $\beta$: fraction of methylated reads at CpG

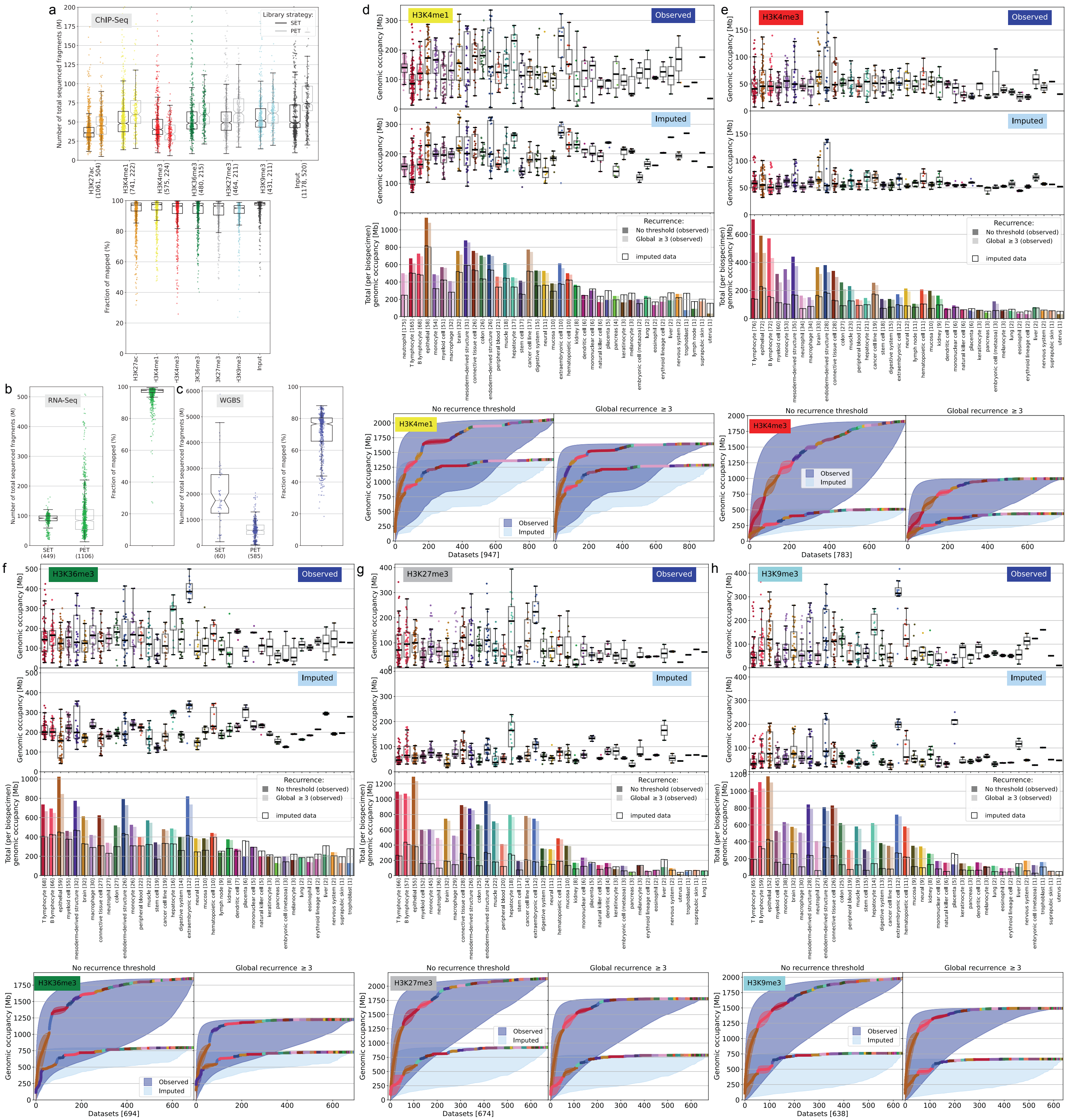
a
ChIP-Seq
Library strategy:
SET
PET
Number of total sequenced fragments (M)
H3K27ac (1061, 504)
H3K4me1 (741, 222)
H3K4me3 (575, 224)
H3K36me3 (480, 215)
H3K27me3 (464, 211)
H3K9me3 (431, 211)
Input (1178, 520)
Fraction of mapped (%)
b
RNA-Seq
SET (449)
PET (1106)
c
WGBS
SET (60)
PET (585)
d
H3K4me1
Observed
Imputed
Genomic occupancy [Mb]
Total (per biospecimen) genomic occupancy [Mb]
Recurrence:
No threshold (observed)
Global ≥ 3 (observed)
imputed data
No recurrence threshold
Global recurrence ≥ 3
Datasets [947]
e
H3K4me3
Datasets [783]
f
H3K36me3
Datasets [694]
g
H3K27me3
Datasets [674]
h
H3K9me3
Datasets [638]

**Supplementary Figure 1 |** Sequencing statistics for EpiATLAS data. **a,** Summary of the number of sequenced fragments for ChIP-seq. Top: distributions of total number of fragments per dataset; single-end (SET) and paired-end (PET) data shown separately (for pair-end data total number of sequenced reads is twice the number of fragments); x-axis labels show split between single-end and pair-end data. Bottom: distributions of fractions of mapped reads; for all data types, median mapping rate>95%. **b, c,** same as **a,** with left analogous to top and right analogous to bottom for RNA-seq and DNA methylation (WGBS) experiments. **d,** Top: Distributions of genomic enrichments per H3K4me1 sample for observed (top) and imputed (middle) data. Bottom barplot shows genomic occupancy for each biospecimen source, given by the union of all samples with and without a recurrence threshold. The recurrence threshold is calculated globally using all datasets regardless of their biospecimen source. The black outline shows the same for imputed data. The enriched regions on autosomes only are summarised. Bottom: Saturation dynamics without (left) and with (right) recurrence threshold of three**.** Middle curves are coloured and grouped by biospecimen source, and groups are sorted to reach the fastest saturation. Dark blue shows all possible saturation paths, independent of biospecimen source, for the observed data; light blue is for imputed data. **e-h,** Same as **d,** but for H3K4me3, H3K36me3, H3K27me3, and H3K9me3.

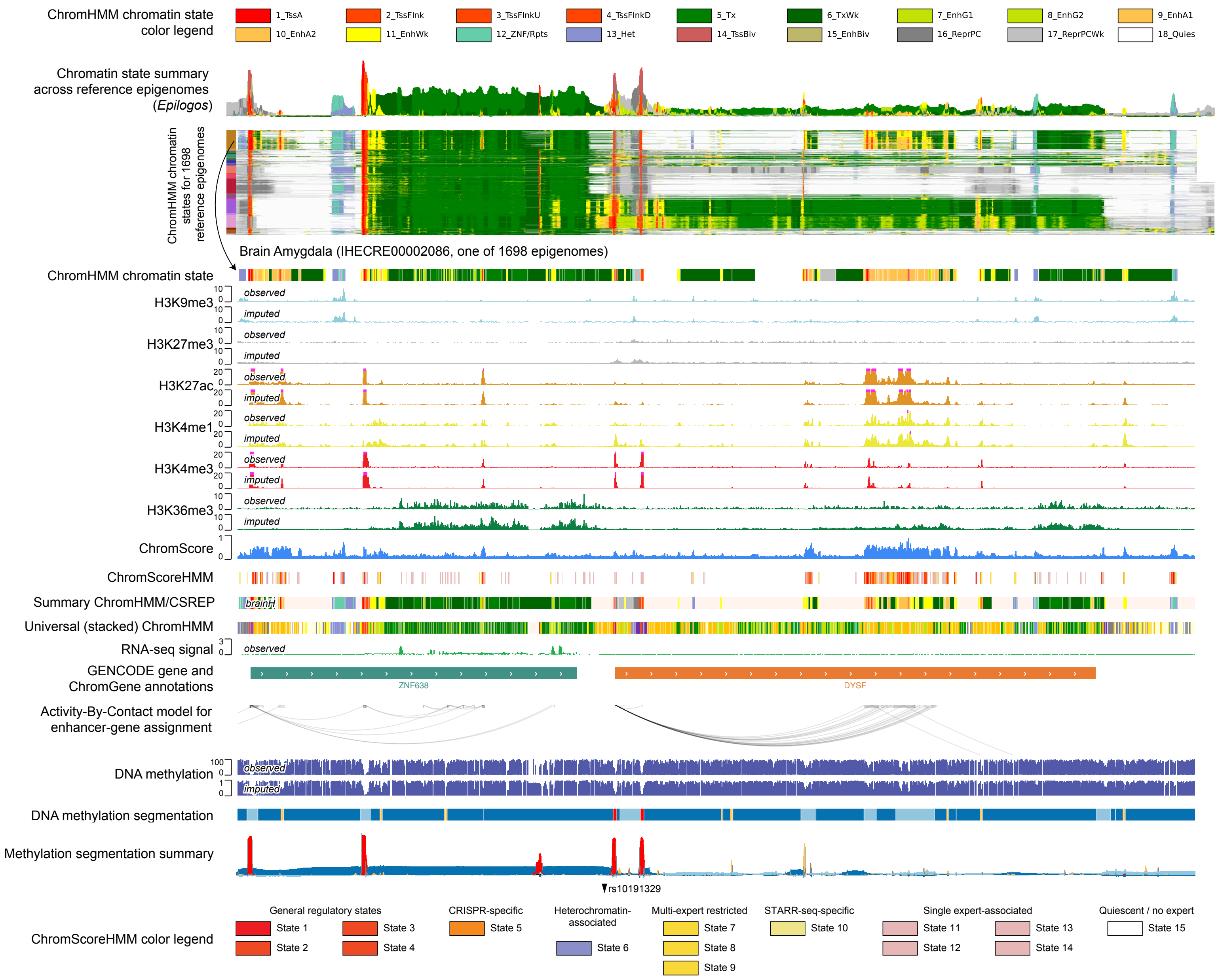
ChromHMM chromatin state color legend
1_TssA
2_TssFlnk
3_TssFlnkU
4_TssFlnkD
5_Tx
6_TxWk
7_EnhG1
8_EnhG2
9_EnhA1
10_EnhA2
11_EnhWk
12_ZNF/Rpts
13_Het
14_TssBiv
15_EnhBiv
16_ReprPC
17_ReprPCWk
18_Quies
Chromatin state summary across reference epigenomes (Epilogos)
ChromHMM chromatin states for 1698 reference epigenomes
Brain Amygdala (IHECRE00002086, one of 1698 epigenomes)
ChromHMM chromatin state
H3K9me3
H3K27me3
H3K27ac
H3K4me1
H3K4me3
H3K36me3
observed
imputed
ChromScore
ChromScoreHMM
Summary ChromHMM/CSREP
brainH
Universal (stacked) ChromHMM
RNA-seq signal
GENCODE gene and ChromGene annotations
ZNF638
DYSF
Activity-By-Contact model for enhancer-gene assignment
DNA methylation
DNA methylation segmentation
Methylation segmentation summary
rs10191329
ChromScoreHMM color legend
General regulatory states
State 1
State 2
State 3
State 4
CRISPR-specific
State 5
Heterochromatin-associated
State 6
Multi-expert restricted
State 7
State 8
State 9
STARR-seq-specific
State 10
Single expert-associated
State 11
State 12
State 13
State 14
Quiescent / no expert
State 15

**Extended Data Figure 2: Overview of epigenomic data across tissues and integrative annotations.** Similar to **Fig. 2**, but for another locus. The coordinates of the genomic region in hg38 are chr2:71270228-71735055. From top to bottom: (1) Colour legend for ChromHMM[3] chromatin state model for the 18-state model that was previously learned[4] and applied here. (2) Epilogos visualisation[5] of the 18-state ChromHMM annotations for 1698 reference epigenomes. The upper facet is the Epilogos information-theoretic visualisation summary, and below that are the individual epigenome chromatin state annotations. On the left is a colour bar corresponding to the biospecimen groups as in **Fig. 1C**. (3) ChromHMM chromatin state for a Brain Amygdala reference epigenome (IHECRE00002086), which is highlighted in individual tracks that follow. (4) The observed followed by the ChromImpute[6] imputed signal tracks for H3K9me3, H3K27me3, H3K27ac, H3K4me1, H3K4me3, H3K36me3 for epigenome IHECRE00002086. (5) The ChromScore and ChromScoreHMM tracks for epigenome IHECRE00002086. (6) A summary ChromHMM track of state assignments based on the 18-state model for epigenomes of the brain healthy group, which is one of 56 summary tracks corresponding to combinations of tissue/cell ontology and disease state. (7) A universal chromatin state annotation based on the stacked modelling approach[7] applied jointly to all EpiATLAS ChIP-seq data. The emission parameters and colour legend are shown in **Fig. 4a**. (8) Observed RNA-seq signal track for epigenome IHECRE00002086. (9) GENCODE gene annotations with protein coding genes colored by their ChromGene annotation[8] in epigenome IHECRE00002086. ChromGene emission parameters and colour legend can be found in **Supplementary Fig. 2a**. (10) An activity-by-contact model enhancer gene assignments for epigenome IHECRE00002086 from the gABC framework[9]. (11) Observed and ChromImpute imputed DNA methylation signal tracks. (12) Four-state methylation segmentation track from MethylSeekR[10] for epigenome IHECRE00002086. Corresponding colour legend can be found in **Fig. 3b**. (13) A summary track of methylation segmentations for 505 epigenomes, similar to the Epilogos summary track for chromatin states. (14) Genome-wide association study variant associated with severity of multiple sclerosis (rs10191329), which was enriched in the central nervous system[11]. (15) Colour legend for ChromScoreHMM states based on what was defined previously[12].

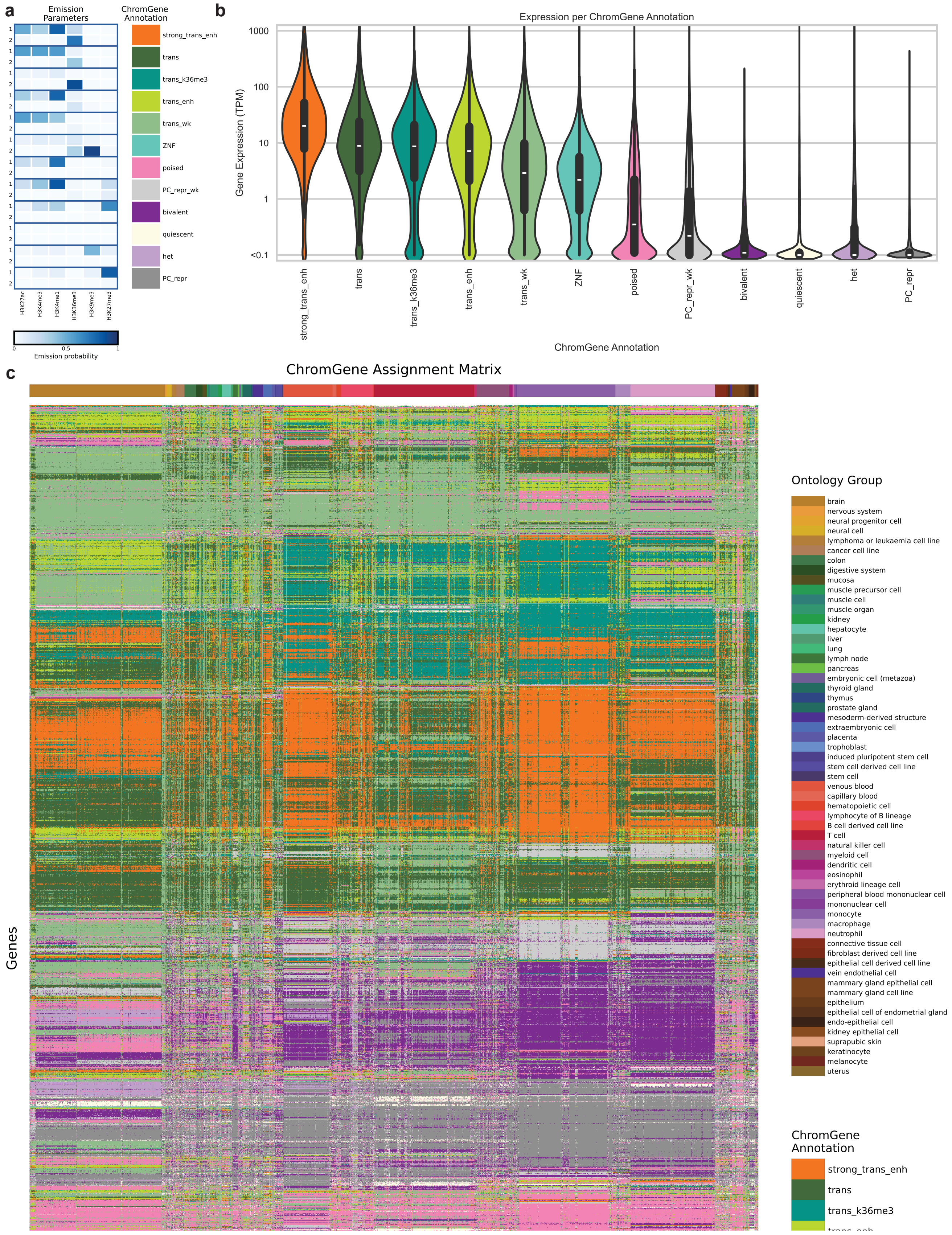
a
Emission Parameters
ChromGene Annotation
strong_trans_enh
trans
trans_k36me3
trans_enh
trans_wk
ZNF
poised
PC_repr_wk
bivalent
quiescent
het
PC_repr
H3K27ac
H3K4me3
H3K4me1
H3K36me3
H3K9me3
H3K27me3
0
0.5
1
Emission probability
b
Expression per ChromGene Annotation
Gene Expression (TPM)
1000
100
10
1
<0.1
strong_trans_enh
trans
trans_k36me3
trans_enh
trans_wk
ZNF
poised
PC_repr_wk
bivalent
quiescent
het
PC_repr
ChromGene Annotation
c
ChromGene Assignment Matrix
Genes
Ontology Group
brain
nervous system
neural progenitor cell
neural cell
lymphoma or leukaemia cell line
cancer cell line
colon
digestive system
mucosa
muscle precursor cell
muscle cell
muscle organ
kidney
hepatocyte
liver
lung
lymph node
pancreas
embryonic cell (metazoa)
thyroid gland
thymus
prostate gland
mesoderm-derived structure
extraembryonic cell
placenta
trophoblast
induced pluripotent stem cell
stem cell derived cell line
stem cell
venous blood
capillary blood
hematopoietic cell
lymphocyte of B lineage
B cell derived cell line
T cell
natural killer cell
myeloid cell
dendritic cell
eosinophil
erythroid lineage cell
peripheral blood mononuclear cell
mononuclear cell
monocyte
macrophage
neutrophil
connective tissue cell
fibroblast derived cell line
epithelial cell derived cell line
vein endothelial cell
mammary gland epithelial cell
mammary gland cell line
epithelium
epithelial cell of endometrial gland
endo-epithelial cell
kidney epithelial cell
suprapubic skin
keratinocyte
melanocyte
uterus
ChromGene Annotation
strong_trans_enh
trans
trans_k36me3

**Supplementary Figure 2 | a,** Emission parameters of ChromGene model. Each pair of two rows corresponds to a mixture component and each individual row corresponds to a state of the mixture component. The columns correspond to different IHEC histone modifications. Blue corresponds to higher emission values and white to lower values. The mixture components are given color and mnemonic labels as indicated on the right. Descriptions of abbreviations: trans: Transcribed; enh: Enhancer; het: heterochromatin; k36me3: H3K36me3; PC_repr: Polycomb repressed; wk: weak; ZNF: zinc finger proteins **b,** Gene expression distribution based on ChromGene annotation. Shown is the distribution of gene expression values in TPM for each ChromGene over all combinations of gene and reference epigenome. The y-axis is on a log scale. **c,** Representation of ChromGene assignment matrix. Rows correspond to 2000 randomly sampled genes and columns to the reference epigenomes. The assignments are indicated based on the color legend on the lower right. The top indicates the ontology group of the reference epigenome based on the color legend on the upper right.

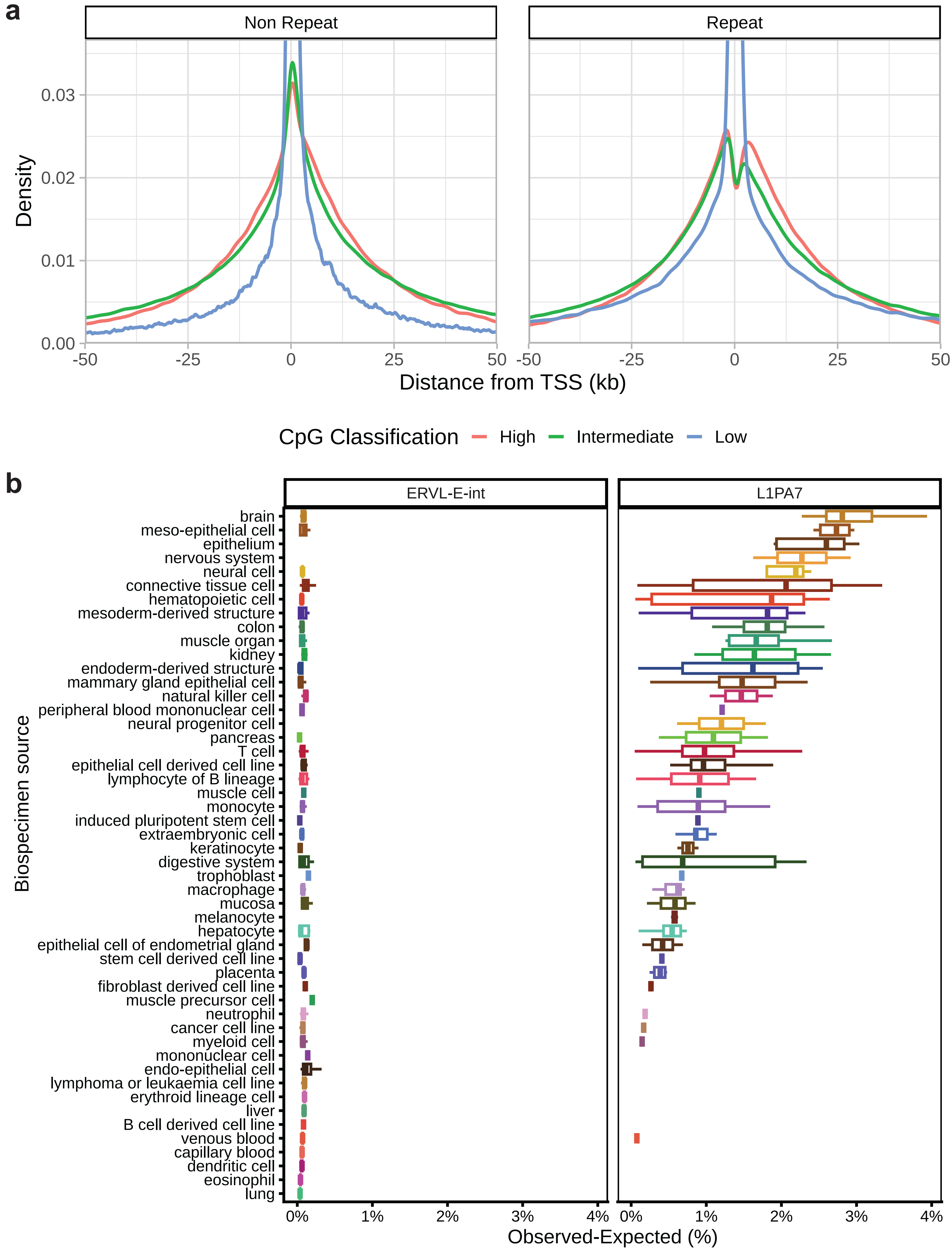
a
Non Repeat
Repeat
Density
0.00
0.01
0.02
0.03
-50
-25
0
25
50
Distance from TSS (kb)
CpG Classification
High
Intermediate
Low
b
ERVL-E-int
L1PA7
Biospecimen source
brain
meso-epithelial cell
epithelium
nervous system
neural cell
connective tissue cell
hematopoietic cell
mesoderm-derived structure
colon
muscle organ
kidney
endoderm-derived structure
mammary gland epithelial cell
natural killer cell
peripheral blood mononuclear cell
neural progenitor cell
pancreas
T cell
epithelial cell derived cell line
lymphocyte of B lineage
muscle cell
monocyte
induced pluripotent stem cell
extraembryonic cell
keratinocyte
digestive system
trophoblast
macrophage
mucosa
melanocyte
hepatocyte
epithelial cell of endometrial gland
stem cell derived cell line
placenta
fibroblast derived cell line
muscle precursor cell
neutrophil
cancer cell line
myeloid cell
mononuclear cell
endo-epithelial cell
lymphoma or leukaemia cell line
erythroid lineage cell
liver
B cell derived cell line
venous blood
capillary blood
dendritic cell
eosinophil
lung
0%
1%
2%
3%
4%
Observed-Expected (%)

**Supplementary Figure 3 | a,** Distance to TSS of the CpG sites depending on their methylation classification, separated between repeat overlapping and non-repeat overlapping. **b,** Complete set of all biospecimen sources and their enrichment in ERVL-E-int in contrast to L1PA7 for H3K9me3.

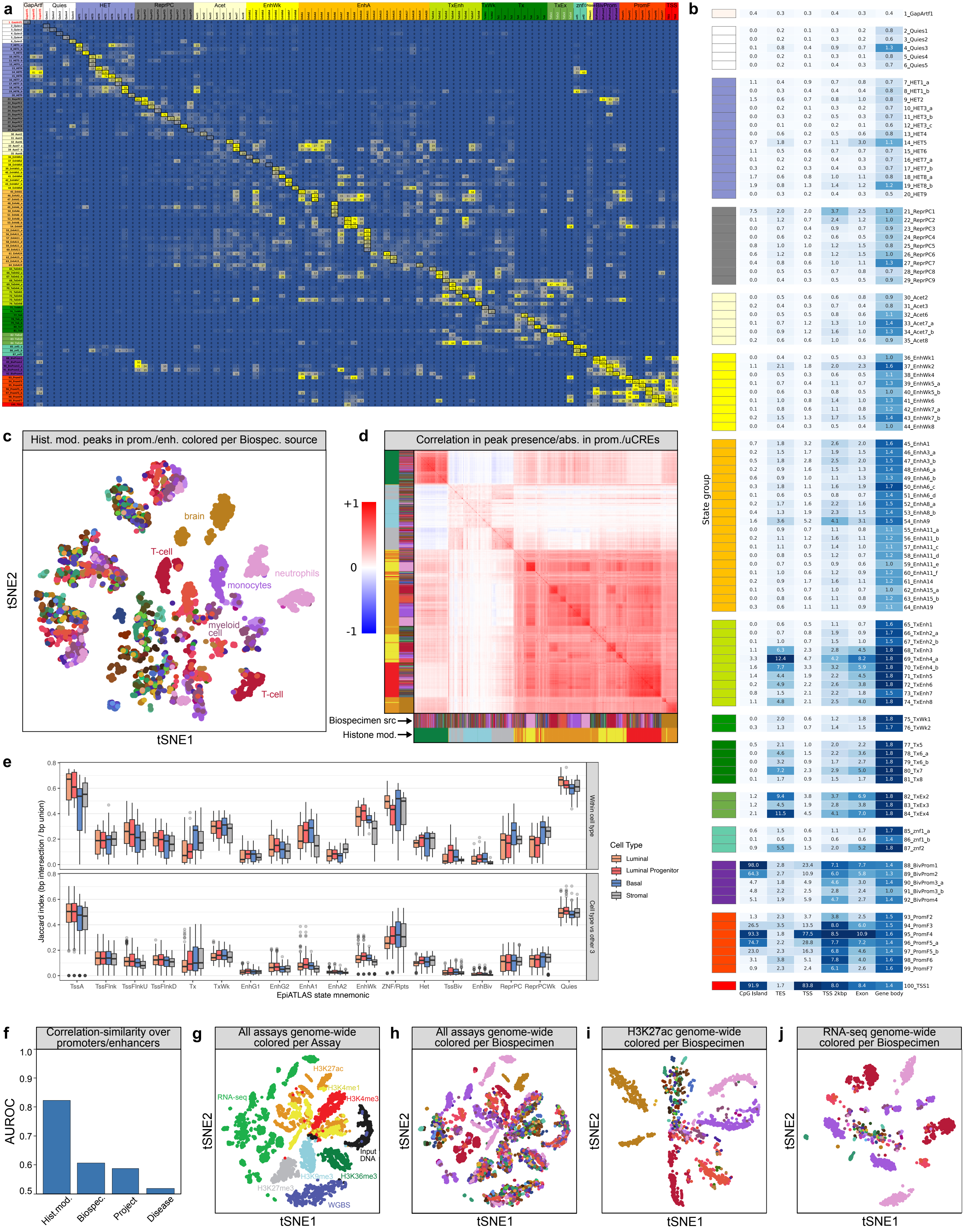

a
GapArtf
Quies
HET
ReprPC
Acet
EnhWk
EnhA
TxEnh
TxWk
Tx
TxEx
znf
BivProm
PromF
TSS
b
State group
CpG Island
TES
TSS
TSS 2kbp
Exon
Gene body
c
Hist. mod. peaks in prom./enh. colored per Biospec. source
brain
T-cell
neutrophils
monocytes
myeloid cell
T-cell
tSNE1
tSNE2
d
Correlation in peak presence/abs. in prom./uCREs
+1
0
-1
Biospecimen src
Histone mod.
e
Jaccard index (bp intersection / bp union)
Within cell type
Cell type vs other 3
TssA
TssFlnk
TssFlnkU
TssFlnkD
Tx
TxWk
EnhG1
EnhG2
EnhA1
EnhA2
EnhWk
ZNF/Rpts
Het
TssBiv
EnhBiv
ReprPC
ReprPCWk
Quies
EpiATLAS state mnemonic
Cell Type
Luminal
Luminal Progenitor
Basal
Stromal
f
Correlation-similarity over promoters/enhancers
AUROC
Hist.mod.
Biospec.
Project
Disease
g
All assays genome-wide colored per Assay
H3K27ac
H3K4me1
H3K4me3
RNA-seq
Input DNA
H3K9me3
H3K36me3
H3K27me3
WGBS
tSNE1
tSNE2
h
All assays genome-wide colored per Biospecimen
i
H3K27ac genome-wide colored per Biospecimen
j
RNA-seq genome-wide colored per Biospecimen

**Extended Data Figure 3| a,** Fold enrichment between states of the EpiATLAS stacked model (rows) and states of a previous stacked chromatin state model based primarily on Roadmap Epigenomics data[13] (columns). The mnemonic of the EpiATLAS stacked model state is based on the mnemonic of the state from this prior model for which it had greatest enrichment. The corresponding enrichments are boxed in this figure. Yellow values correspond to higher enrichments and blue lower. The state group colors are the same as in Vu and Ernst 2022[13]. **b,** EpiATLAS universal chromatin states' enrichments for external genomic annotations. The heatmap shows each of the 100 states' fold enrichments with CpG islands, transcription end sites (TES), transcription start sites (TSS), 2kbp regions surrounding TSS (TSS 2kbp), exons, and gene bodies. Rows correspond to states organized into groups. Each column is colored such that the largest fold enrichment value has the darkest color and the smallest value has the lightest color. **c,** tSNE embedding of ChIP-seq datasets colored by biospecimen source. tSNE embedding of ChIP-seq datasets with same coordinates as **Fig. 4b**, but colored by biospecimen source, indicates the secondary importance of biospecimen source in the structure of the embedding, particularly for H3K27ac and H3K4me1. **d,** Hierarchically-clustered correlation heatmap of the rows of binary peak presence-absence matrix. A heatmap of pairwise correlation distances between binary vectors representing different ChIP-seq datasets indicates that histone modification is the dominant driver of similarity. Rows and columns are hierarchically clustered, and colors indicate histone modification and biospecimen source (**Fig. 1**). **e,** Jaccard index of EpiATLAS ChromHMM states within and across four breast cell types from eight patients. **f,** AUROC statistics for metadata value agreement by correlation similarity. Each histone modification ChIP-seq experiment is represented as a binary vector of peak presence/absence at all promoters and uCREs on stacked ChromHMM calls, and all pairs of samples are traversed in order of decreasing correlation. For each of the four metadata categories (histone mark, project, biospecimen label (harmonized_sample_ontology_intermediate), and disease status (harmonized_sample_disease_high)), agreement of label is counted as a true positive and disagreement as a false positive, and area under the ROC curve is calculated. **g, h,** tSNE embedding of datasets represented by read count densities in 100 kb bins across the genome, colored by assay type (**g**) or biospecimen source (**h**). **i, j,** Same as **h** but only using H3K27ac ChIP-seq data (**i**) or RNA-seq data (**j**).

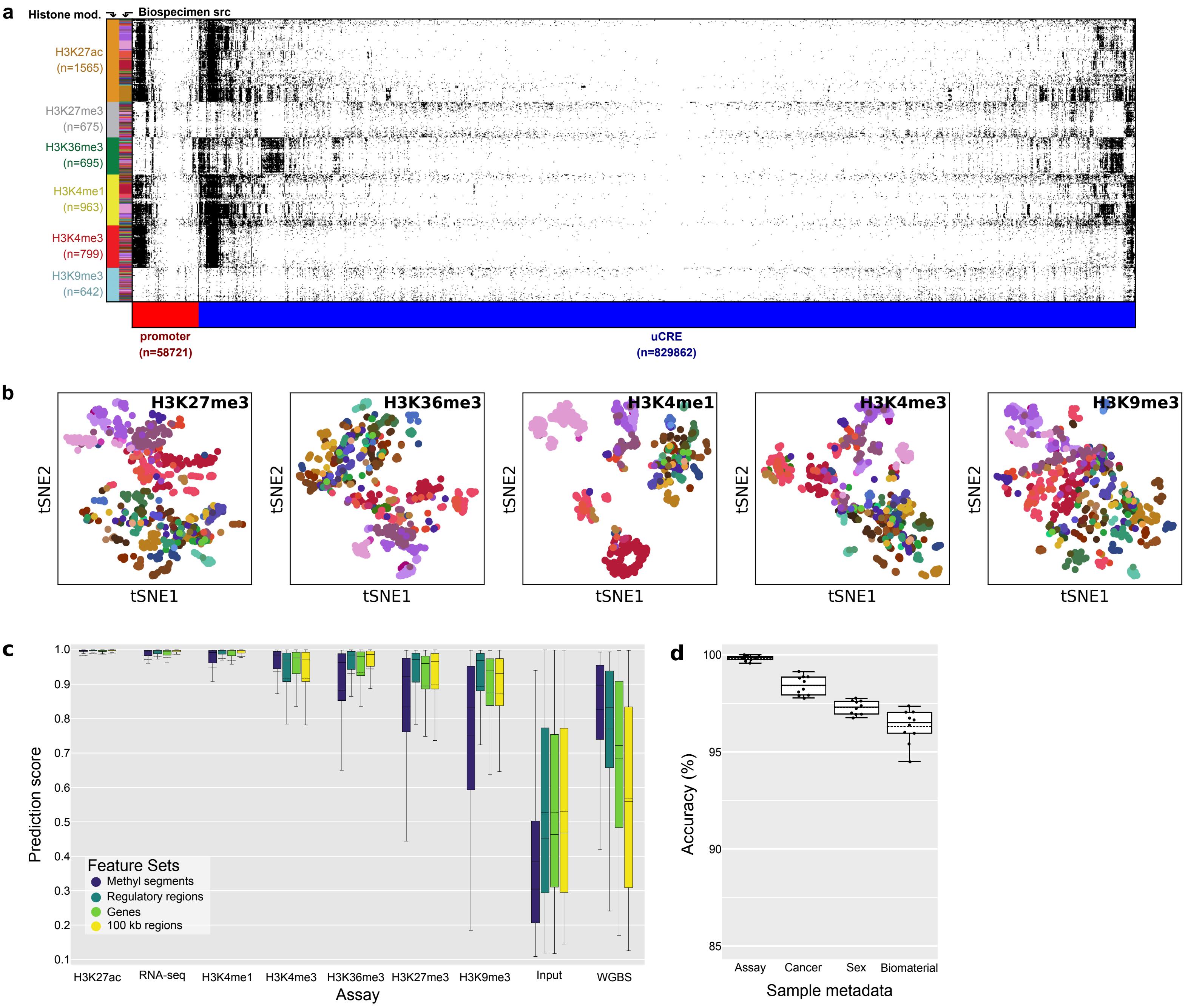

a
Histone mod.
Biospecimen src
H3K27ac (n=1565)
H3K27me3 (n=675)
H3K36me3 (n=695)
H3K4me1 (n=963)
H3K4me3 (n=799)
H3K9me3 (n=642)
promoter (n=58721)
uCRE (n=829862)
b
H3K27me3
H3K36me3
H3K4me1
H3K4me3
H3K9me3
tSNE1
tSNE2
c
Prediction score
Feature Sets
Methyl segments
Regulatory regions
Genes
100 kb regions
H3K27ac
RNA-seq
H3K4me1
H3K4me3
H3K36me3
H3K27me3
H3K9me3
Input
WGBS
Assay
d
Accuracy (%)
Assay
Cancer
Sex
Biomaterial
Sample metadata

**Supplementary Figure 4 | a,** Binary peak presence-absence matrix at promoters (1 kb upstream of the TSS) and regulatory regions (uCREs). Visualization of the binary matrix of peak presence (black) or absence (white) at promoters (red columns) and regulatory regions (blue columns). Each row is one ChIP-seq dataset and is organized and color-labeled by histone modification first and biospecimen source second. Given the size of the matrix, only every 10th row and every 500th column are shown. Columns are hierarchically clustered by Euclidean distance using average linkage agglomerative clustering and optimal leaf ordering. **b,** tSNE embeddings of peak presence-absence vectors by histone mark. tSNE embeddings for peak presence-absence vectors at promoters and regulatory regions for each histone mark's datasets separately, colored by cell/tissue type, show cell/tissue type-specific clustering. Embedding for H3K27ac is shown in **Fig 4c. c,** Distribution of the EpiClass prediction scores for biospecimen category per assay over the 10-fold cross-validation training for the classifier trained on different types of genomic regions (~300k most variable WGBS 200 bp bins, ~300k top-correlated regulatory regions (average size ~2.3 kb), ~19k genes (average size ~68 kb) and 30k non-overlapping 100 kb bins covering the whole genome). **d,** EpiClass accuracy per fold (dots) of cross-validation to predict four selected metadata attributes on the EpiATLAS data. Dashed lines represent means, solid lines the medians, boxes the quartiles, and whiskers the farthest points within 1.5× the interquartile range.

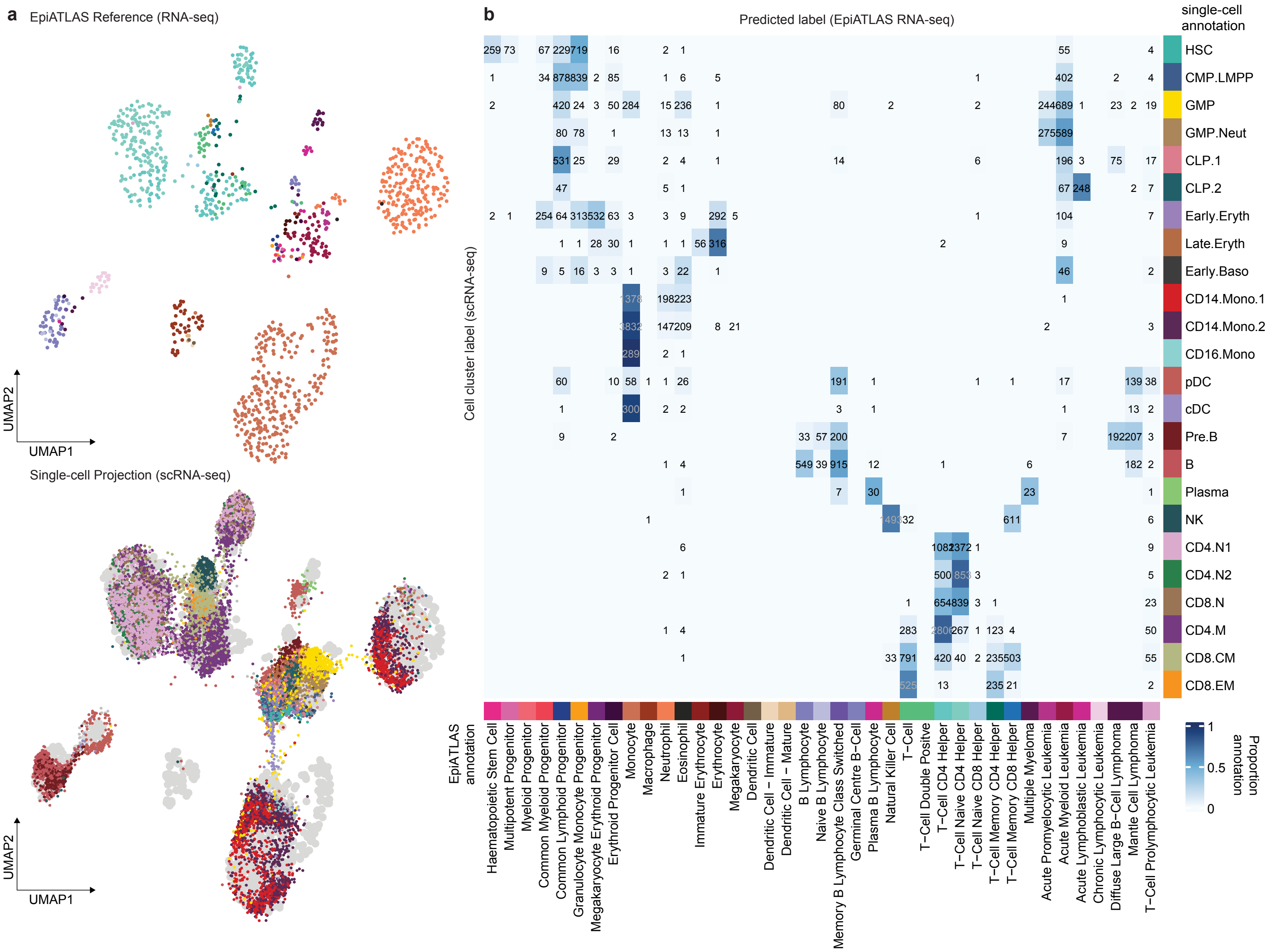

a
EpiATLAS Reference (RNA-seq)
UMAP2
UMAP1
Single-cell Projection (scRNA-seq)
UMAP2
UMAP1
b
Predicted label (EpiATLAS RNA-seq)
Cell cluster label (scRNA-seq)
single-cell annotation
HSC
CMP.LMPP
GMP
GMP.Neut
CLP.1
CLP.2
Early.Eryth
Late.Eryth
Early.Baso
CD14.Mono.1
CD14.Mono.2
CD16.Mono
pDC
cDC
Pre.B
B
Plasma
NK
CD4.N1
CD4.N2
CD8.N
CD4.M
CD8.CM
CD8.EM
EpiATLAS annotation
Haematopoietic Stem Cell
Multipotent Progenitor
Myeloid Progenitor
Common Myeloid Progenitor
Common Lymphoid Progenitor
Granulocyte Monocyte Progenitor
Megakaryocyte Erythroid Progenitor
Erythroid Progenitor Cell
Monocyte
Macrophage
Neutrophil
Eosinophil
Immature Erythrocyte
Erythrocyte
Megakaryocyte
Dendritic Cell
Dendritic Cell − Immature
Dendritic Cell − Mature
B Lymphocyte
Naive B Lymphocyte
Memory B Lymphocyte Class Switched
Germinal Centre B−Cell
Plasma B Lymphocyte
Natural Killer Cell
T−Cell
T−Cell Double Positve
T−Cell CD4 Helper
T−Cell Naive CD4 Helper
T−Cell Naive CD8 Helper
T−Cell Memory CD4 Helper
T−Cell Memory CD8 Helper
Multiple Myeloma
Acute Promyelocytic Leukemia
Acute Myeloid Leukemia
Acute Lymphoblastic Leukemia
Chronic Lymphocytic Leukemia
Diffuse Large B−Cell Lymphoma
Mantle Cell Lymphoma
T−Cell Prolymphocytic Leukemia
Proportion annotation
1
0.5
0

**Extended Data Figure 4 | Mapping of single-cell transcriptomes to the EpiATLAS reference**.

**a,** UMAP embedding of blood-related EpiATLAS transcriptomes (top) and projection of single-cell transcriptomes into the embedding (bottom). EpiATLAS bulk cell types are annotated from the 'harmonized_sample_label' metadata. Single-cell transcriptomes and cell type annotation were taken from Granja et al 2019[14]. In the projection UMAP, the EpiATLAS reference embedding is depicted as grey colored points. **b,** Heatmap denoting the number of predictions of single-cell transcriptomes as EpiATLAS annotated cell types ('harmonized_sample_label'). Values are absolute and colors are scaled by row. Predictions were made using classification with linear discriminant analysis (LDA) based on projected principal component embeddings.

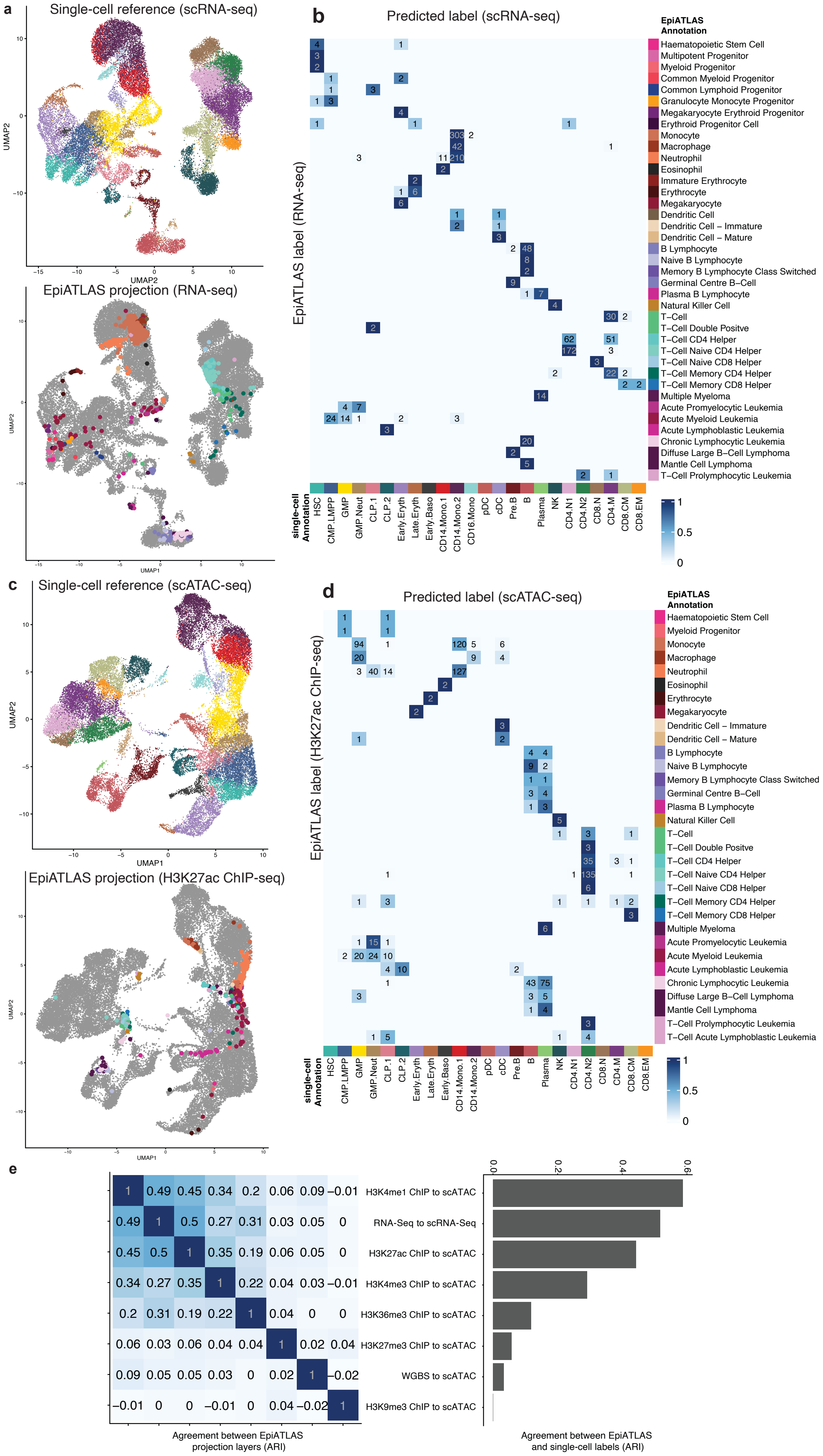
a
Single-cell reference (scRNA-seq)
UMAP1
UMAP2
EpiATLAS projection (RNA-seq)
b
Predicted label (scRNA-seq)
EpiATLAS label (RNA-seq)
EpiATLAS Annotation
Haematopoietic Stem Cell
Multipotent Progenitor
Myeloid Progenitor
Common Myeloid Progenitor
Common Lymphoid Progenitor
Granulocyte Monocyte Progenitor
Megakaryocyte Erythroid Progenitor
Erythroid Progenitor Cell
Monocyte
Macrophage
Neutrophil
Eosinophil
Immature Erythrocyte
Erythrocyte
Megakaryocyte
Dendritic Cell
Dendritic Cell – Immature
Dendritic Cell – Mature
B Lymphocyte
Naive B Lymphocyte
Memory B Lymphocyte Class Switched
Germinal Centre B–Cell
Plasma B Lymphocyte
Natural Killer Cell
T–Cell
T–Cell Double Positve
T–Cell CD4 Helper
T–Cell Naive CD4 Helper
T–Cell Naive CD8 Helper
T–Cell Memory CD4 Helper
T–Cell Memory CD8 Helper
Multiple Myeloma
Acute Promyelocytic Leukemia
Acute Myeloid Leukemia
Acute Lymphoblastic Leukemia
Chronic Lymphocytic Leukemia
Diffuse Large B–Cell Lymphoma
Mantle Cell Lymphoma
T–Cell Prolymphocytic Leukemia
single-cell Annotation
HSC
CMP.LMPP
GMP
GMP.Neut
CLP.1
CLP.2
Early.Eryth
Late.Eryth
Early.Baso
CD14.Mono.1
CD14.Mono.2
CD16.Mono
pDC
cDC
Pre.B
B
Plasma
NK
CD4.N1
CD4.N2
CD8.N
CD4.M
CD8.CM
CD8.EM
1
0.5
0
c
Single-cell reference (scATAC-seq)
EpiATLAS projection (H3K27ac ChIP-seq)
d
Predicted label (scATAC-seq)
EpiATLAS label (H3K27ac ChIP-seq)
T–Cell Acute Lymphoblastic Leukemia
e
H3K4me1 ChIP to scATAC
RNA-Seq to scRNA-Seq
H3K27ac ChIP to scATAC
H3K4me3 ChIP to scATAC
H3K36me3 ChIP to scATAC
H3K27me3 ChIP to scATAC
WGBS to scATAC
H3K9me3 ChIP to scATAC
Agreement between EpiATLAS projection layers (ARI)
Agreement between EpiATLAS and single-cell labels (ARI)

**Supplementary Figure 5 | Single-cell views of the EpiATLAS. a,** UMAP embedding of single-cell RNA-seq of human hematopoiesis (top) and projection of IHEC EpiATLAS bulk RNA-seq profiles into the embedding (bottom). Single-cell labels are annotated by Granja et al 2019[14] and EpiATLAS bulk cell type labels were taken from the 'harmonized_sample_label' annotation. Single-cell reference is depicted as grey colored points in the projection UMAP. **b,** Heatmap denoting the number of predictions of EpiATLAS bulk RNA-seq samples with annotated cell types ('harmonized_sample_label') as single-cell labels (scRNA-seq). Values are absolute and colors are scaled by row. Predictions were made using classification with linear discriminant analysis (LDA) based on projected principal component embeddings. **c,d,** Analogue to **a,b,** but projecting IHEC EpiATLAS bulk H3K27ac ChIP-seq profiles into an embedding of single-cell ATAC-seq. The embedding is based on aggregating Tn5 insertion counts of scATAC-seq data across uCREs. **e,** Comparison between different EpiATLAS projection layers in their agreement of predicted single-cell labels with each other (left) and the EpiATLAS bulk labels ('harmonized_sample_label') (right). Single-cell label classification was performed using LDA based on projected principal component embeddings. Agreement was quantified by the Adjusted Rand Index (ARI).

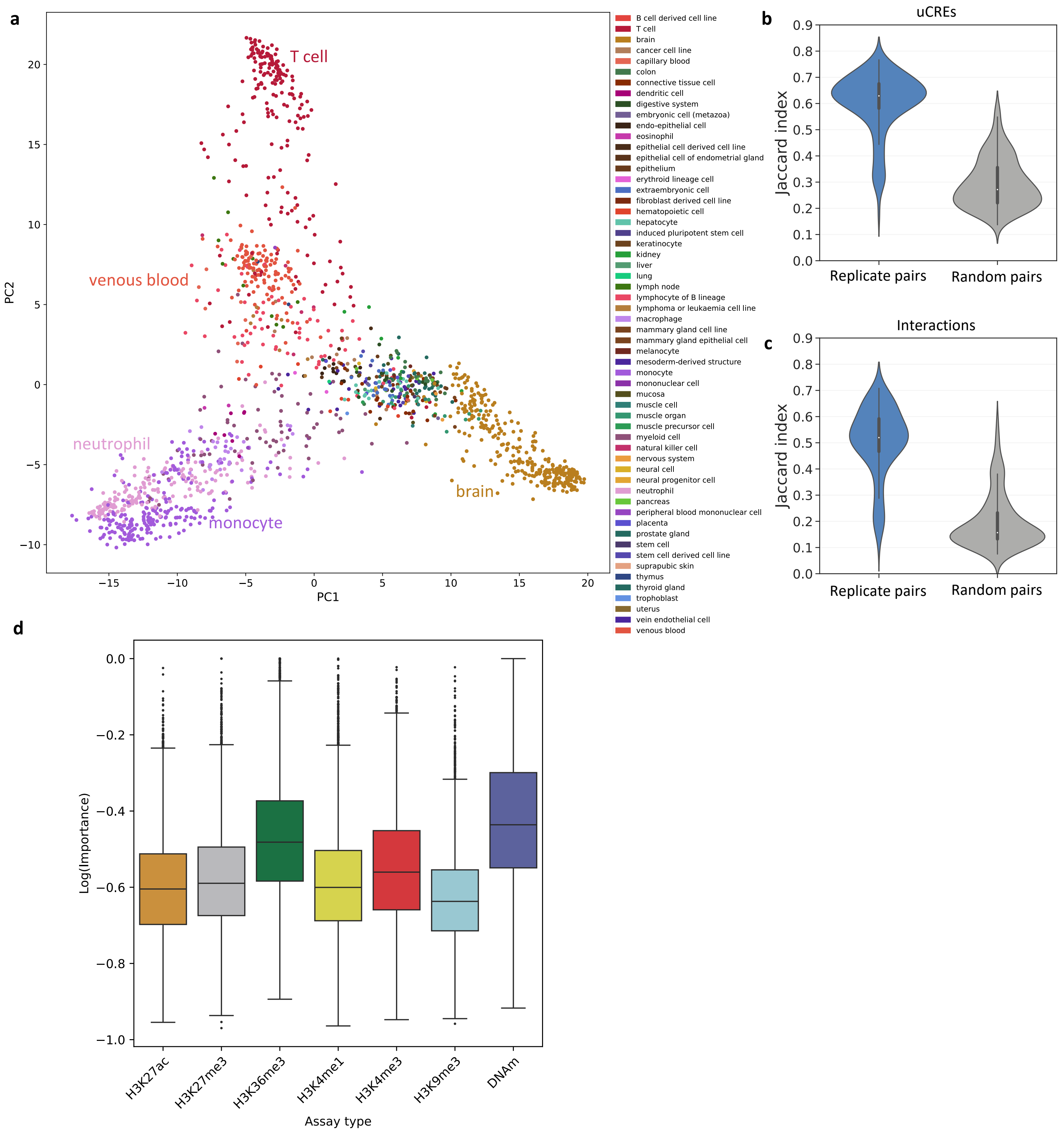
a
T cell
venous blood
neutrophil
monocyte
brain
PC1
PC2
B cell derived cell line
T cell
brain
cancer cell line
capillary blood
colon
connective tissue cell
dendritic cell
digestive system
embryonic cell (metazoa)
endo-epithelial cell
eosinophil
epithelial cell derived cell line
epithelial cell of endometrial gland
epithelium
erythroid lineage cell
extraembryonic cell
fibroblast derived cell line
hematopoietic cell
hepatocyte
induced pluripotent stem cell
keratinocyte
kidney
liver
lung
lymph node
lymphocyte of B lineage
lymphoma or leukaemia cell line
macrophage
mammary gland cell line
mammary gland epithelial cell
melanocyte
mesoderm-derived structure
monocyte
mononuclear cell
mucosa
muscle cell
muscle organ
muscle precursor cell
myeloid cell
natural killer cell
nervous system
neural cell
neural progenitor cell
neutrophil
pancreas
peripheral blood mononuclear cell
placenta
prostate gland
stem cell
stem cell derived cell line
suprapubic skin
thymus
thyroid gland
trophoblast
uterus
vein endothelial cell
venous blood
b
uCREs
Jaccard index
Replicate pairs
Random pairs
c
Interactions
Jaccard index
Replicate pairs
Random pairs
d
Log(Importance)
H3K27ac
H3K27me3
H3K36me3
H3K4me1
H3K4me3
H3K9me3
DNAm
Assay type

**Extended Data Figure 5 | a,** PCA of the predicted gABC[9] interactions and their scores across biospecimens, using only interactions from chromosome 1. Scores $\leq 0.02$ were set to zero. Samples coloured by the metadata field harmonized_sample_ontology_intermediate (metadata version 2.0). **b,** Jaccard index of the set of putative cis-regulatory elements based on the universal ChromHMM annotation (uCREs) overlapping a H3K27ac ChIP-seq peak in a biospecimen between all possible pairs of biospecimen replicates (n = 135), compared to the same number of random non-replicate pairs. Replicates were defined as biospecimens where all metadata fields match except the EpiRR ID. **c,** Same as **b,** but calculating the Jaccard index for the overlap of uCRE-gene interactions predicted with the gABC score. **d,** Feature importance per histone mark and DNA methylation of a Graph Convolutional Neural Network (GCN) predicting transcript expression. The importance is summarised across the gene regions that were used as features.

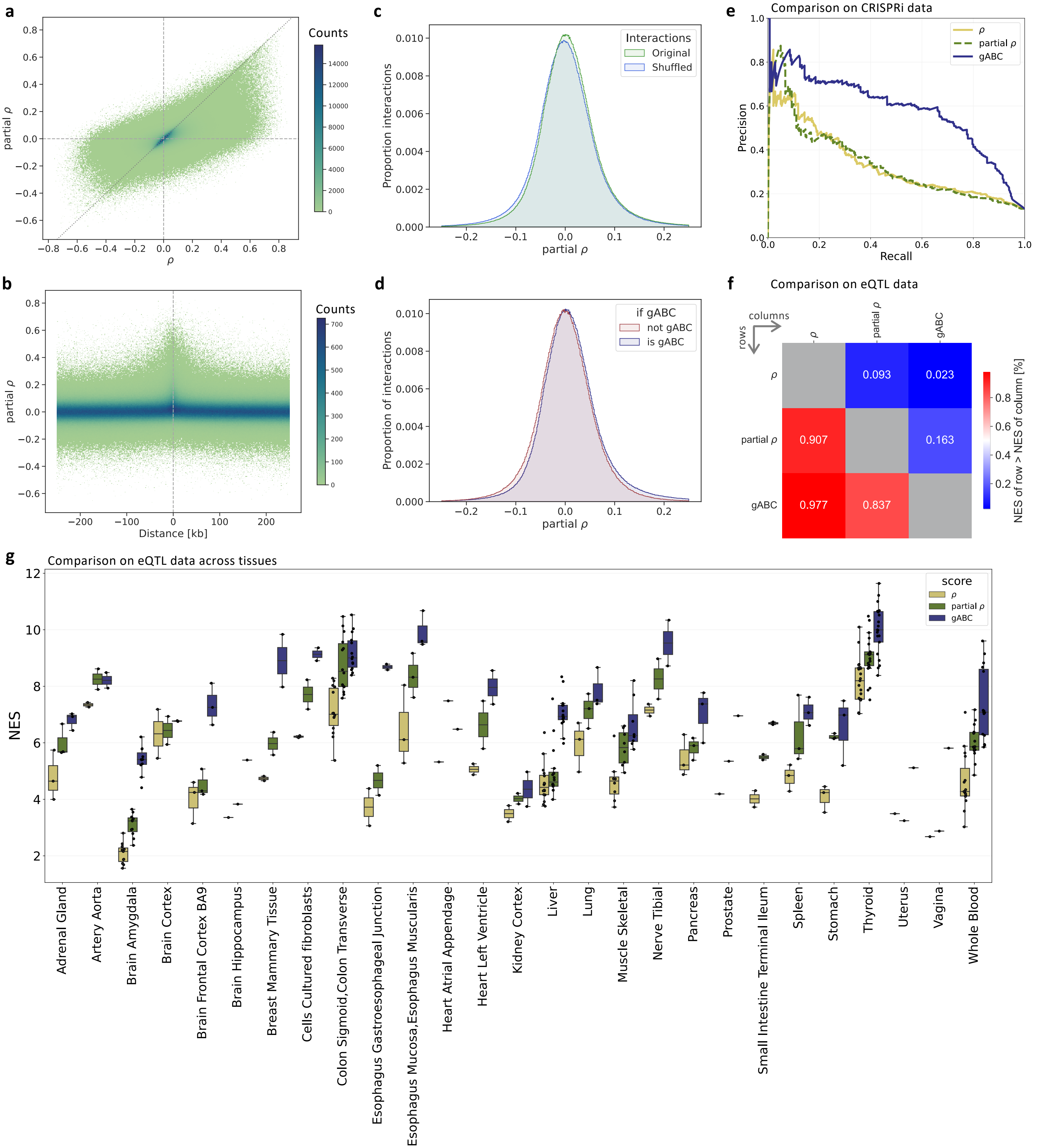

a
Counts
partial ρ
ρ
b
Counts
partial ρ
Distance [kb]
c
Interactions
Original
Shuffled
Proportion interactions
partial ρ
d
if gABC
not gABC
is gABC
Proportion of interactions
partial ρ
e
Comparison on CRISPRi data
ρ
partial ρ
gABC
Precision
Recall
f
Comparison on eQTL data
columns
rows
ρ
partial ρ
gABC
0.093
0.023
0.907
0.163
0.977
0.837
NES of row > NES of column [%]
g
Comparison on eQTL data across tissues
score
ρ
partial ρ
gABC
NES
Adrenal Gland
Artery Aorta
Brain Amygdala
Brain Cortex
Brain Frontal Cortex BA9
Brain Hippocampus
Breast Mammary Tissue
Cells Cultured fibroblasts
Colon Sigmoid,Colon Transverse
Esophagus Gastroesophageal Junction
Esophagus Mucosa,Esophagus Muscularis
Heart Atrial Appendage
Heart Left Ventricle
Kidney Cortex
Liver
Lung
Muscle Skeletal
Nerve Tibial
Pancreas
Prostate
Small Intestine Terminal Ileum
Spleen
Stomach
Thyroid
Uterus
Vagina
Whole Blood

**Supplementary Figure 6 |** Correlation analyses between H3K27ac and RNA-seq. **a,** Spearman correlation coefficient (ρ) between a uCRE's H3K27ac signal (mean fold-change over the input) and the RNA-seq (TPM) of the associated gene versus the partial Spearman correlation coefficient (partial ρ) for all possible uCRE-gene pairs that span ≤ 250kb (n = 7,865,786). The partial correlation controls for the expression of all other genes within 250 kb. Interactions where the uCRE overlaps a promoter were excluded (± 200 bp around any annotated TSS). **b,** Partial correlation coefficient versus the distance between uCRE and gene. **c,** Distribution of the partial correlation coefficient of the interactions (Original) and interactions that were shuffled (Shuffled). For the shuffled interactions, we randomly rewired the uCRE-gene pairs, allowing also interchromosomal interactions, following Xie et al. (2023)[15]. The x-axis is limited to -0.25 to 0.25. **d,** Same distribution of the partial correlation coefficients of the original interactions from **c,** but split by whether an interaction surpassed the gABC cutoff in any biospecimen (≥ 0.02). **e,** Precision recall curve for identifying CRISPRi-validated uCRE-gene interactions[16]. Only interactions where all three metrics produced a score were used, resulting in 280 positives out of 2117 validated interactions. Area under the curve for ρ ≈ 0.331, partial ρ ≈ 0.335, and gABC ≈ 0.577. **f,** Comparison of the normalized enrichment score (NES) of eQTL-supported uCRE-gene interactions among interactions ranked by ρ, partial ρ or gABC-score. An interaction was considered supported if an overlapping eQTL was associated with the same gene. The higher the NES, the higher the ranking of the interactions with eQTL support. Shown are the fractions of biospecimens from **g** for which one metric achieved a higher NES than another metric. The same uCRE-gene pairs were ranked for each metric, and for each biospecimen, only interactions where the uCRE overlapped a H3K27ac ChIP-seq peak from that sample were kept. The NES was calculated with GSEApy[17]. eQTL-gene pairs fine-mapped with DAP-G from GTEx were used[18,19]. **g,** Distribution of the NES across GTEx tissues that were matched to EpiATLAS biospecimens (boxplot center line the median, box limits inter-quartile range, whiskers up to 1.5x inter-quartile range). Each dot represents one EpiATLAS biospecimen.

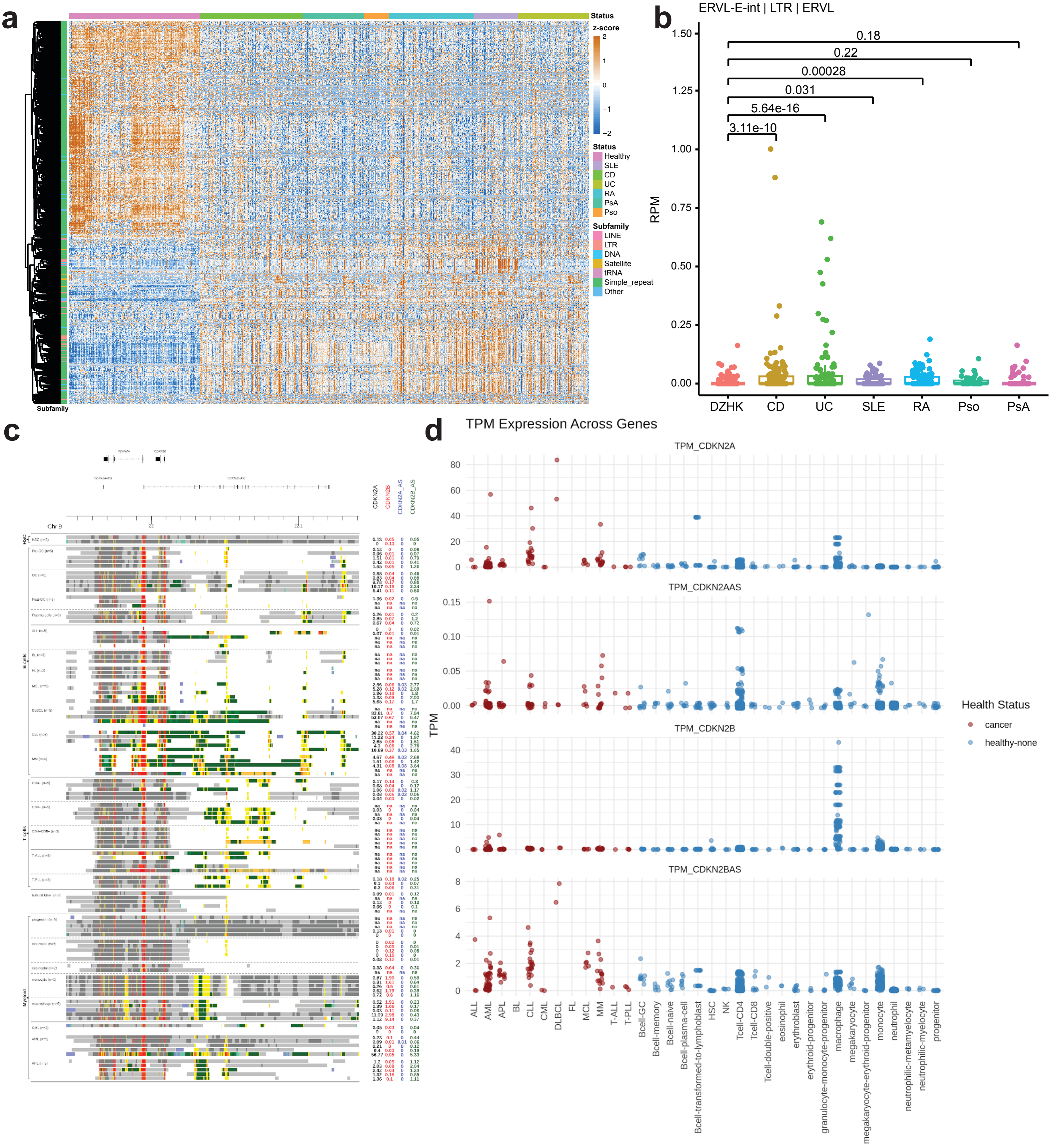

a
Status
z-score
2
1
0
−1
−2
Status
Healthy
SLE
CD
UC
RA
PsA
Pso
Subfamily
LINE
LTR
DNA
Satellite
tRNA
Simple_repeat
Other
Subfamily
b
ERVL-E-int | LTR | ERVL
RPM
1.50
1.25
1.00
0.75
0.50
0.25
0.00
0.18
0.22
0.00028
0.031
5.64e-16
3.11e-10
DZHK
CD
UC
SLE
RA
Pso
PsA
c
Chr 9
CDKN2A
CDKN2B
CDKN2A_AS
CDKN2B_AS
HSC
B cells
T cells
Myeloid
d
TPM Expression Across Genes
TPM_CDKN2A
TPM_CDKN2AAS
TPM_CDKN2B
TPM_CDKN2BAS
TPM
Health Status
cancer
healthy-none
ALL
AML
APL
BL
CLL
CML
DLBCL
FL
MCL
MM
T-ALL
T-PLL
Bcell-GC
Bcell-memory
Bcell-naive
Bcell-plasma-cell
Bcell-transformed-to-lymphoblast
HSC
NK
Tcell-CD4
Tcell-CD8
Tcell-double-positive
eosinophil
erythroblast
erythroid-progenitor
granulocyte-monocyte-progenitor
macrophage
megakaryocyte
megakaryocyte-erythroid-progenitor
monocyte
neutrophil
neutrophilic-metamyelocyte
neutrophilic-myelocyte
progenitor

**Extended Data Figure 6 | a,** Dysregulation of repetitive elements in Chronic Inflammatory Diseases (CID) patients. Heatmap showing differentially expressed individual repetitive elements (n = 5723) between disease and healthy controls in RNA-seq datasets (top). Datasets are obtained from the SYSCID data repository. Upregulated individual repetitive elements (Fold change > 4 and two-tailed Wald test p.value < 1e-4) and downregulated individual repetitive elements (Fold change < 0.25 and two-tailed Wald test p.value < 1e-4) are called by comparing all disease patients to healthy controls. Color bar represents the normalized total count calculated z-score. Differential expression was tested using the negative binomial generalized linear model. **b,** A boxplot showing an example of a disease-specific differentially expressed individual repetitive element (bottom). The individual ERVL-E-int element (chromosome location chr19:43365063-43365376) is significantly upregulated (Fold change > 4 and p.value < 1e-4) only in Inflammatory Bowel Disease patients (CD and UC) but not in other disease patients (SLE, RA, PsA). Each dot represents one individual. The centre and bounds of boxes indicate the median and quartile of all data points, respectively. The minima and maxima of whiskers indicate quartile 1-1.5×the interquartile range and quartile 3+1.5×the interquartile range, respectively. P-values are calculated using DESeq2 two-tailed Wald test. Healthy (N = 197); CD: Crohn's Disease (N = 155); UC: Ulcerative Colitis (N = 107); SLE: Systemic Lupus Erythematosus (N = 66); RA: Rheumatoid Arthritis (N = 128); Pso: Psoriasis (N = 38); PsA: Psoriatic Arthritis (N = 93). **c,** Integration of chromatin state segmentation with RNA-seq data across hematopoietic cell types and associated malignancies at the *CDKN2A/B* locus. Chromatin profiles (left) are shown alongside gene expression levels (right; TPM) for *CDKN2A*, *CDKN2B*, *CDKN2A-AS1* and *CDKN2B-AS1*. **d,** Gene expression values for all healthy and malignant hematopoietic samples with RNA-seq data available. Despite the enrichment of the transcription-associated chromatin states in malignant samples, expression of *CDKN2A* and *CDKN2B* remains low or variable and is not consistently increased compared to normal B-cell populations. In contrast, the antisense transcript *CDKN1B-AS1* shows relatively higher and more consistent expression in malignant samples. TPM: transcripts per million; HSC: hematopoietic stem cells; GC: germinal center; PCs: plasma cells; ALL: acute lymphoblastic leukemia; BL: Burkitt lymphoma; FL: follicular lymphoma; MCL: mantle cell lymphoma; DLBCL: diffuse large B-cell lymphoma; CLL: chronic lymphocytic leukemia; MM: multiple myeloma; T-ALL: T-cell acute lymphoblastic leukemia; T-PLL: T-cell prolymphocytic leukemia; CML: chronic myeloid leukemia; AML: acute myeloid leukemia; APL: acute promyelocytic leukemia.

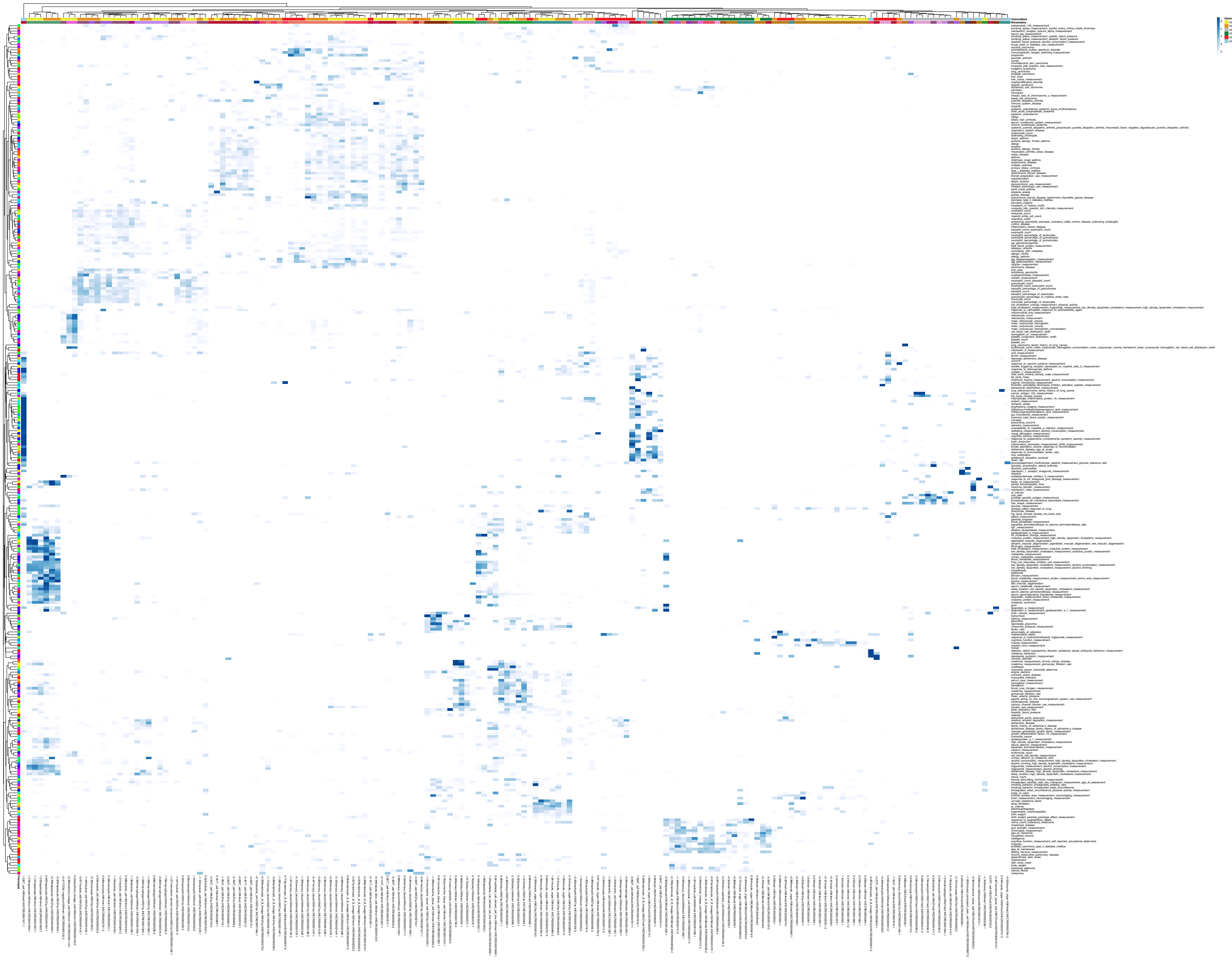

**Supplementary Figure 7 |** High-resolution global visualisation of FORGE2 functional overlap analysis.

Showing the same data as in **Fig. 8c,** including all row and column labels, but at high resolution. See also **Supplementary Table 7**.
