## Supplementary Data 01 for "EpiATLAS – a reference for human epigenomic research"

### IHEC EpiAtlas Integrative Analysis
### Data Processing and Sharing

*Misha Bilenky, Young Cheng, Joe Chen, Edmund Su, Annaick Carles, Sinead Aherne, Reanne Bowlby, Sitanshu Gakkhar, Patrick Plettner, Jonathan Steif, Martin Hirst.*

#### Document version: Data Freeze (previous version 12.2)
last updated: December/9/2022

#### Table of Contents

Document Purpose and Scope

Definitions

Data Overview

Data organization on SFTP

Analysis File Nomenclature

Data Processing Pipelines

Consensus Resources

Sample Metadata Harmonization Project

Data Processing Decision Tree

- Dealing with mixed read lengths in ChIP-Seq data
- Redundant Input Analysis
- Dealing with mixed read lengths in RNA-Seq data

ChIP File Types For Dissemination

WGBS File Types For Dissemination

RNA File Types For Dissemination

Data/Metadata Update Notes

Data Posting Dates

#### Document Purpose and Scope
This document provides an explanation for the processes and file types generated as part of the International Human Epigenome Consortium EpiAtlas integrative analysis.

#### Definitions
**REFERENCE EPIGENOME**

➔ A collection of ANALYSIS objects performed on a biosample represented by an **EpiRR identifier.**

**ANALYSIS**

➔ An object that references one or more EXPERIMENTs. The ANALYSIS object does not have a unique mapping to an object in the EpiRR registry .

**EXPERIMENT**

➔ A collection of one or more **EGAR/SRR/ENCFF** objects represented by an **EGAX/SRX/ENCSR identifie**r in the EpiRR registry.

**DATASET**

➔ An **EGAR/SRR/ENCFF** object that references one or more **fastq** files.

#### Data Overview

*Data represents **337 complete** and **2022 partial** epigenomes.*

| IHEC Consortium | Summary (Epigenomes) | | | Summary (Analyses) | | | | |
|---|---|---|---|---|---|---|---|---|
| | # EpiRR | # Human References | # Human Complete References | # Analysis Objects | # Staged | # Processed | #Posted | #Failed |
| **BLUEPRINT** | 1249 | 1249 | 112 | 3829 | 3430 | 3430 | 3422 | 407 |
| **NIH Roadmap Epigenomics** | 147 | 147 | 27 | 826 | 518 | 499 | 764 | 62 |
| **DEEP** | 157 | 60 | 34 | 370 | 364 | 346 | 342 | 28 |
| **AMED-CREST** | 278 | 258 | 20 | 548 | 310 | 299 | 295 | 253 |
| **ENCODE** | 210 | 118 | 18 | 784 | 726 | 726 | 726 | 58 |
| **Korea Epigenome Project (KNIH)** | 37 | 37 | 0 | 74 | 61 | 51 | 50 | 24 |
| **CEEHRC (CEMT up to CEMT_212)** | 125 | 125 | 107 | 1035 | 1028 | 1028 | 1028 | 7 |
| **CEEHRC (McGill)** | 343 | 338 | 32 | 898 | 891 | 824 | 818 | 80 |
| **GIS** | 291 | 291 | 0 | 291 | 288 | 287 | 287 | 4 |
| **EpiHK** | 1 | 1 | 1 | 8 | 8 | 8 | 8 | 0 |
| **Total** | **2897** | **2691** | **346** | **8663** | **7624** | **7498** | **7740** | **923** |

**Final data posted on SFTP (December 08, 2022)**

| IHEC Consortium | # Posted RNA-seq | # Posted Bisulfite-seq | # Posted ChIP-seq |
|---|---|---|---|
| **BLUEPRINT** | 868 | 225 | 2329 |
| **NIH Roadmap Epigenomics** | 61 | 55 | 648 |
| **DEEP** | 86 | 37 | 219 |
| **AMED-CREST** | 33 | 46 | 216 |
| **ENCODE** | 99 | 27 | 600 |
| **Korea Epigenome Project (KNIH)** | 13 | 37 | 0 |
| **CEEHRC (CEMT up to CEMT_212)** | 128 | 123 | 777 |
| **CEEHRC (McGill)** | 333 | 94 | 391 |
| **GIS** | 0 | 0 | 287 |
| **EpiHK** | 1 | 1 | 6 |
| **Total** | **1622** | **645** | **5473** |

A list of data that was not processed can be found here: IHEC EpiAtlas Edge Cases 2022

#### Data organization on SFTP

The folder **incoming/readme/** contains the following files:

- ➔ **epiatlas_metadata.csv**
  - This file contains metadata for all analysis objects combined
    - RNA-Seq: 1622
    - WGBS: 645
    - ChIP-Seq: 5473
  - This file combines the three previously posted separate metadata files (ihec_metada.csv, ihec_metadata_rna.csv and ihec_metdata_wgbs.csv) with all fields harmonized.
  - The 2nd column in the file records the status of the EpIRR ID (complete/partial) [this differs slightly from the status @ EpiRR due to failure/retraction of few datasets]
  - The last column of epiatlas_metadata.csv contains the path to all analysis files corresponding to the analysis object.
  - If a certain metric/info is not applicable for a particular assay ‘nan’ has been entered (e.g. information on ‘ctl’ samples for RNA-Seq and WGBS).
  - The file contains 18 fields which are listed in the table below:

| | Field | Description |
|---|---|---|
| 1 | epirr_id | IHECRE01234567.8 (as recorded @ EpiRR) |
| 2 | status | complete/partial |
| 3 | data_generating_centre | Name of one of 10 data generating centers |
| 4 | assay_type | ChIP-Seq, RNA-Seq, WGBS |
| 5 | experiment_type | ChIP-Seq: antibody (H3K4me3, H3K27ac, H3K4me1, H3K36me3, H3K27me3, H3K9me3)<br>WGBS: standard or PBAT protocols<br>RNA-Seq: mRNA-Seq or total-RNA-Seq |
| 6 | uuid | unique universal identifier |
| 7 | inputs | ";" separated list of file IDs from the sequence archive (EGAR, SRR or ENCFF) |
| 8 | inputs_ctl | same as above for the DNA Input control (only for ChIp_Seq) |
| 9 | original_read_length | ";" separated list of original read lengths |
| 10 | original_read_length_ctl | same as above for the DNA Input control (only for ChIp_Seq) |
| 11 | trimmed_read_length | ";" separated list of read lengths after trimming (actually used for the analysis) |
| 12 | trimmed_read_length_ctl | same as above for the DNA Input control (only for ChIp_Seq) |
| 13 | software_version | the version of the container pipeline |
| 14 | chastity_passed | whether the reads used in the analysis were chastity filtered |
| 15 | paired_end | whether paired-end or single-end mode was used for the analysis |
| 16 | analyzed_as_stranded | whether the analysis was performed as stranded (only for RNA-Seq) |
| 17 | upload_date | MM-DD-YYYY of the upload |
| 18 | data_file_path | data path on SFTP: e.g. incoming/RNA-Seq/05-16-2022/ihec.rna-seq.ihec-grapenf-containerv1.1.0.IHECRE00004719.1.94bf25c4-a5c0-4e8f-8667-5600e67b184d.* |

- ➔ **epiatlas_metadata_14_partly_revoked_ChIP_Seq..csv**
  - ◆ This file contains metadata information for 14 ChIP-Seq datasets for which processed analysis objects were revoked and only alignment files (BAM and raw.bigwig) for the histone modifications were posted in incoming/ChIP-Seq/12-06-2022/
- ➔ **QC Files:** All three QC files contain an analysis object **uuid**, the **data_generating_centre**, and **epirr_id** as the first 3 columns.
  - ◆ **epiatlas_chipseq_qc_summary.csv**
    - Summarizes 89 QC metrics from qc.json generated by ChIP-Seq container
  - ◆ **epiatlas_rnaseq_qc_summary.csv**
    - Summarizes 5 QC metrics for each RNA-Seq object
  - ◆ **epiatlas_wgbs_qc_summary.csv**
    - Reports mean CpG_Coverage, bisulfite conversion rate, and GC bias (GC content coverage correlation)
- ➔ **The readme file (this document: ihec_readme.pdf)**

#### Analysis File Nomenclature

- ➔ ihec.<epirr_id>.<analysis_type>.<pipeline&version>.<uuid>
  - ◆ epirr_id will include the epirr minor version
  - ◆ experiment type will be one of: chipseq, bsseq, rnaseq
  - ◆ uuid - a unique alpha-numeric string (lower-case)
- ➔ Example:
  - ◆ ihec.IHEC98509485.1.chipseq.ihec_chip_containerv1.4.0.948hr93fnvnvfnjf8499i3r

#### Data Processing Pipelines

- ChIP-seq data was processed using the following container pipeline:
  - https://github.com/IHEC/integrative_analysis_chip/releases
- WGBS data was processed using the following container pipeline:
  - https://github.com/heathsc/gemBS/releases/tag/v3.5.0
- RNA-seq data was processed using the following container pipeline:
  - https://github.com/IHEC/grape-nf/releases

#### Consensus Resources

**GRCh38 REFERENCE BUILD**

- The GRCh38 reference build chosen by IHEC is available at:
  - http://www.epigenomes.ca/data/CEMT/resources/GCA_000001405.15_GRCh38_no_alt_analysis_set.fna.gz
  - http://www.epigenomes.ca/data/CEMT/resources/GCA_000001405.15_GRCh38_no_alt_analysis_set.fna.gz,md5sum
- and via EBI at:
  - http://ftp.ebi.ac.uk/pub/databases/blueprint/reference/20150407_reference_files/GRCh38_no_alt_analysis_set.201503031.fa.gz
- The NCBI url is:
  - ftp://ftp.ncbi.nlm.nih.gov/genomes/archive/old_genbank/Eukaryotes/vertebrates_mammals/Homo_sapiens/GRCh38/seqs_for_alignment_pipelines/GCA_000001405.15_GRCh38_no_alt_analysis_set.fna.gz

**GENE ANNOTATIONS**

- Please note that the IHEC has moved from v22 to v29 annotations. For original v22 references see
  - http://www.epigenomes.ca/data/CEMT/resources/index.v22.html
- The v29 comprehensive gene annotations are available at:
  - ftp://ftp.ebi.ac.uk/pub/databases/gencode/Gencode_human/release_29/gencode.v29.annotation.gtf.gz
- For full details, see: Gencode v29 (Note GENCODE 29 corresponds to the Ensembl *Homo_sapens* hg38v94)

#### Sample Metadata Harmonization Project

- Discussion and version 1 can be found here:
  - https://github.com/IHEC/epimap-metadata-harmonization/tree/main/openrefine/v1.0

#### Data Processing Decision Tree

- Sequence depth goals:
  - ChIP-seq: 50M for narrow marks (H3K27ac,H3K4me3), 100M for broad marks (H3K4me1,H3K9me3,H3K27me3,H3K36me3) and 100M for Input control.
  - WGBS: 30X coverage of the genome.
  - RNA-seq: a minimum depth of ~200 million paired-end reads per replicate (representing 100 million cDNA fragments).
- ENCODE data:
  - Archived data will not be used in the EpiAtlas analysis.
- FASTQ extraction:
  - From BAM using SamToFastq (Picard tools)
    - Blueprint (Bisulfite-Seq)
    - GIS (ChIP-Seq)
  - From SRA binary using fastq-dump (SRA toolkit)
    - Roadmap (all)
- Chastity filtering (if the chastity information is available from the data generators):
  - Only reads that have passed the chastity filter will be used for analysis.
  - Files derived from fastq-dump don't contain chastity filter status, so all reads are used
- Data merging:
  - In instances where there are multiple EGAX identifiers for a given EpiRR for the **same** experiment type, sequenced data from multiple EGAX IDs are merged and processed together to form a single ANALYSIS object.
  - For example, in IHECRE00000016; the 2 EGAX IDs for Histone H3K27ac and the 2 EGAX IDs for ChIP-Seq Input were merged together.
- Trimming:
  - ChIP-Seq:

        - The EGA metadata does not indicate what readlength the data submitter intends for use in reference alignment, for example, it may not be clear if the phasing base is left untrimmed.
        - In instances where the sequenced reads are shorter than the recommended read-length (e.g. 75bp for PET ChIP) the entire read length available is used.
    - WGBS:
        - No trimming will be performed on WGBS fastqs; if the phasing base has been included in the submitted fastqs, then it will be used in the analysis. For ENCODE data, however, we did trim the phasing base.
        - If an experiment has multiple read lengths, all runs will be processed without trimming.
    - RNA-Seq:
        - RNA-Seq fastq files will be trimmed to 75bp prior to analysis.
- EpiRRs with both SET and PET data:
    - For ChIP ANALYSIS objects where the antibody fastqs and control fastqs are not the same sequence type (ie. one is single-end (SET) and the other is paired-end (PET), only read 1 fastq(s) are used from the PET data and the container is run in single-end mode.
    - In cases where there is both SET and PET data for assay and/or control (i.e. at least one of either assay or control have both SET and PET fastqs):
        - If either assay or control only has SET data, the chip analysis is run in SET mode, using all SET fastqs and one fastq (mate 1) from each PET fastq pair (if PET data is available for the corresponding assay/control).
        - In cases where there is some PET data for both assay or control, in addition to SET data:
            - If the read counts in the PET fastqs meet threshold requirements, only PET data is used and the analysis is run in PET mode (note thresholds are pre-alignment).
            - If the thresholds are not met then we use all SET fastqs and mate 1 from PET fastqs pairs; the chip analysis done in SET mode.
    - For WGBS analysis objects with both SE and PE data, the container is run in the stranded (PBAL) configuration.

#### Dealing with mixed read lengths in ChIP-Seq data

This section describes the logic used to select readlength to pair control and assays to account for the fact that there may be more than one readlength present in experiments for the same reference epigenome.

- 75 and 74 bp readlengths are collected together under "74+75" readlength group, all others readlengths are in their own readlength group. 75PET is "75,75"
- Readcounts are computed across readlength groups
- Only readlength groups for which readcounts pass the minimum thresholds are kept per epirr/experiment as acceptable
    - If in case some epirr/experiment has no readlength group passing minimum thresholds then the deepest one is kept, so there's always one readlength group that is acceptable.
- For assay/control pairs where at least one of assay and control have more than one readlength group in the acceptable list, the assay and control readlength groups are selected as follows (in the order of priority):
    - If both (assay/control) have 75bp PET as acceptable, then that is chosen for analysis
    - If both (assay/control) have 75bp SET as acceptable, then that is chosen for analysis
    - If both (assay/control) have 75bp SET or PET as acceptable, then that is chosen for analysis (i.e. one has PET other SET, but of readlength 75).
      (All this avoids 75bp vs 32bp analysis if 32bp readlength group is the deepest, but sufficiently deep 75bp data is also available)
        - In case 75bp SET is chosen for analysis, then the first read from any 75bp PET data is also included.
    - Otherwise, the deepest assay and controls that are acceptable are chosen.
      This means we can still end up with 75bp vs 32bp (assay vs control) analysis if control has 75bp readlength, but fails to be deep enough to be acceptable.

#### Redundant Input Analysis

- Input redundancy across ChIP ANALYSIS objects:
    - The ENCODE pipeline which acts as the foundation for the IHEC ChIP container is configured to align and analyze an input control with each chromatin mark analyzed, thus producing up to 6 input bam files for each EpiRR.
    - In these cases, one input control bam file is randomly selected and disseminated for each EpiRR. The input control will be identified with the UUID for the selected container run and marked as *.ctl.bam
    - For the GIS project, one of the following three runs was used as controls for all 286 ChIP analyses corresponding to different EpiRR ids.
        - EGAR00001483412,EGAR00001483413,EGAR00001483414

In this case, only one analysis for these three EGAR ids has been retained across the 286. This means for 283 EpiRRs the control analysis files are present under a different EpiRR; this can be looked up using EGAR ids for control present in the metadata file.

##### Dealing with mixed readlengths in RNA-Seq data

- In the case where there are multiple files of various readlengths:
  - If one of the files is 75bp and has sufficient sequence coverage this file will be used for analysis.
  - In the instance that there is not sufficient coverage, the files will be merged and trimmed to the shortest readlength.

#### ChIP File Types For Dissemination

- .pval.signal.bigwig (not for input)
- .fc.signal.bigwig (not for input)
- .pval0.01.500K.narrowPeak.gz (not for input)
- .pval0.01.500K.bfilt.narrowPeak.gz (not for input)
- .noseq.bam: sequence stripped bam (for each histone modification and one file per EpiRR ID for the DNA Input control: .ctl_noseq.bam)
  - This is a bam file in which the sequence is replaced by a single N, qualities by a single # and the cigar string by 1M. This is done to avoid sharing genotype information while being able to use standard tools to work with the data. The code to do this is available at:
    - https://github.com/IHEC/integrative_analysis_chip/blob/dev-organize-output/encode-wrapper/bamstrip/bamstrip
    - https://github.com/IHEC/integrative_analysis_chip/blob/dev-organize-output/encode-wrapper/bamstrip/bamstrip.scala

- Raw bigwig (for each histone modification and one file per EpiRR ID for the DNA INput control: .ctl_raw.bigwig)
  - Since MACS bigwig tracks are normalized, and while the raw coverage tracks can be generated from stripped bam, we suggest making the following bigwig raw signal tracks available:
    - bigwig file using bamCoverage [bamCoverage 2.5.4, https://deeptools.readthedocs.io/en/2.5.4/content/tools/bamCoverage.html?highlight=bamcoverage ] available on ChIP container using command line: bamCoverage --extendReads -b $bam -o $bigwig -of bigwig --binSize 1 --maxFragmentLength 1000 -p 4
      - For SET, we propose using the extension length as the fragment length reported by the pipeline.
- .qc.json
- .qc.html

- Example:
  - ./call-macs2/shard-0/execution/ChIP-Seq.IX1239-A26688-GGCTAC.134224.D2B0LACXX.2.1.merged.nodup_x_ctl_for_rep1.pval.signal.bigwig
  - ./call-macs2/shard-0/execution/ChIP-Seq.IX1239-A26688-GGCTAC.134224.D2B0LACXX.2.1.merged.nodup_x_ctl_for_rep1.fc.signal.bigwig
  - ./call-macs2/shard-0/execution/ChIP-Seq.IX1239-A26688-GGCTAC.134224.D2B0LACXX.2.1.merged.nodup_x_ctl_for_rep1.pval0.01.500K.narrowPeak.gz
  - ./call-macs2/shard-0/execution/ChIP-Seq.IX1239-A26688-GGCTAC.134224.D2B0LACXX.2.1.merged.nodup_x_ctl_for_rep1.pval0.01.500K.bfilt.narrowPeak.gz
    Urls:
    http://www.epigenomes.ca/data/CEMT/resources/integrative_analysis/chip_example/ChIP-Seq.IX1239-A28471-ATCACG.134224.D2B0LACXX.2.1.merged.nodup_x_ctl_for_rep1.fc.signal.bigwig
    http://www.epigenomes.ca/data/CEMT/resources/integrative_analysis/chip_example/ChIP-Seq.IX1239-A28471-ATCACG.134224.D2B0LACXX.2.1.merged.nodup_x_ctl_for_rep1.pval0.01.500K.bfilt.narrowPeak.gz
    http://www.epigenomes.ca/data/CEMT/resources/integrative_analysis/chip_example/ChIP-Seq.IX1239-A28471-ATCACG.134224.D2B0LACXX.2.1.merged.nodup_x_ctl_for_rep1.pval0.01.500K.narrowPeak.gz
    http://www.epigenomes.ca/data/CEMT/resources/integrative_analysis/chip_example/ChIP-Seq.IX1239-A28471-ATCACG.134224.D2B0LACXX.2.1.merged.nodup_x_ctl_for_rep1.pval.signal.bigwig

#### WGBS File Types For Dissemination

- gemBS_reports.tar.gz - quality report, multiple files from GemBS
- .chg.bed.gz
- .chh.bed.gz

- ➔ .cpg.bed.gz
- ➔ .gembs_pos.bw
- ➔ .gembs_neg.bw
- ➔ Methylation files provided in ENCODE BED9+5 bedMethyl format:

| *Field* | *Description* |
|---|---|
| 1 | Contig or chromosome name. |
| 2 | Start position (0 offset) |
| 3 | End position (1 offset) |
| 4 | Name of item |
| 5 | Score from 0 - 1000 (capped methylation informative coverage) |
| 6 | Strand: +, - or ? |
| 7 | Start of where display should be thick |
| 8 | End of where display should be thick |
| 9 | Color value (RGB) |
| 10 | Methylation informative coverage |
| 11 | Percentage of reads showing methylation |
| 12 | Reference genotype |
| 13 | Sample genotype |
| 14 | Quality score for genotype call |

#### RNA File Types For Dissemination

All output of the 'quantification' step from grape-nf pipeline:

- ➔ .genes.results
- ➔ .isoforms.results
- ➔ .report.pdf
- ➔ .cnt
- ➔ .theta
- ➔ .model

The ‘coverage’ bigwig files
In case of ‘stranded’ RNA-Seq analysis 4 files

- ➔ .Unique.minusRaw.bw
- ➔ .Unique.plusRaw.bw
- ➔ .UniqueMultiple.minusRaw.bw
- ➔ .UniqueMultiple.plusRaw.bw

or 2 files for ‘strand-agnostic’ analysis:

- ➔ .UniqueMultiple.raw.bw
- ➔ .Unique.raw.bw

The output of the 'bamStats' process

- ➔ .stats.json

#### Data/Metadata Update Notes

The following analyses have been retracted:
September 14, 2021:

- ➔ aba54d95-4689-4915-917b-945c5f4002e5 for IHECRE00001921.1/H3K27ac
- ➔ c038c344-013d-4fc1-bae3-72a2c04e0f5f for IHECRE00001865/RNA-Seq
- ➔ ChIP Control analysis for following uuid’s removed as it was duplicated:

November 29, 2021:

- ➔ b870aaaf-89bd-41fc-a25d-012449f36024
- ➔ c1a9f844-6eda-4dd7-8282-c3d553cbcb4f

- 5571d7d7-4894-4e51-8082-c8a874cb2438
- 25743ca3-8e5d-4376-9b83-277f75741469
- de9f2c4e-6cfb-40a7-9b40-4fa537815a39
- 2c1739a8-c0c4-4f1f-9e6b-46c1ce8ff180
- e5395343-6b49-4355-a168-0f9b6320ac6f
- 088d52ff-571b-447d-8cd1-62f5248229a2
- c74b9473-7abb-43f6-8885-3c40e0552ac6
- 461b697a-4c20-47a7-8de2-3fc5af949e79
- 4c807da6-ffa8-4451-adb2-428243de39ad
- 313549b4-5e36-4d5c-9816-17b9c7f898e3
- dbaae77e-d923-4bd2-91ca-d217d8a52f61
- 517b2333-37c5-4b4b-b65c-1f93067b93ca
- 622e81e8-ba3b-4b86-bd94-afc1dae18757
- 6ca4dc93-02be-42e3-b833-5fbb7e12c934

April 01, 2022:

- 37 DEEP WGBS were removed as they were re-analyzed; UUIDs:
  - edc9d8dc-c2da-4060-b44d-c179b4fbb1a6
  - bba9e10a-f122-4b3a-8df0-6d2225820321
  - e209d428-7657-483f-89c4-558a6e95978c
  - efea66f6-e2f0-46a5-91c5-6d238e37bec6
  - 12561dc0-3b88-4454-b875-66b7af22b438
  - e415f4e5-80a1-40f4-8612-75956f5fba8f
  - 02ed1418-a901-4c9b-b419-81102121d302
  - b0156715-9be2-4cf3-a932-461a5ae3cefc
  - dedeb346-341e-4256-aea7-e41557ee1f5c
  - e6eb15cc-698f-42b6-8f74-22a2356b4051
  - 1d0590e7-206f-4fee-b841-67f5511f269f
  - a93097c4-9621-4e68-aed9-171b31a21a0b
  - 86794e0c-6dba-4281-88cc-bb49d3d2aee7
  - 7cdce640-7336-4997-9df8-63d232e82967
  - 8c318fe3-7e6f-4ad3-b081-33e19041eac0
  - a733133b-1435-4c64-b8cf-c52465a8ca17
  - e63867fb-e230-4175-947c-2be262173828
  - 27b754ea-a36a-48a3-b26f-465b7e843e4f
  - 2ac707d5-6902-460b-ab47-d1a1014395cf
  - 06f2b5a8-ea9c-46d2-8f8d-3a84d4ad1372
  - 6c71d8c6-5b6f-4246-99e5-b754555a40a5
  - d19d323e-1b45-4d70-8e1c-740f9f5a161d
  - 0bb3d15d-fb3b-4a8e-a76d-b5a0323bdb98
  - 2e3f584a-99ca-46da-9b16-74e3e01ee489
  - 3532f909-1de1-44d1-8311-e54aac541f5e
  - 39f9e2a8-2db0-4704-8bb4-c56faf4bf2ef
  - 4455ed91-d174-4800-ba54-2173ad849177
  - 5297aa01-65e2-4db1-a18e-983fb42457f8
  - 705716e2-d0ce-4e51-8723-7bef8b5e0926
  - 8a7e893c-ebf9-4990-a919-1b31c9d620ae
  - 95f55cc3-ac96-4425-b742-12d3c4cbf898
  - 99685d4f-8e18-475b-ada0-902adabf78d1
  - b1632df8-af62-44cb-bc9d-9a99b5af3703
  - cc0fb6d0-ecaa-45ee-a44d-bd7f2bb4dfb1
  - d655fa0c-1d91-402c-8054-d002cebef4eb
  - ec4694e4-e105-4f7f-91cb-64d42a69544e
  - fa86c062-62d3-4dbb-ab05-0b25a23a7690

May 25, 2022:

- All files (110 total) for the following 5 RNA-Seq objects were removed from incoming/RNA-Seq/03-18-2022 as these are non-human species:
  - IHECRE00000673.1 ef9d1c9e-4a31-4400-b170-3a3caf3cc923
  - IHECRE00000735.1 b0c27534-73b6-458b-8241-fa2df012d321
  - IHECRE00000755.1 ab834ff8-aee9-434a-b1ed-e881cdfa90bf
  - IHECRE00000812.1 db23f1db-e76f-4510-8267-8e8d04a90a15
  - IHECRE00000892.1 4c52eff9-5217-4c10-b6ac-9284ea5643c7
- ChIP-Seq metadata file incoming/readme/ihec_metadata.csv is updated for 1 entry (7a6a3fc8-9cb2-4f15-b9eb-2dd3707b3b59) :

- ◆ 'paired_end_mode' field for this entry is corrected 'true' → false
- ➔ A folder for special ChIP-Seq cases: when for one EpiRR ID IHECRE… there is a mixture of both pair-end and single-end objects. Analysis for PET objects always uses PET Input and SET objects SET input (read 1). In **incoming/ChIP-Seq/special_cases** we added control files (*.ctl_noseq.bam *.ctl_raw.bigwig) for the outlier objects matching the 'pair_end_mode' of the object.

September 13, 2022:

- ➔ 40 ENCODE ChIP-Seq analysis objects were removed from 'incoming/ChIP-Seq/03-10-2021/' (there were some omissions in the old processing).
  - ◆ 0649a668-6eaf-4952-87b7-38389b5571de IHECRE00004693.3:H3K4me3
  - ◆ 0e7ba49c-d144-46f9-9142-e2df62454489 IHECRE00004639.3:H3K4me3
  - ◆ 1946bf6f-f88d-480b-8508-3fc5d6e3d2cd IHECRE00001889.4:H3K4me1
  - ◆ 1ae5443f-9a13-401f-a7ab-7704d265e082 IHECRE00004653.3:H3K9me3
  - ◆ 1fa24192-b43a-452e-ac02-f8802707e2e5 IHECRE00001889.4:H3K27me3
  - ◆ 23405835-67d8-4d62-94d2-8e2a0c509bfa IHECRE00001853.4:H3K27ac
  - ◆ 2a109be7-9a2d-47eb-b953-9130ee7e1ac1 IHECRE00004653.3:H3K36me3
  - ◆ 302909ad-6dc8-44d2-87df-62820d3c994f IHECRE00003714.4:H3K36me3
  - ◆ 329f660b-f08f-46ff-8cd2-fb5d7b32726b IHECRE00003705.4:H3K27ac
  - ◆ 36b3686f-70b6-465f-bd81-1a30942b400a IHECRE00003709.4:H3K27ac
  - ◆ 37d619a3-e813-4171-8059-de82e05c9722 IHECRE00003705.4:H3K36me3
  - ◆ 3d13277e-62d9-4648-aa99-14f657578951 IHECRE00004653.3:H3K4me1
  - ◆ 3e78d219-bac4-4fdb-bad5-d6204841b86d IHECRE00001893.4:H3K4me1
  - ◆ 40913545-ead8-4b5c-8dd4-6279977b3e89 IHECRE00001887.4:H3K27me3
  - ◆ 45ef4425-6b9e-4657-8fcc-9b246b5d467f IHECRE00001853.4:H3K36me3
  - ◆ 49649872-771a-43a8-8839-c967f7ee4c12 IHECRE00001887.4:H3K36me3
  - ◆ 71688e84-128f-483c-aedb-7bfd34d33421 IHECRE00003705.4:H3K27me3
  - ◆ 72171428-26b5-4e6b-9e20-7ac7b0b9e437 IHECRE00004653.3:H3K27ac
  - ◆ 76912096-e086-4d35-9a38-b78d3b1335ab IHECRE00001853.4:H3K27me3
  - ◆ 78a982a4-f81a-4bba-abbf-78766d160906 IHECRE00001889.4:H3K27ac
  - ◆ 82e10062-512a-4347-9149-645e70874010 IHECRE00001887.4:H3K4me3
  - ◆ 86432d64-2f82-4499-b348-759c14d68d04 IHECRE00001887.4:H3K4me1
  - ◆ 8f314ecf-34a0-43cb-8b0f-e5c775d83a1a IHECRE00001887.4:H3K9me3
  - ◆ 97f5c762-ec63-486f-896e-be217697fa95 IHECRE00003705.4:H3K9me3
  - ◆ 98744231-f8dc-47dc-a676-8277daa2e9e5 IHECRE00001893.4:H3K9me3
  - ◆ 9aab601e-bf13-4104-ade2-d381d02f561d IHECRE00004658.3:H3K4me3
  - ◆ a6f7dca3-2de6-4ea4-8c6a-ab4e283df59f IHECRE00004658.3:H3K27me3
  - ◆ a88248a8-71df-411b-a718-80fb6d4dd0d5 IHECRE00003705.4:H3K4me3
  - ◆ adcfea8c-9a81-4b7c-a2ee-7288e6f0ae48 IHECRE00001893.4:H3K27ac
  - ◆ b4e5e69c-f1d6-41b8-9299-2e4f03ab49be IHECRE00001893.4:H3K36me3
  - ◆ c2b1aef1-49e7-455b-a074-221e9950bd33 IHECRE00001887.4:H3K27ac
  - ◆ c6f5372a-c77d-426c-801c-dfa5e2589606 IHECRE00003709.4:H3K4me3
  - ◆ d656aac8-069c-4ff2-aa29-d33f1d294996 IHECRE00003709.4:H3K36me3
  - ◆ d91cce00-3b22-47f8-aa5b-acfbdf73c3a2 IHECRE00001889.4:H3K4me3
  - ◆ e82fdcde-d9a9-4156-97dd-ba6eadba706a IHECRE00001893.4:H3K27me3
  - ◆ ecd1ad75-cc2d-445e-9f4f-257fe6ae93b1 IHECRE00001889.4:H3K9me3
  - ◆ ece99048-e9cb-4403-b4de-396dd286b601 IHECRE00004639.3:H3K4me1
  - ◆ f00c29fd-1661-4d8a-9e74-d995607022a2 IHECRE00003709.4:H3K4me1
  - ◆ f141e979-777d-44d3-bc88-4cb297c1093b IHECRE00001889.4:H3K36me3
  - ◆ f788e248-6847-4a76-95a2-c2cdaabd8ec8 IHECRE00003714.4:H3K9me3
- ➔ Most (all, but 3) of those objects were replaced with new re-analyzed objects and can be found in
- ➔ incoming/ChIP-Seq/09-11-2022/
- ➔ The outstanding 3 will be posted together with the rest ~20 ENCODE ChIP-Seq objects.

October 18, 2022:

- ➔ a number of redundant *.ctl_noseq.bam and *.ctl_raw.bigwig (as well as the corresponding md5 sum files) were removed from the SFTP. Those correspond to the cases when the same DNA Input control was used to process multiple 'Histone mod' analysis objects.
- ➔ ChIP-Seq metadata file : ihec_metada.csv file was cleaned (in a number of lines the control file ID and read length was recorded twice; now all entries are harmonized and duplications are removed).

October 25, 2022:

- ➔ The following 7 'non-canonical' objects are removed from the SFTP.
  - ◆ 436978ce-b636-4480-a0a0-f834297d6953 IHECRE00000016.3 H3K9/14ac
  - ◆ 45a3cb71-c6fa-4373-8607-e320ee8c7fbd IHECRE00000058.3 H3K9/14ac
  - ◆ 7c97afdb-41f0-4cee-b769-24774a553443 IHECRE00000101.3 H3K9/14ac

- 96262a89-5f4f-45c8-8c9d-4153c57db9ba IHECRE00001516.1 H3K9/14ac
- 593c85eb-6bba-46c7-8938-f2db6a00acfc IHECRE00000094.3 H2A.Zac
- 8bbbb97c-d89a-47d6-b245-90c4424a4356 IHECRE00001909.1 reH3K4me3
- fe115099-c04c-44b4-a4ee-797164e5dc86 IHECRE00001909.1 reH3K27me3

December 7, 2022:
Revoking and partly revoking previously posted ChIP-Seq analysis objects

➔ 21 duplicated/posted NIH Roadmap ChIP-Seq objects were retracted:
  - 1ec4db60-59cd-40d5-b56a-125c146d436c,IHECRE00000957.7,H3K27me3
  - 8715f4a9-62f6-4092-8d5c-ad287b772798,IHECRE00000957.7,H3K36me3
  - 665ea296-d46d-4d46-90d3-b6be949766a5,IHECRE00000980.8,H3K36me3
  - 19c6e2d6-2297-47f1-b931-4cae2f516aa7,IHECRE00000980.8,H3K4me1
  - fba37eb9-ecde-46ba-9efc-e1865f63715c,IHECRE00000985.7,H3K27ac
  - f2ec28f9-8751-4051-a4fe-d5aac77cd037,IHECRE00000985.7,H3K36me3
  - 6ef9a144-279c-4f5d-87d1-e557a3d88488,IHECRE00000985.7,H3K4me3
  - 5383890b-a46b-457f-aae1-45b409439b73,IHECRE00000985.7,H3K9me3
  - b9506ce5-a99c-48a6-9a22-b7f58ec3ed71,IHECRE00000992.7,H3K27me3
  - a81cc705-d717-4774-a11d-77760afc1bc0,IHECRE00000992.7,H3K9me3
  - e837e45f-a6be-401d-8207-b0a3a1f64ce6,IHECRE00001025.7,H3K36me3
  - fa67a27a-3515-43ee-9984-f45b4625ea6f,IHECRE00001025.7,H3K4me1
  - 7059117a-04ca-40a1-ab0b-3a8e9db2e9d3,IHECRE00001025.7,H3K9me3
  - 0f8ee4ba-af78-48a4-8b53-3a2fc4d9455a,IHECRE00001027.7,H3K27ac
  - b50ae014-ba49-4e74-bbd2-f6f724facedd,IHECRE00001027.7,H3K4me1
  - 0c1061c3-b47e-4623-ac31-003a4d78a375,IHECRE00001050.8,H3K27ac
  - af35967a-3d9e-4468-bb4f-9422940f5e39,IHECRE00001050.8,H3K27me3
  - fdf1caf5-a901-40e5-899d-c3bb6ccdfd78,IHECRE00001050.8,H3K36me3
  - 9c911e6b-1e99-4947-8fbb-7e825a8b310f,IHECRE00001050.8,H3K4me1
  - 808aa963-69a8-4d1a-acfa-76ba99955376,IHECRE00001050.8,H3K4me3
  - a28f4062-44b2-458c-a8ba-d906f8f9460c,IHECRE00001050.8,H3K9me3

➔ 7 ChIP-Seq objects were revoked
  - ➔ 0b2f031a-c90a-4a28-b291-a1dddcf91924,IHECRE00003403.1,H3K36me3
  - ➔ 3ba69f70-1250-4c19-aabf-beb85e72ffba,IHECRE00003399.1,H3K36me3
  - ➔ 4db25a37-8089-4580-95e5-5f391fc9b3f4,IHECRE00001897.1,H3K27me3
  - ➔ 502793f2-8bf9-4ed5-8925-cb68c1390393,IHECRE00001896.1,H3K27me3
  - ➔ 7b42a3e3-8f3c-4d44-baf4-460cb386de2d,IHECRE00003399.1,H3K9me3
  - ➔ b9d09b38-47e0-4fc0-878f-f81351b475f8,IHECRE00001053.7,H3K4me1
  - ➔ d0f06464-403b-42d2-8352-74af2794534f,IHECRE00003403.1,H3K9me3

➔ 14 ChIP-Seq objects partly revoked
  - 2dbe488e-1c42-49eb-9690-80090a9b19c1,IHECRE00000928.7,H3K9me3
  - 926e770c-0dff-4937-b642-529da65a20bd,IHECRE00000928.7,H3K4me1
  - c7b87805-6a4a-46a8-8e4b-5256f02641c6,IHECRE00000928.7,H3K4me3
  - d24b2884-99cd-4ee9-8b21-696930091915,IHECRE00000928.7,H3K27me3
  - da34a9ed-d7c6-4abe-8466-f69230cd8874,IHECRE00000928.7,H3K27ac
  - e6582d8d-bc61-47ec-9dff-63d198ba121d,IHECRE00000928.7,H3K36me3
  - 82201785-c340-4021-8723-c3568095060c,IHECRE00000998.7,H3K4me1
  - bb87de8a-2776-4cdd-b51a-53685dbf9503,IHECRE00000998.7,H3K9me3
  - bed55195-70ef-4efc-bf5c-ac16b3c316ad,IHECRE00000664.3,H3K4me3
  - ed7d4e5c-c3c3-4cdb-af6a-6db10a11a3cb,IHECRE00000664.3,H3K9me3
  - 4966aa29-638b-4cd4-ac8b-845df7ec532c,IHECRE00000664.3,H3K27me3
  - 7a96b85e-edf5-4620-8bd9-9bb72841ba7e,IHECRE00000664.3,H3K36me3
  - 27c8da51-0b57-409b-a30e-95c9bc040cc0,IHECRE00000664.3,H3K27ac
  - 807ac7f6-351d-4e41-ab0a-54ea6e0d397d,IHECRE00000664.3,H3K4me1

alignment files for ChIP-Seq data can be found in incoming/ChIP-Seq/12-06-2022/

#### Data Posting Dates

December 09, 2022

**Please note that for the data freeze we combined all 3 metadata files into one file and harmonized the fields. The path to the Analysis object files was also added into the same file incoming/readme/epiatlas_metadata.csv**

November 26, 2022:

- REMC ChIP-Seq
  - incoming/ChIP-Seq/11-18-2022/ 182 objects (+40 controls)
  - incoming/ChIP-Seq/11-21-2022/ 113 objects (+21 controls)

October 25, 2022:
- 7 ChIP-Seq objects for the antibodies/marks not included in the analysis are removed (see Notes above)

October 05, 2022 - updated:
- incoming/readme/ihec_metadata_wgbs.csv - fixed protocol PBAT/standard for all objects
- incoming/readme/ihec_metadata_rna.csv - fixed protocol mRNA-Seq/total-RNA-Seq for all objects

October 03, 2022:
- incoming/WGBS/10-03-2022/ (4 stragglers moved)

September 20, 2022:
- McGill [EMC] ChIP-Seq
  - incoming/ChIP-Seq/09-15-2022/ (1 ‘straggler’ object added: 26047187-2aea-47ae-97b4-1842ebc14e59)
- BLUEPRINT DNA Methylation
  - incoming/WGBS/09-20-2022/ (28 stragglers WGBS-standard objects)
- DEEP DNA Methylation
  - incoming/WGBS/09-20-2022/ (4 stragglers PBAT objects)

September 19, 2022:
- ENCODE ChIP-Seq
  - incoming/ChIP-Seq/09-18-2022/ (20 objects added)
    NOTE corresponding controls for all but 1 IHEC Id can be found in other ENCODE posting folders:
    incoming/ChIP-Seq/03-10-2021/
    incoming/ChIP-Seq/09-11-2022/

September 18, 2022:
- REMC RNA-Seq
  - incoming/RNA-Seq/09-18-2022/ (26 mRNA objects)
- REMC WGBS
  - incoming/WGBS/09-18-2022/ (25 objects)
- McGill [EMC] ChIP-Seq
  - incoming/ChIP-Seq/09-15-2022/ (38 objects added)
    NOTE1: One struggler object has to be worked on
    NOTE2: The Inputs (10 different cell types) - were not registered at the EpiRR; as a result we could not record any sequence archive file ID for the DNA Inputs.

September 13, 2022:
- ENCODE ChIP-Seq
  - incoming/ChIP-Seq/09-11-2022/ (95 objects added)
  - incoming/ChIP-Seq/03-10-2021/ (40 objects removed, see above ‘Data Update Notes’ section)

  NOTE if ‘ctl’ files for some objects in ‘incoming/ChIP-Seq/09-11-2022/’ are missing they can be found in ‘incoming/ChIP-Seq/03-10-2021/’. For IHECRE00001893.4 we also added a 75bp control (for H3K4me3 mark) into incoming/ChIP-Seq/special_cases; the rest of the marks for the EpiRR are 36bp analysis.

August 31, 2022:
- BLUEPRINT 8 ChIP-Seq ‘strugglers’ objects
  - incoming/ChIP-Seq/08-30-2022/ (8 objects)

  NOTE: files for controls for this ChIP-Seq objects (*.ctl_noseq.bam *.ctl_raw.bigwig) can be found in incoming/ChIP-Seq/08-24-2022/ or incoming/ChIP-Seq/08-25-2022/

August 28, 2022:
- BLUEPRINT 1019 ChIP-Seq objects
  - incoming/ChIP-Seq/08-24-2022/ (499 objects)
  - incoming/ChIP-Seq/08-25-2022/ (520 objects)

August 18, 2022:
- one BLUEPRINT RNA-Seq object from incoming/RNA-Seq/05-06-2022/ removed due to truncated isoform file (IHECRE00000053.3.f6335dc5-9347-4ef4-ae4d-c7b7fbb5f565)

August 17, 2022:

- ➔ BLUEPRINT 675 RNA-Seq objects: incoming/RNA-Seq/08-17-2022/

August 08, 2022:
- ➔ Roadmap 383 ChIP-Seq objects: incoming/ChIP-Seq/07-21-2022/
  NOTE: one 'struggler' was added on August 12.

July 09, 2022:
- ➔ BLUEPRINT 52 WGBS objects: incoming/WGBS/07-07-2022/
- ➔ DEEP 33 WGBS objects (reprocessed): incoming/WGBS/07-09-2022/

May 25, 2022;
- ➔ Removed 5 McGill RNA-Seq Non-human objects (see above)

May 16, 2022:
- ➔ ENCODE 99 RNA-Seq objects: incoming/RNA-Seq/05-16-2022/

May 09, 2022:
- ➔ CEMT 43 WGBS objects: incoming/WGBS/05-09-2022
- ➔ BLUEPRINT 191 RNA-Seq objects: incoming/RNA-Seq/05-06-2022/

May 03, 2022:
- ➔ CEMT 6 ChIP-seq objects: incoming/ChIP-Seq/05-03-2022
- ➔ CEMT 21 RNA-seq objects: incoming/RNA-Seq/05-02-2022

April 26, 2022:
- ➔ CEMT 6 ChIP-Seq objects: incoming/ChIP-Seq/04-26-2022/ (note ctrl bam/bigwig files can be found in: incoming/ChIP-Seq/03-16-2022)

April 25, 2022:
- ➔ ChIP-Seq: 652+247 analysis objects
  - ◆ metadata : incoming/readme/ihec_metadata.csv (currently 3691 objects)
  - ◆ data: incoming/ChIP-Seq/04-21-2022/ (652 CEMT ChIP-seq objects) - please note some ctrl bam/bigwig files can be found in: incoming/ChIP-Seq/03-16-2022)
  - ◆ data: incoming/ChIP-Seq/04-22-2022/ (247 BLUEPRINT analysis objects)

April 05, 2022:
- ➔ RNA-Seq: 1 DEEP dataset (IHECRE00003819.1)
  - ◆ updated metadata: incoming/readme/ihec_metadata_rna.csv (613 objects)
  - ◆ data path: incoming/RNA-Seq/04-05-2022
- ➔ WGBS: 1 ENCODE dataset (IHECRE00001851.4)
  - ◆ updated metadata: incoming/readme/ihec_metadata_wgbs.csv (460 objects)
  - ◆ data path: incoming/WGBS/04-05-2022

April 01, 2022:
- ➔ ChIP-Seq: DEEP (9) analysis objects
  - ◆ metadata : incoming/readme/ihec_metadata.csv (currently 2792 objects)
  - ◆ data: incoming/ChIP-Seq/04-01-2022/
  - ◆ Note: that corresponding 'ctl_noseq' files for the corresponding EpiRRs were posted previously
- ➔ RNA-Seq: REMC (NIH Road Map) 35 data sets
  - ◆ metadata: incoming/readme/ihec_metadata_rna.csv (currently 612 objects)
  - ◆ data: incoming/RNA-Seq/04-01-2022/
- ➔ WGBS: 37 DEEP datasets were removed from SFTP site (as they are being reanalyzed). UUID are posted in the notes (above)
  - ◆ metadata is updated: incoming/readme/ihec_metadata_wgbs.csv (currently 459 objects)

March 23, 2022:
- ➔ ChIP-Seq: EpiHK(6) + CEMT(113)
  - ◆ metadata : incoming/readme/ihec_metadata.csv
  - ◆ (currently 2783 objects)
  - ◆ data:
    - ● EpiHK(6): incoming/ChIP-Seq/03-15-2022/
    - ● CEMT(113): incoming/ChIP-Seq/03-16-2022/
- ➔ RNA-Seq: DEEP(85) + McGill(338) + EpiHK(1)
  - ◆ metadata: incoming/readme/ihec_metadata_rna.csv
  - ◆ currently 577 RNA-Seq datasets: 107 CEMT, 33 CREST, 85 DEEP, 1 EpiHK, 13 KNIH, 338 MCGILL_EMC

- Note some EpiRR IDs have several RNA-Seq data (mRNA and total RNA-Seq). The information on the RNA-Seq experiment type is recorded in the metadata file.
- ➔ data:
  - DEEP(85) + EpiHK(1): incoming/RNA-Seq/02-17-2022/
  - McGill(338): incoming/RNA-Seq/03-18-2022/

February 14, 2022:
- ➔ 15 Deep2 WGBS libraries
  - Path to updated metadata: incoming/readme/ihec_metadata_wgbs.csv
    - Path to the data: incoming/WGBS/04-02-2022

February 01, 2022:
- ➔ 9 "stragglers" [6 CREST, 2 McGill, 1 GIS ChIP-Seq analysis]
  - Path to updated metadata: incoming/readme/ihec_metadata.csv
    - Path to the data: incoming/ChIP-Seq/01-22-2022/

January 28, 2022:
- ➔ 90 Deep2 ChIP-Seq analysis
  - Path to updated metadata: incoming/readme/ihec_metadata.28012022.csv
    - Path to the data: incoming/ChIP-Seq/01-26-2022/

November 29, 2021:
- ➔ 31 Blueprint ChIP-Seq analysis + 107 CEMT RNA-Seq analysis
  - Path to updated metadata: incoming/readme/ihec_metadata.csv
    - Path to Blueprint ChIP-Seq data: incoming/ChIP-Seq/11-29-2021/
  - Path to updated metadata: incoming/readme/ihec_metadata_rna.csv
    - Path to CEMT RNA-Seq data: incoming/RNA-Seq/11-12-2021/

October 22, 2021:
- ➔ 33 CREST RNA-Seq analysis
  - Path to updated metadata: incoming/readme/ihec_metadata_rna.csv
  - Path to CREST RNA-Seq data: incoming/RNA-Seq/10-18-2021/
    - incoming/RNA-Seq/CREST_QC_summary_Oct222021.txt

September 14, 2021:
- ➔ 80 CEMT WGBS analysis + 1 EpiHK WGBS analysis
  - Path to updated metadata: incoming/readme/ihec_metadata_wgbs.csv
  - Path to WGBS data: incoming/WGBS/09-08-2021
  - Path to EpiHK and CEMT QC metrics
    - incoming/WGBS/EpiHK_QC_summary.txt
    - incoming/WGBS/CEMT_QC_summary_Sep142021.txt

August 17, 2021:
- ➔ 14 RNA-Seq analysis + 22 WGBS analysis
  - Path to updated metadata: incoming/readme/ihec_metadata_wgbs.csv
  - Path to WGBS data: incoming/WGBS/08-10-2021/
  - Path to updated metadata: incoming/readme/ihec_metadata_rna.csv
  - Path to RNA-Seq data: incoming/RNA-Seq/08-11-2021/
  - Path to QC metrics:
    - incoming/WGBS/REMC_QC_summary_dissemination.txt

June 15, 2021:
- ➔ 8 WGBS analysis
  - Path to updated metadata: incoming/readme/ihec_metadata_wgbs.csv
  - Path to QC metrics:
    - incoming/WGBS/REMC_PairedEnd_only_QC_summary_dissemination.txt
  - Path to data: incoming/WGBS/06-09-2021

May 18, 2021:
- ➔ 322 WGBS analysis
  - Path to updated metadata: incoming/readme/ihec_metadata_wgbs.csv
  - Path to QC metrics:
    - incoming/WGBS/211315_CREST_QC_summary_dissemination.txt
    - incoming/WGBS/211316_EMC_QC_summary_dissemination.txt
    - incoming/WGBS/211317_KNIH_QC_summary_dissemination.txt

    - incoming/WGBS/211643_Blueprint_QC_summary_dissemination.txt
  - Path to data: incoming/WGBS/05-07-2021

April 04, 2021:

- ➔ 214 ChIP CREST analysis
  - path to data: incoming/ChIP-Seq/04-06-2021
  - metadata file: incoming/readme/ihec_metadata.csv

March 17, 2021:

- ➔ 811 ChIP analysis made available on SFTP
  - path to data: incoming/ChIP-Seq/03-10-2021 (525 analysis for ENCODE)
  - path to data: incoming/ChIP-Seq/03-11-2021 (286 analysis for GIS)
  - metadata file: incoming/readme/ihec_metadata.csv
  - Please see note about GIS project in “Redundant Input Analysis” section

February 17, 2021:

- ➔ 3 WGBS analysis made available on SFTP
  - path to data: incoming/WGBS/02-03-2021
  - metadata file: incoming/readme/ihec_metadata_wgbs.csv
  - WGBS QC summary: incoming/WGBS/DATA-277_Jan27_2021_3_rerun_DEEP_samples_6_metrics.xlsx

January 19, 2021:

- ➔ 45 WGBS analysis made available on SFTP
  - path to data: incoming/WGBS/01-19-2021
  - metadata file: incoming/readme/ihec_metadata_wgbs.csv
  - WGBS QC summary: incoming/WGBS/WGBS_ENCODE_DEEP_6_QC_Metrics_Jan19_ac.xlsx

December 18, 2020:

- ➔ 1029 ChIP-Seq analysis made available on SFTP

September 14, 2020:

- ➔ 356 ChIP-Seq analysis made available on SFTP

November 17, 2020:

- ➔ 124 ChIP-Seq analysis made available on SFTP
